# A critical role for the anterior insula in impaired adult social interaction after early life maltreatment

**DOI:** 10.64898/2026.05.31.729133

**Authors:** Matthew G. Kearney, Zoe Beatty, Miranda Mayeaux, Ziyi (Zephyr) Wang, Oluwarotimi Folorunso, Celia Middleton, Emily L. Newman, William A. Carlezon, James McNally, Kerry J. Ressler, Erin E. Hisey

## Abstract

Maltreatment in children and adolescents poses a major public health crisis as a primary factor for the development of adult psychiatric disorders. However effective treatments for symptoms resulting from early life maltreatment are lacking, in part due to a lack of understanding of how neural circuits are disrupted in adulthood after childhood and adolescent abuse. We used a novel model of developmental maltreatment (early adolescent chronic social defeat stress, eaCSDS) in C57/B6J mice to examine behavioral and circuit-level effects in the adult anterior insula (AI). We combined chemogenetics, whole brain c-fos imaging and multi-site LFP to investigate the role of the AI and related circuitry in adult behavioral dysfunction after early life maltreatment. Behavioral analysis in adult animals reveals that males, but not females, show robust generalized social avoidance after social defeat in early adolescence. Chemogenetic silencing of AI neurons in males, but not females, reduces social avoidance in adulthood after eaCSDS. In males, whole brain c-fos imaging and multi-site LFP recordings further reveal circuit-level disruptions in connectivity to AI, implicating AI dysregulation as a driver of adult male social avoidance after adolescent maltreatment. Together these findings reveal AI as a novel circuits-level target whose activity normalization may reduce fear-based symptoms of adolescent maltreatment in adulthood specifically in males.

## INTRODUCTION

Childhood maltreatment is a key predictor of mental illness in adulthood^1–4^ and increases the risk of development of obesity, diabetes and heart disease^5–7^. It presents a pressing public health crisis as nearly 20% of children are affected across the globe, independent of race or nationality^8^. Animal models are needed in order to examine how maltreatment reshapes specific neural circuits and gene pathways to affect adult behavior. However, while some well-characterized rodent models of ***infant*** neglect and abuse alone or paired with adult physical trauma exist^9–12^, models that specifically target a window akin to ***childhood*** (i.e. in post weaning young animals before the onset of puberty) with a physical component that can robustly affect adult behavior are nearly absent. This is notable in that different forms of early life stress can have differential effects on the brain in both humans and rodent models; early life neglect has been seen to accelerate maturation of neural circuits while early life physical abuse has been seen to delay maturation^13,14^. Thus, we recently developed a model of childhood to early adolescent social defeat stress in mice to allow us to determine the neural circuit components most critically affected in adulthood after the exposure to early life maltreatment.

We wanted to examine the role of the anterior insula (AI) in the adult brain after early adolescent social defeat stress as AI structure and function are markedly disrupted across a number of psychiatric disorders that are comorbid with early life maltreatment in humans^15–17^. AI acts as a hub of salience processing in humans, allowing for switching between network states depending on the behavioral context^18,19^. Patients with anxiety and depressive disorders show decreases in salience network connectivity; these disruptions are strongly associated with childhood trauma^20^ and predictive of substance use disorders^21,22^. Furthermore, AI is known to regulate social approach and avoidance in animal models^23,24^. AI as a whole^25^ as well as its specific input to prefrontal cortex^26^ is necessary for social approach. Thus, we predicted AI may play a fundamental role in social interaction after early life maltreatment.

## RESULTS

### Early adolescent social defeat results in robust fear generalization in adulthood in social and conditioning paradigms in males

We first wanted to more comprehensively characterize the behaviors affected by our early adolescent chronic social defeat stress (eaCSDS) model^27^, which we modified from a traditional adult chronic social defeat stress paradigm^28,29^. The exposure to physical abuse between the age of 10 and 12 has been reported to result in the highest incidence of psychiatric disorders in adulthood^30^; thus we developed our model to target the end of the prepubescent phase through early pubescence in male and female mice. Briefly, we performed 10 day chronic social defeat stress on postnatal (P29 +/-3 days) C57/B6J mice by introducing the adolescents to the home cage of an adult CFW of the same sex for an aggressive encounter (Figure 1a, Supplemental Video 1). We subsequently housed the adolescent beside the CFW aggressor overnight, separating them with a perforated plexiglass partition that allowed olfactory and visual, but not physical, contact. We repeated this procedure for the next 9 days with a novel CFW aggressor each day.

**Figure 1.**
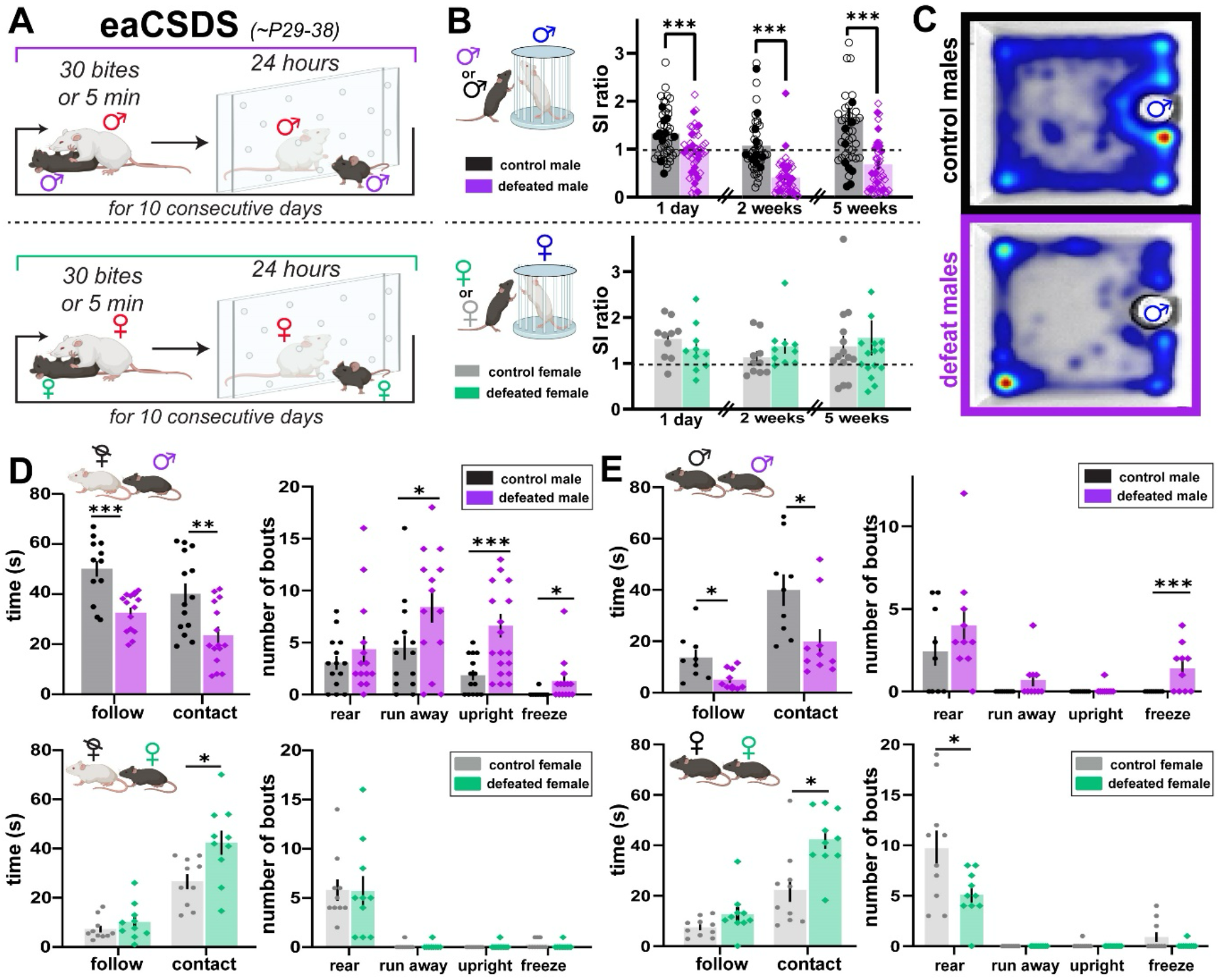
eaCSDS results in robust avoidance in adulthood in social interaction paradigms in males but not females. A) Schematic of early adolescent chronic social defeat stress paradigm (eaCSDS). Purple male symbol, defeated C57/B6 male. Red male symbol, aggressive CFW male. Green female symbol, defeated C57/B6 female. Red female symbol, aggressive CFW female. B) Social interaction ratios from OFSI with a non-aggressive CFW of the same sex for control male (black, n = 46), defeated male (purple, n = 50), control female (grey, n = 10 for 1 day and 2 weeks, n = 15 for 5 weeks post), and defeated female (green, n = 10 for 1 day and 2 weeks post, n = 15 for 5 weeks post) mice at multiple time points after defeat:1 day (unpaired two-tailed t-test, p = 0.0002), 2 weeks (unpaired two-tailed t-test, p < 0.0001; 2 defeated mice detected as outliers with ROUT (Q=1%) outlier test and removed), and 5 weeks (unpaired two-tailed t-test, p < 0.0001; 4 control and 2 defeated mice detected as outlier with ROUT (Q=1%) outlier test and removed) after the last day of eaCSDS. Social interaction ratio lower than 1 (denoted by dashed line) indicates socially avoidant behavior. For males, filled symbols are mice defeated beginning exactly at P29, non-filled symbols are mice defeated beginning at P29 +/-3 days. All females defeated beginning at P29 +/-3 days. C) Averaged heat maps during open field social interaction (OFSI) with non-aggressive adult CFW male (blue male symbol) from 10 control male mice (top) and 10 defeated male mice (bottom) in adulthood (P70). D) Behavior during 90 second home cage social interaction (HCSI) with ovariectomized adult CFW female (blue dashed female symbol) 10 days after defeat in males (top) and females (bottom). Behaviors quantified based on time spent performing behavior (left) or number of bouts (right) (unpaired two-tailed t-test between control (black, n = 14 males; grey, n = 10 females) and defeated (purple, n = 14 males; green, n = 10 females) mice for each behavior). *, p < 0.05; **, p < 0.01; ***, p < 0.001 for all figures shown. Error bars indicate standard error of the mean for all figures shown. E) As in D but with non-aggressive C57 animal of the same sex.

To determine the impact of eaCSDS on social behavior, we first examined social interaction at multiple timepoints across development using the open field social interaction test (OFSI)^31^. Example heat maps averaged across 10 mice in the presence of an unfamiliar nonaggressive CFW male (contained in a wire cup, white circle in heatmaps) show decreased time spent in the interaction zone in defeated mice (Figure 1b). In comparison to control male mice (black symbols), we found that male mice defeated beginning at exactly P29 (filled purple symbols) as well as at P29 +/-3 days (hollow purple symbols) had significantly lower social interaction ratios (defined as amount of time in interaction zone in presence of CFW / amount of time in interaction zone in absence of CFW) in early (1 day post defeat) and late (2 weeks post defeat) adolescence and young adulthood (5 weeks post defeat) (Figure 1b,c), with over 75% of defeated adult males showing susceptibility to defeat (susceptibility to defeat characterized as social interaction ratio < 1^12,32^). In contrast to males, defeated female animals showed no significant changes in social interaction ratio at any of the time points tested (1 day post defeat, 2 weeks post defeat, 5 weeks post defeat) (Figure 1b, green).

To examine if specific prosocial and defensive behaviors were altered after eaCSDS during more naturalistic social interaction we recorded 90 second interactions with an unfamiliar nonaggressive CFW female in the home cage of defeated and control mice 10 days after defeat (home cage social interaction, HCSI) (Supplemental Videos 2,3). Using a machine learning based approach^33,34^, we quantified the amount of time spent walking, following, or with nasal contact to the body of the CFW female, and the number of bouts of more rapid onset-offset behaviors including rearing, rapid movement away from the CFW female (‘run away’), upright submissive posturing (‘upright’) and freezing. Strikingly, we found that defeated male mice spent significantly less time engaging in prosocial behaviors such as following and making contact with the CFW female, but they performed significantly more bouts of defensive behaviors such as running away, upright submissive posturing and freezing (Figure 1d, top). Notably, we found that defeated females showed no significant differences in the number of defensive behaviors towards the CFW female and instead showed a significant increase in the time spent making contact with the CFW female (Figure 1d, bottom). Similar patterns of behavior held during home cage social interaction with C57 mice of the same sex and age as the defeated males and females; namely, defeated males showed decreased following and contact and increased freezing (Figure 1e, top) while defeated females showed increased contact and no change in defensive behaviors (Figure 1e, bottom) towards age matched same sex mice of the same strain. Thus, both more traditional (OFSI) and freely moving (HCSI) paradigms for social interaction revealed an increase in social avoidance in defeated males after the completion of eaCSDS. Interestingly, defeated females showed no difference in social interaction ratio in OFSI and increased, rather than decreased, social contact in HCSI, underscoring a notable difference between males and females in their social response after eaCSDS.

The long-term avoidance exhibited in defeated males towards unfamiliar, nonaggressive males, as well as females, of the same strain as the eaCSDS aggressors as well as to mice of their own strain (C57) may represent a translationally-relevant feature of our model; indeed, generalization of fear to innocuous stimuli that share features with dangerous ones is a hallmark of trauma-related disorders in humans^35–37^. To examine generalization in a more controlled setting in adult male mice with and without defeat history, we used classical fear conditioning to threat (CS+, paired with shock) and safety (CS-, not paired with shock) tones and examined freezing to conditioned tones and novel intermediate tones the following day (Supplemental Figure 1a). While freezing during training (day 1) to the CS+ and CS-was not significantly different between controls and defeated mice (Supplemental Figure 1b), we found a significant group difference in freezing across all tones during testing (day 2) between control and defeated (Supplemental Figure 1c) and that freezing to novel tones was significantly increased in defeated compared to controls (Supplemental Figure 1d). This increased generalization of fear to novel tones was complementary to our finding of increased generalization of fear to social partners, suggesting that increased generalization in adulthood is driven by trauma exposure in early life. The most striking behavioral phenotype we observed in adulthood after eaCSDS in males was that of generalization; defeated males did not display significantly different behavior from controls in a test of anxiety-like behavior (open field test, OFT, Supplemental Figure 2a,b) or in a test of simple working memory (Y-maze) (Supplemental Figure 2c,d,e). Additionally, we compared traditionally used stress biomarkers^38,39^ in defeated versus control male mice in adulthood; free cortisol levels after re-exposure to a non-familiar CFW male in adult male mice trended towards an increase in defeated compared to controls but were not significant (Supplemental Figure 3a). A non-significant increase in adrenal weight was also observed in defeated relative to control male mice (Supplemental Figure 3b) while thymus weights were not significantly different (Supplemental Figure 3c).

### Anterior insula (AI) inactivation can reduce avoidant adult male social behavior

After characterizing social behavior after defeat in both males and females, given the role of anterior insula (AI) in approach and avoidance behavior in healthy animals we wanted to determine if AI was necessary to the social avoidance phenotype seen in susceptible defeated males. Given that preliminary c-fos staining after social interaction revealed increased c-fos labeling in the AI of socially avoidant males exposed to eaCSDS as compared to controls (Figure 2a), we chose to test the necessity of increased AI activity to social avoidance by silencing AI in defeated and control animals. We performed bilateral injections of an adeno-associated viral vector enabling neuron-specific expression of the inhibitory DREADD construct (‘Gi’) into AI of young adult males and females subjected to eaCSDS. We then performed OFSI testing in adulthood (∼P90) on two different days, counterbalancing treatment with saline or deschloroclozapine (DCZ, 5 mg/kg), which acts as a specific DREADD ligand (Figure 2b). While silencing of AI did not significantly affect social interaction ratio in DREADD-injected control male and female mice or DREADD-injected susceptible defeated female mice, AI silencing did significantly improve social interaction ratios in susceptible defeated male mice injected with DREADDs (Figure 2c), inducing a “resilient” phenotype in 7 of the 8 susceptible mice with DREADDs injection. Additionally, DREADD-injected susceptible male mice showed a significant increase in social interaction ratio compared to susceptible defeated male mice that received saline and DCZ intraperitoneally that were injected with mCherry instead of DREADDs (Figure 2d). We then examined specific behaviors in the DREADDs injected male mice; while we found no differences in exploratory rearing or grooming with DCZ treatment, we found a significant decrease in the number of ‘startles backwards’, a behavior observed in susceptible male mice during approach towards the CFW contained under the wire cup (Figure 2e). We also did not observe biting behavior or tail rattling in any of the DREADDs injected susceptible male mice during saline or DCZ treatment (data not shown), suggesting AI silencing facilitates social approach rather than increasing aggression in males. Interestingly, AI silencing during auditory fear generalization testing did not affect fear generalization behavior in defeated male mice (Supplemental Figure 4a,b), suggesting that AI may play a unique role in social, but not auditory conditioned, fear generalization in males.

**Figure 2.**
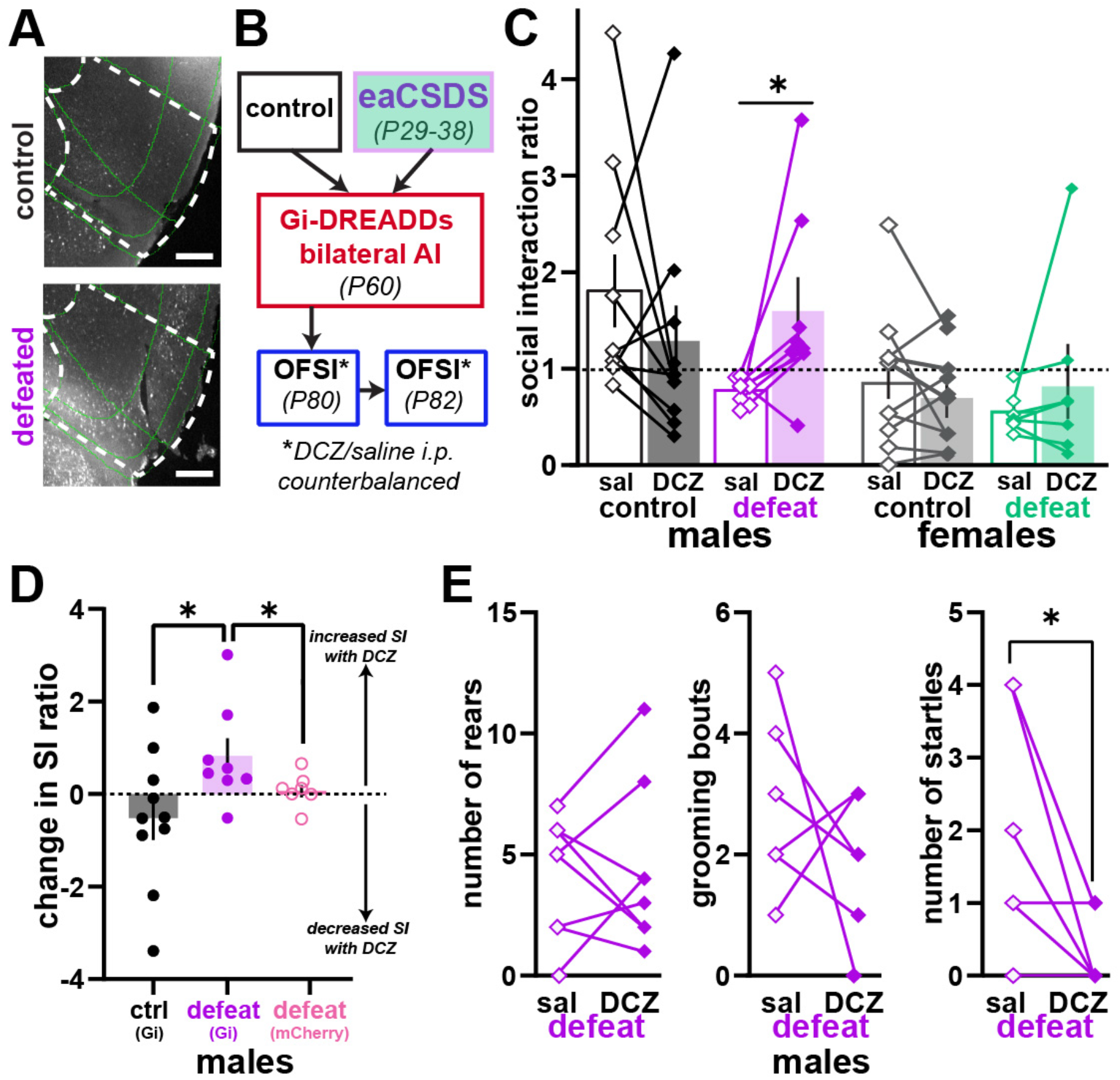
Behavioral avoidance in adult males after eaCSDS can be reduced with anterior insula (AI) inactivation. A) C-fos labeling in anterior insula in example control and defeated images. Scale bar, 500 um. B) Schematic of AI DREADDs inactivation experiment. C) Social interaction ratio during open field social interaction testing (OFSI) with saline or with DCZ for Gi-DREADDs injected control animals (black, n = 10 males) (grey, n = 8 females) and Gi-DREADDs injected susceptible defeated mice (purple, n = 8 males, 2-way repeated measures ANOVA, interaction effect, p = 0.049; post-hoc paired one tail t-test for defeated saline versus DCZ treatment, p = 0.034) (green, n = 7 females) . D) Change in social interaction ratio (social interaction ratio with DCZ - social interaction ratio with saline) (unpaired two tailed t-test for control versus defeat (Gi injected), p = 0.049; Mann Whitney two-tailed test for defeat (Gi injected) versus defeat (mCherry injected), p = 0.04). E) Paired comparisons of rearing, grooming and startle behavior (paired two-tailed t-test, p = 0.03, 1 defeated animal detected as outlier (ROUT (Q=1%) test) and removed).

### Social interaction in adulthood after eaCSDS reveals increased brainwide network density and decreased AI connectivity with other nodes in behaviorally avoidant adult males

We then wanted to further understand how activity and connectivity in AI and the brain more globally changes after early life maltreatment in socially avoidant adult males. First, we used whole brain c-fos to compare neuronal activation 1 hour after social interaction in control and susceptible defeated male mice (Figure 3a). We selected 6 defeated male mice that were “susceptible” to eaCSDS in adulthood (social interaction score < 1 in OFSI^12,32^) but showed no significant differences in distance traveled in comparison to controls (Figure 3b). As an initial analysis to determine additional regions of interest, we directly compared average c-fos density across hemispheres between control and susceptible defeated males across the brain (Supplemental Table 1). In addition to AI, we found over 20 regions with nominally significantly elevated c-fos density in susceptible defeated mice compared to controls in cortical, subcortical and hindbrain regions (Figure 3c). We then performed a comprehensive network analysis in which we found striking increases in connectivity between regions as measured by Pearson correlation in susceptible defeated mice not found in control mice (Figure 3d). Examination of network density revealed that susceptible defeated mice showed increased network density across multiple Pearson correlation coefficients (PCC) and when controlling for significance across multiple comparisons (Figure 3e, Methods). We then examined the connectivity between social, salience, and default mode networks^40^ (Figure 2f, PCC = 0.8). We observed a decrease in connectivity within both the social (between prelimbic cortex (PL), mediodorsal thalamus (MD) and central amygdala (CEA)) and salience (lack of connectivity between AI and all other salience nodes) networks specific to susceptible defeated mice. In the default mode network, we found an increase in connectivity between every node in susceptible defeated mice. Overall, both regional and network level comparisons revealed notable increases in regional and network activity in susceptible defeated mice with aberrant connectivity across social, salience and default mode networks. This finding is notable given its parallels with the human literature of aberrant default mode network connectivity in stress- and trauma-related disorders^41,42^.

**Figure 3.**
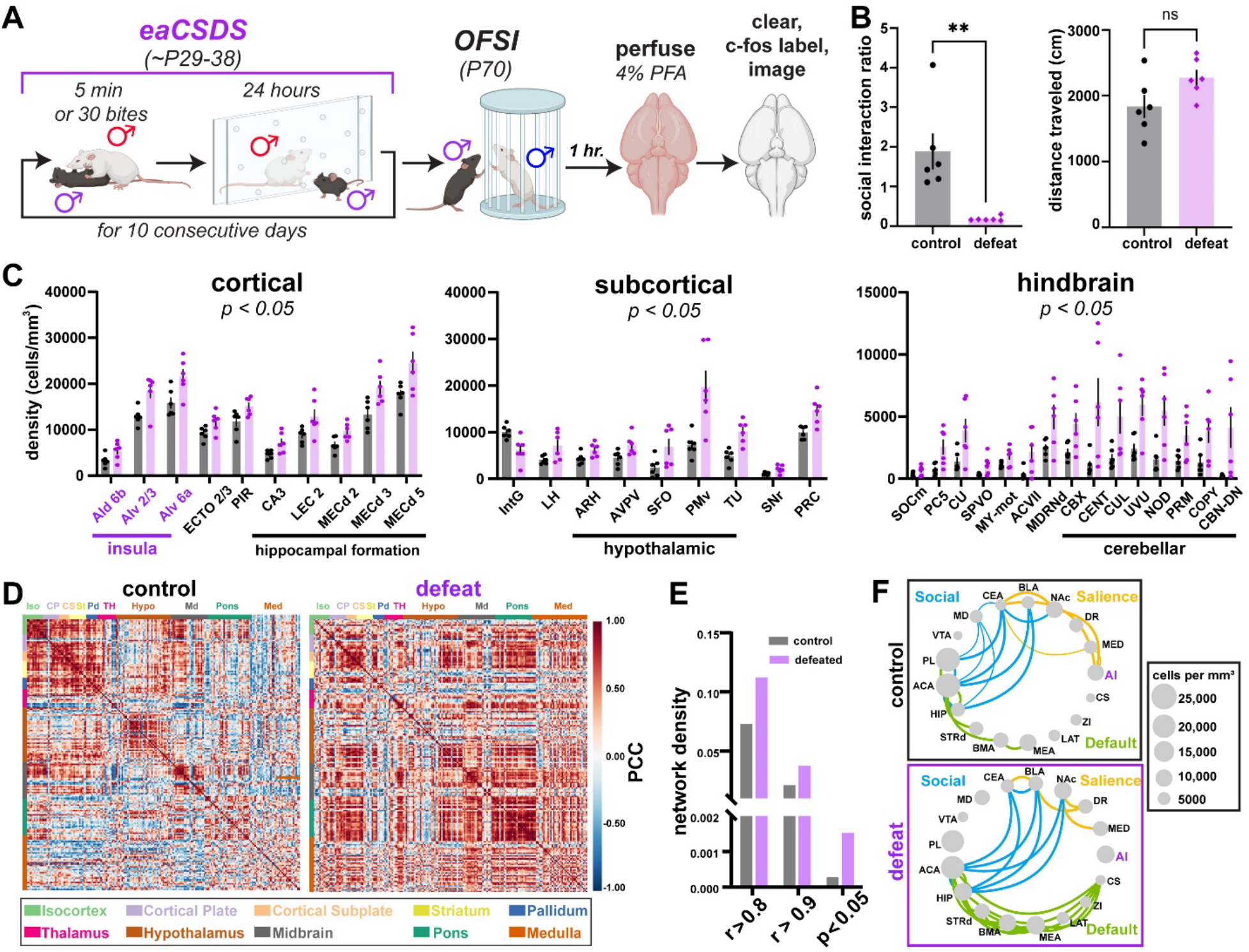
Social interaction testing in adulthood after eaCSDS results in increased regional activation and network density as revealed by whole brain c-fos labeling. A) Schematic of whole brain c-fos experiment. B) Social interaction ratio (unpaired two-tailed t-test, p = 0.004) and distanced traveled (unpaired two-tailed t-test, p = 0.07) for control (black, n = 6) and susceptible defeated (purple, n = 6) mice during OFSI. C) Regions of interest with significantly increased c-fos density in defeated compared to control mice (unpaired two-tailed t-test for control versus defeated for each region, p < 0.05). D) Pearson correlation coefficients (PCC) for c-fos density across multiple brain regions in control (left) and defeated (right) mice. E) Network density measures for varying correlation coefficients (PCC= 0.8 or 0.9) or for only p-value corrected regions. F) Network connectivity maps between social, salience and default mode networks for control and defeated mice (PCC = 0.8). Thickness of line indicates strength of correlated c-fos activity. Size of node indicates density of c-fos labeling in region. ACA: anterior cingulate cortex; ARH: arcuate hypothalamic; AVPV: anteroventral periventricular nucleus; AI: anterior insula; BLA: basolateral amygdala; BMA: basomedial amygdala; CBN-DN: dentate nucleus; CBX: cerebellar cortex; CEA: central amygdala; CENT: central lobule; COPY: copula pyramidis; cRN: central Raphe nuclei; CUL: culmen; DRN: dorsal Raphe nuclei; dStr: dorsal striatum; ECTO: ectorhinal cortex; HIP: hippocampal formation; IntG: intermediate geniculate; LAT: lateral group of dorsal thalamus; LEC: lateral entorhinal cortex; LH: lateral habenula; MD: mediodorsal nucleus of the thalamus; MEA: medial amygdala; MEC: medial entorhinal cortex; MED: medial group of dorsal thalamus; NAc: nucleus accumbens; NOD: nodulus (X); PC5: parvicellular motor nucleus; PIR: piriform cortex; PL: prelimbic region of prefrontal cortex; PMv: premamillary nucleus; PRC: precomissural nucleus; PRM: paramedian lobule; SFO: subfornical organ; SNr: substantia nigra pars reticulata; SOCm: superior olivary complex, medial region; TU: tuberal nucleus; UVU: uvula (IX); ZI: zona incerta.

We then wanted to confirm that the putative connectivity changes observed with c-fos labeling, namely the decreases in AI functional connectivity between other network nodes, were indicative of changes in neural circuit function and functional network activity using in vivo electrophysiology. We implanted adult defeated and control male mice with electrodes to record local field potentials (LFPs) targeting AI and 6 additional brain regions central to fear circuitry. This includes PL, MD, nucleus accumbens (NAc), ventral hippocampus (HIP), ventral tegmental area (VTA) and primary visual cortex (V1; control) (Supplemental Figure 4a). We then performed 5 minute recordings of spontaneous neural activity in the home cage with the implanted mouse alone (‘habituation’) followed by 5 minute recordings with an ovariectomized CFW female in the cage (‘social interaction’). We first examined activity within AI, given its increase in putative activity and decrease in putative functional connectivity with other brain regions observed with whole-brain c-fos. Power spectral density analysis of AI LFP revealed significant changes between control and defeated mice during social interaction in the delta (2-5 Hz) and theta (5-8 Hz) bands in both raw and differential power spectral density (Figure 4b, c; respectively). We then examined changes in information flow between AI and the 6 other sites using Granger causality analysis (Figure 4d). Most strikingly, this analysis revealed a profound disruption in information flow from AI to 5 of the 6 sites (MD, NAc, HIP, VTA, V1, Figure 4e) in defeated mice, suggesting a robust dysregulation of information transfer out of AI. In combination, whole brain c-fos analysis and multi-site in vivo LFP recordings reveal robust changes in functional connectivity in adulthood after eaCSDS in socially avoidant male mice.

**Figure 4.**
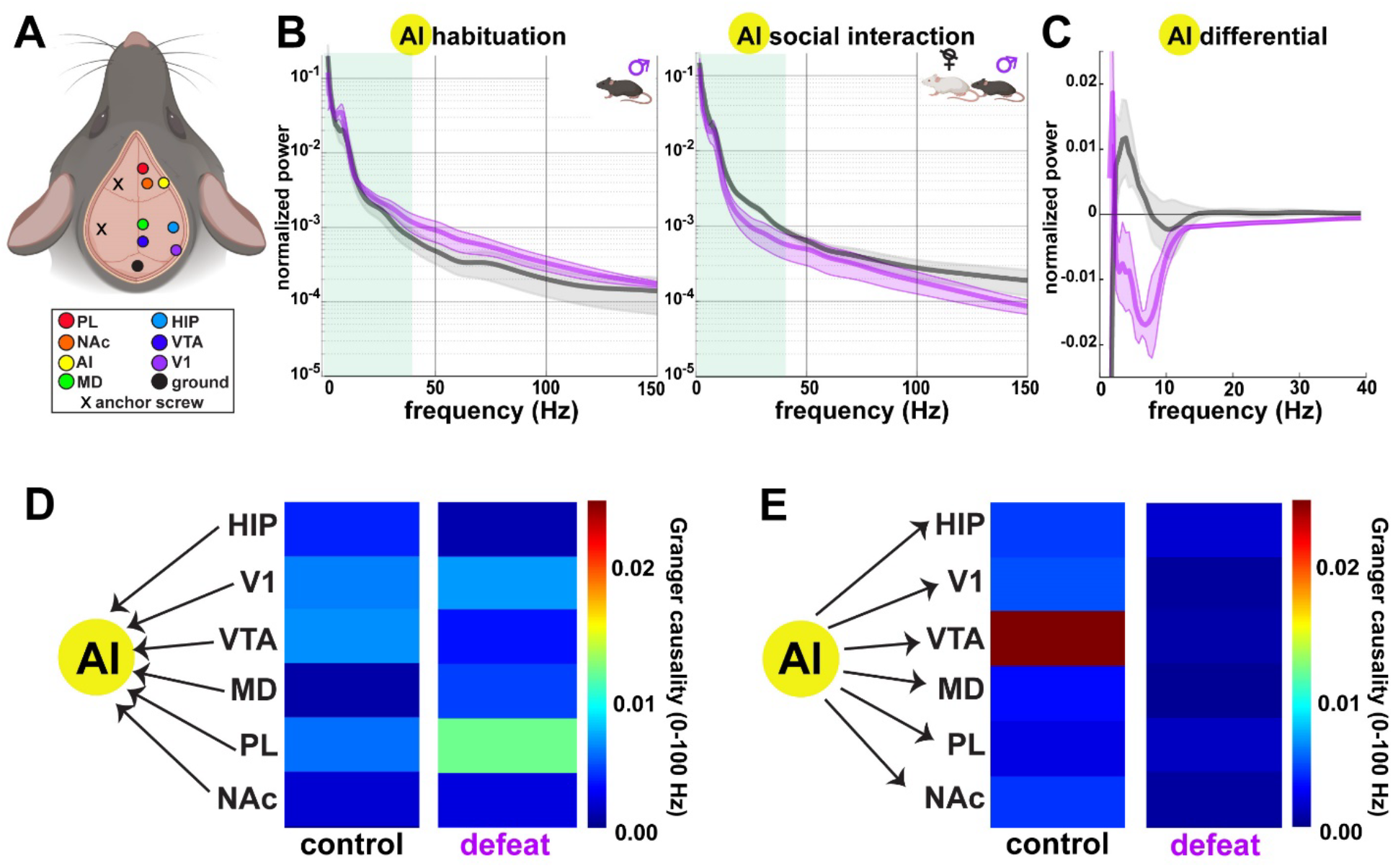
Multi-site LFP recordings reveal changes in AI oscillatory activity and decreased functional connectivity in defeated males in adulthood. A) Schematic of electrode implant sites. B) Power density spectrogram during habituation (left) and social interaction with an ovariectomized CFW female (right) in control (grey, n = 3) and defeated (purple, n = 3) male mice. Here we observed a significant decrease in both delta (Con: 0.352 ± 0.032 Normalized Power; Defeated: 0.201± 0.008; p = 0.012 t-test) and beta (Con: 0.067 ± 0.003 Normalized Power; Defeated: 0.035± 0.011; p = 0.049 t-test) power during social interaction comparing defeated mice to controls. C) Differential power density spectrogram (habituation power subtracted from social interaction power) shows significant difference in the change in delta (Con: 0.064 ± 0.035 Δ Normalized Power; Defeated: - 0.066± 0.030; p = 0.048 t-test) and beta (Con: 0.005 ± 0.013 Normalized Power; Defeated: -0.080± 0.019; p = 0.021 t-test) power induced by the shift from homecage to social interaction in defeated mice versus controls. D) Pairwise conditional Granger causality (time) for input to AI. E) Pairwise conditional Granger causality (time) for output from AI.

## DISCUSSION

Overall, we have shown that chronic stress in early adolescence has profound impacts on behavior, neural activity, and gene expression in adulthood. In males, eaCSDS results in robust social avoidance behavior that generalizes to unfamiliar mice of the same strain as the aggressor and to age matched mice of the same strain. Neural activity in susceptible adult males is altered across numerous brain regions, as evidenced by whole brain c-fos and multi-site LFP recordings; the anterior insula (AI) in particular shows marked increases in c-fos labeling and decreases in functional communication with frontal and subcortical regions in susceptible males. Chemogenetic silencing of AI neurons can ameliorate maladaptive social behavior in defeated males.

Surprisingly, early adolescent defeat in females has a very different impact on social behavior; instead of avoidance, adult females show no change in social interaction ratio in OFSI and a lack of avoidant behaviors in direct social interaction. Indeed, previous work using female defeats or witness of defeats in adulthood has shown a lack of impact on social behavior as measured by social interaction ratio in OFSI^43,44^. This is not to say their social behavior is typical as a marked increase in social contact in defeated females is observed with both different-strain and same-strain mice. The difference in male-avoidance and female-hypervigilance/affiliative phenotypes after early life stress has also been described in humans^45,46^ and may reflect a cross-species sex specific response to early life maltreatment. Future work will examine the neurobiological changes underlying this aberrant increased social contact in female mice.

AI has gained attention in the human literature for its role as a hub of the salience network that enables switching between attentive and restful brain states^18,19^. The loss of the ability to modulate switching between these states is thought to underlie symptoms of PTSD and anxiety^47,48^. The insular cortex has also been implicated in social approach behavior towards stressed conspecifics in healthy rats^25,49,50^ but its role in social processing in stressed animals themselves was previously unknown. Our findings support the notion that AI dysfunction after trauma underlies aberrant behavioral responding and raise the novel possibility that normalization of AI activity with targeted non-invasive neural circuit therapies such as transcranial magnetic stimulation (TMS)^51,52^ or focused ultrasound^53^ could potentially improve psychiatric symptoms.

## METHODS AND MATERIALS

### Animals

Three week-old male C57BL/6J (C57) mice were obtained from Jackson Laboratory (Bar Harbor, ME, USA) or bred in house. Adult CFW males and ovariectomized females were obtained from Charles River Laboratory (Boston, MA, USA). The mice were housed under a 12-hour light and 12-hour dark schedule temperature (21 ±2 C and humidity (50 ±20%), and food and water were available *ad libitum*. Following arrival at McLean Hospital, the mice were given seven days to habituate to the vivarium prior to starting the defeat. All experimental procedures were approved by the Institutional Animal Care and Use Committee at McLean Hospital and VA Boston and were performed in accordance with the National Institutes of Health’s (NIH) Guide for the Care and Use of Animals.

### Early adolescent chronic social defeat stress (eaCSDS)

As was described previously, virgin male CFW mice (8 weeks, Charles River) were housed with ovariectomized female CFW (8 weeks, Charles River) mice for at least 1 week and screened for aggression with C57 male mice prior to defeat sessions. Only male CFWs that show an attack latency of less than 30 seconds for 2 consecutive days are used for defeats. Before the start of the first defeat, female CFW mice were permanently removed from the cage.

Male C57 mice (P29 ±3 days) were exposed to a traditional 10-day defeat paradigm, with slight modifications. Specifically, C57 juveniles were placed into the home cage of an aggressive male CFW mouse. Each session consists of physical interactions that proceed until the aggressor has delivered 30 bites. Defeat sessions end after 5 minutes maximum if fewer than 30 bites occur. The juvenile mice are then housed side-by-side with the male CFW aggressor overnight using a plastic cage divider that physically separates the mice but enables visual and olfactory contact. This procedure was repeated each day with a novel aggressor for 10 consecutive days. At 24 hours after the last defeat, each defeated C57 mice was re-housed with another defeated mouse but separated by plastic divider, to control for potential effects of social isolation in adolescence. Control mice were housed with other control mice but separated by a plastic divider from P29 onward, to ensure consistent housing conditions across conditions.

### Open Field Social Interaction with CFW male (OFSI)

Mice are transported from the animal facility to the behavioral testing room. After 1 hour of habituation to the testing room, C57 male mice (∼P39 for testing 24 hours after defeat at early adolescent time point, ∼P50 for testing 2 weeks after defeat at late adolescent time point, ∼P70 for testing 5 weeks at adult time point) are placed opposite an inverted black wire cup in a large (44 x 44 cm) arena. After 150 seconds of exploration of the arena and empty cup, the C57 is placed back in the arena with the cup now containing a novel nonaggressive CFW male inside and allowed to explore for 150 seconds. All interactions are captured and scored with Ethovision. Social interaction ratio is calculated as time spent with the cup with a CFW inside divided by time spent with the empty cup. Mice with a social interaction ratio ≥1 are classified as “resilient” while mice with a social interaction ratio <1 are classified as “susceptible”.

### Home Cage Social Interaction with ovariectomized CFW female (HCSI)

Adolescent (∼P50) C57 male mice are singly housed the night before testing. The following day they remain in their home cage and an ovariectomized CFW female is placed inside the home cage for 90 seconds. If any biting is observed from either male or female, interaction is immediately ended, and the female is replaced with another female. The interaction is recorded at 30 fps and later tracked and analyzed for specific behaviors with DeepLabCut and SimBA.

### Fear Generalization Training and Testing

Adult (>P70) C57 male mice are transported from the animal facility to a holding room outside the facility. After they are habituated to the room for 1 hour, mice are placed in conditioning chambers (context A, lights on, quatricide for cleaning, shock grid floor) and undergo fear generalization training (3 minutes of habituation followed by 5 30-second 12 kHz tones terminating in a one-second 0.7 MA shock (CS+) and 5 30-second 3 kHz tones that never terminate in a shock (CS-), randomly interleaved over 15 minutes (10 tones total)). The following day, mice are again habituated to the holding room for 1 hour then placed in a novel context (context B, lights off, 50% ethanol for cleaning, opaque black plexiglass floor). Mice then undergo fear generalization testing (3 minutes of habituation followed by 2 30-seccond 3, 6, 9, or 12 kHz tones randomly interleaved over 20 minutes (8 tones total)). All video from training and testing was recorded at 30 frames per second and analyzed for percentage freezing in 30 second bins with FreezeFrame 5.

### Tissue collection for whole brain c-fos imaging

5 weeks after defeat (∼P70), mice were habituated to a behavioral testing room within the animal facility for 1 hour before OFSI testing. After OFSI, mice are placed in a clean empty cage in a quiet room for 1 hour and then perfused with 0.1% heparin in 1x PBS followed by 4% PFA. Brains are removed and placed in PFA overnight then stored in 0.02% sodium azide PBS. Only brains without any visible redness or damage were used for tissue clearing followed by whole brain c-fos labeling and imaging (LifeCanvas Technologies).

### Multisite LFP implants

Stereotaxic surgeries were performed in adult male mice under isofluorane anesthesia as previously described. LFP electrodes (polyimide coated tungsten; 500 kΩ impedance) were stereotaxically implanted unilaterally targeting PL (AP:1.6mm, ML:0.4mm, DV:-2.3mm); HIP (AP:-2.5mm, ML:2.7mm, DV:-2.3mm); AI (AP:1.8mm, ML:2.8mm, DV:-1.9mm); VTA (AP:-3mm, ML:0.3mm, DV:-4.65mm); NAc (AP:1.1mm, ML:0.8mm, DV:-4.5mm); V1 (AP:-4.0mm, ML:2.5mm, DV:-1.5mm); and MD (AP:-1.2mm, ML:0.5mm, DV:-3.2mm). A ground screw was mounted over the cerebellum and one anchoring screw was mounted contralaterally. LFP signals were recorded with a 16-channel system (Intan Tech).

### Viral injections

Stereotaxic viral injections were performed in adult male mice under isofluorane anesthesia as previously described. AAV virus was injected in the anterior insula (AI) bilaterally (from brain surface in millimeters: Allen Mouse Brain Reference Atlas: AP: +1.80 mm, ML: 2.75 mm, DV:1.90 mm) (AAV2/5 hSyn-DREADDS(Gi)-mCherry or AAV2/5 hSyn-mCherry for silencing experiments) over a period of ∼10 minutes (7 injections of 32 nL every 20 seconds) with a pressure injection system (Nanoject). All viruses were allowed to incubate for >1 month before behavioral testing.

### Histology

For post-mortem confirmation of AI injection sites, mice were deeply anesthetized with isofluorane and perfused with 1x PBS followed by 4% paraformaldehyde. Brains were then submerged in 4% paraformaldehyde overnight then transferred to 30% sucrose in 1x PBS for at least 24 hours. After brains sunk to the bottom of the sucrose solution, they were embedded in TissueTek and cut in 50 um sections on a Cryostat. Sections were placed in wells filled with PBS then mounted on slides and coverslipped with FluoroMount. Sections were examined under the fluorescent microscope for mCherry labeling in AI bilaterally. Any animals that did not show bilateral mCherry labeling restricted to AI over 4 consecutive sections was excluded from analysis.

## QUANTIFICATION AND STATISTICAL ANALYSIS

All data were analyzed in GraphPad Prism 10. Data were first tested for normality using the Shapiro-Wilk normality test. If data passed the normality test (p > 0.05), parametric analyses were used; if the normality test failed, non-parametric analyses were used.

### c-fos network analysis

We constructed correlation matrices using cFos density data collected from 196 brain regions. The network density represents the degree of interconnectedness among brain regions and is calculated as the proportion of observed connections relative to the total possible connections (10.31887/DCNS.2018.20.2/osporns). We also focused on the connections within three specific functional networks: the social network, the salience network, and the default mode network (https://doi.org/10.3389/fnins.2019.00585)(https://doi.org/10.1038/s41380-021-01298-5). For the analysis, we used Python libraries, including SciPy(10.1038/s41592-019-0686-2), Statsmodels (https://scholar.google.com/scholar_lookup?author=S+Seabold&author=J+Perktold&publication_year=2010&title=Statsmodels%3A%20Econometric%20and%20statistical%20modeling%20with%20python&journal=SciPy&volume=7&pages=1), and Seaborn (https://doi.org/10.21105/joss.03021), to construct the correlation matrices, perform network analyses, and carry out statistical tests. We applied the Shapiro-Wilk test to assess the normality of the data, Pearson’s correlation for the analysis, and the Benjamini-Hochberg FDR correction to account for multiple comparisons.

### LFP activity analysis

LFP data from each cortical site targeted was analyzed using custom written scripts for MATLAB (2023B, Mathworks). Power spectral density analysis was performed on established frequency bands that are suggested to have physiologically relevance (delta, theta, gamma etc.) using multitaper (Cronux Toolbox). Granger causality analysis was used to assess the directional flow of information between targeted regions using the MVGC toolbox (Trongnetrpunya et al., 2015) as described in prior studies (Nair et al., 2018).

## ACKNOWLEDGMENTS

National Institutes of Health grant T32-MH125786 (MGK and EEH)

Eric Dorris Memorial Research Award (EEH)

Brain and Behavior Foundation Young Investigator Award (EEH)

National Institutes of Health grant K99-MH133869 (ELN)

National Institutes of Health grant P50-MH115874 (WAC, KJR)

National Institutes of Health grant R01-MH115874 (KJR)

McLean Lorenz Pope Fellowship (MGK)

Dupont Warren Award (MGK) McLean Frazier Fund (KJR)

## AUTHOR CONTRIBUTIONS

Conceptualization, MGK, ELN, WAC, KJR, EEH; methodology, MGK, ELN, EEH; Investigation, MGK, JO, CK, LD, CM,ZB, MM, ZW, OF, JM, EEH; writing—original draft, EEH; writing—review & editing, MGK, EEH, OF, WAC, JH, TK, JM, KJR; funding acquisition, MGK, ELN, WAC, KJR, EEH; resources, WAC and KJR; supervision, WAC, KJR, EEH.

