## Supplemental Figures for "A critical role for the anterior insula in impaired adult social interaction after early life maltreatment"

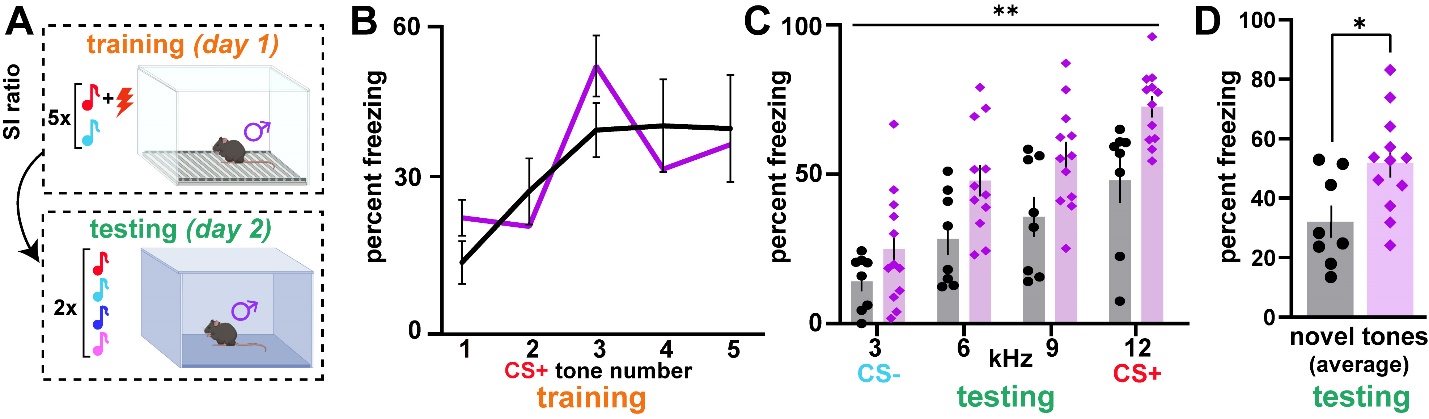

**Figure S1: Fear generalization training and testing in adult males defeated as adolescents.**

A) Schematic for auditory fear generalization protocol in adult (>P70) mice defeated as adolescents. Mice are conditioned to a CS+ (12 kHz) paired with a shock and to a CS- (3 kHz) never paired with a shock during training (day 1). During testing (day 2) mice are presented with the CS-, CS+ and two novel tones (6 kHz and 9 kHz). B) Percent freezing to 5 presentations of CS+ during training (day 1) for control (black, n = 8) and defeated (purple, n = 12) (2-way repeated measures ANOVA, main effect of defeat status, p = 0.98). C) Average percent freezing to 2 presentations of CS- (3 kHz), 6 kHz, 9 kHz, CS+ (12 kHz) during testing (day 2) (2-way repeated measures ANOVA, main effect of defeat status, p = 0.0058). D) Average percent freezing to novel tones (6 kHz and 9kHz) during testing (day 2) (unpaired two-tailed t-test, p = 0.016). *, p < 0.05; **, p < 0.01; ***, p < 0.001; for all figures shown. Error bars indicate standard error of the mean for all figures shown.

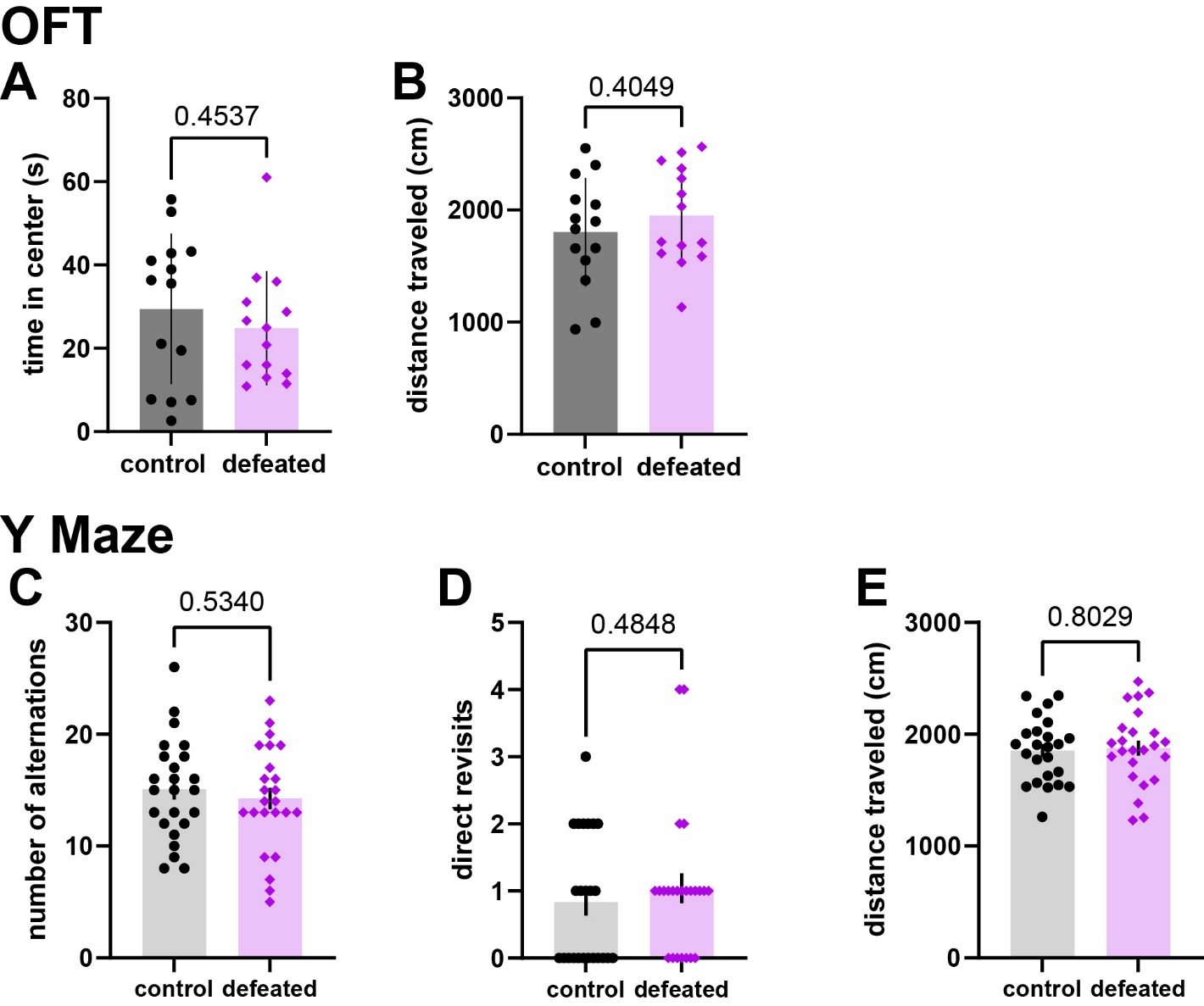

**Figure S2: Behavioral performance of control and defeated adult male mice in tests of anxiety-like behavior and working memory.**

Behavioral performance of control and defeated adult male mice in tests of anxiety-like behavior (open field test (OFT)) and working memory (Y maze). Control (black) and defeated (purple) mice compared with unpaired two-tailed t-tests. N = 14 control mice and 14 defeated mice for S2A,B. A) Time spent in center of open field during 10 minute OFT (p = 0.4537). B) Distance traveled during OFT (p = 0.4049). N = 24 control mice and 24 defeated mice for S2C-E. C) Number of alternations made in 5 minute Y maze testing (p = 0.5340). D) Number of direct revisits to same arm of Y maze (p = 0.4848). E) Distance traveled during Y maze testing (p = 0.8029).

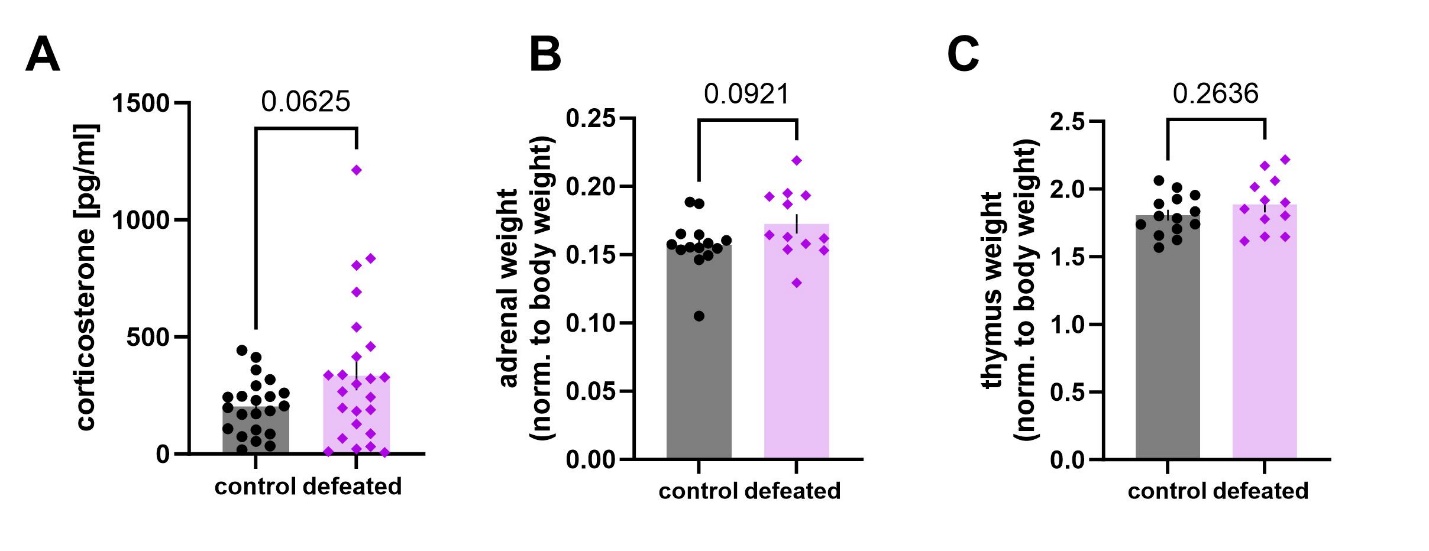

**Figure S3: Biomarkers of stress exposure in control versus defeated adult male mice.**

Traditional biomarkers used to assay stress exposure were collected in defeated and control adult (~P70) male mice. A) Free corticosterone levels (pg/ml) in adult control (black, n = 22 mice) and adult defeated (purple, n = 24 mice) collected from trunk blood and measured via ELISA 2 hours after exposure to an unfamiliar, non-aggressive CFW male (unpaired two-tailed t-test, p = 0.0625). B) Adrenal weights normalized to body weight from control (n = 14 mice) and defeated (n = 12 mice) mice (unpaired two-tailed t-test, p = 0.0921). C) Thymus weights normalized to body weight from control (n = 14 mice) and defeated (n = 12 mice) mice (unpaired two-tailed t-test, p = 0.2636).

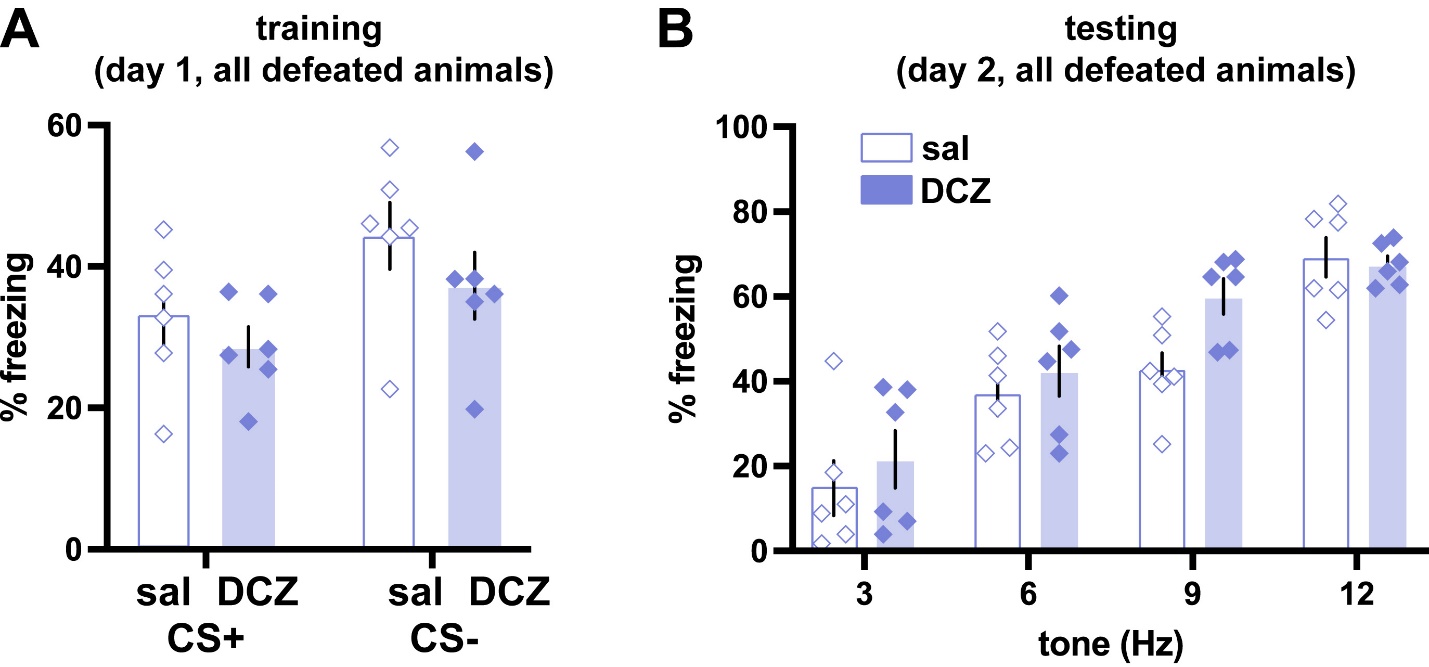

**Figure S4: Chemogenetic silencing of AI does not affect fear recall after classical conditioning based auditory fear generalization training.**

A) Training (day 1) in adult mice defeated as adolescents and injected with Gi DREADDS in AI as adults to be treated with either saline or DCZ during testing (grey, saline treated, n = 6 mice; blue, DCZ treated on testing day, n = 6 mice) (Two-way repeated measures ANOVA, main effect of DCZ treatment: F =1.151, p = 0.309) B) Testing (day 2) in adult mice defeated as adolescents and injected with Gi DREADDS in AI as adults treated with either saline or DCZ during testing (Two-way repeated measures ANOVA, main effect of DCZ treatment: F = 1.728, p = 0.218).

| **Supplemental table 1** | **density (cells/mm^3^)** |  |  |  |
| --- | --- | --- | --- | --- |
|  | **control** |  | **defeat** |  |
| **region** | average | standard error | average | standard  error |
| background | 113.846 | 12.7736 | 96.3489 | 9.84294 |
| left root | 8313.01 | 509.258 | 9895.46 | 773.394 |
| left Basic cell groups and regions | 8863.02 | 545.059 | 10503.5 | 818.294 |
| left Cerebrum | 12414 | 823.234 | 14122.6 | 947.703 |
| left Cerebral cortex | 13747.7 | 884.626 | 15616.1 | 1037.72 |
| left Cortical plate | 14015 | 898.807 | 15849.1 | 1047.21 |
| left Isocortex | 15717.3 | 1141.04 | 18226.2 | 1285.13 |
| left Frontal pole, cerebral cortex | 12518.1 | 2616.05 | 12193.5 | 2000.63 |
| left Frontal pole, layer 1 | 3627.88 | 370.638 | 5737.02 | 1801.35 |
| left Frontal pole, layer 2/3 | 18308.5 | 4964.12 | 14785.2 | 2860.46 |
| left Frontal pole, layer 5 | 16220.2 | 3492.06 | 15660.5 | 3035.39 |
| left Frontal pole, layer 6a | 8304.55 | 1071.18 | 9864.44 | 1663.25 |
| left Frontal pole, layer 6b | 8080.81 | 1244.9 | 15191.9 | 2941.83 |
| left Somatomotor areas | 10233.4 | 762.695 | 10827.9 | 2074.68 |
| left Primary motor area | 8171.36 | 497.876 | 9263.51 | 1747.18 |
| left Primary motor area, Layer 1 | 3375.42 | 400.006 | 2683.03 | 435.991 |
| left Primary motor area, Layer 2/3 | 9147.69 | 683.124 | 9872.93 | 1778.32 |
| left Primary motor area, Layer 5 | 8540.5 | 514.68 | 9555.15 | 2190.13 |
| left Primary motor area, Layer 6a | 8810.21 | 863.945 | 11228.5 | 2031.02 |
| left Primary motor area, Layer 6b | 6717.9 | 989.514 | 8694.06 | 1358.83 |
| left Secondary motor area | 12019.3 | 1147.01 | 12182.9 | 2396.8 |
| left Secondary motor area, layer 1 | 4240.99 | 765.031 | 2950.1 | 459.954 |
| left Secondary motor area, layer 2/3 | 14828.6 | 2011.8 | 14712.4 | 2697 |
| left Secondary motor area, layer 5 | 13214 | 1287.65 | 13777.8 | 3107.57 |
| left Secondary motor area, layer 6a | 12940.5 | 743.12 | 14120.2 | 2643.76 |
| left Secondary motor area, layer 6b | 9811.03 | 1210.11 | 12945.4 | 1318.02 |
| left Somatosensory areas | 10779.9 | 1011.76 | 14155.2 | 1506.34 |
| left Primary somatosensory area | 11515.8 | 1180.46 | 15076.5 | 1762.49 |
| left Primary somatosensory area, nose | 7124.36 | 1436.13 | 14728.2 | 2851.01 |
| left Primary somatosensory area, nose, layer 1 | 5849.41 | 1347 | 6890.6 | 731.439 |
| left Primary somatosensory area, nose, layer 2/3 | 9541.8 | 2112.6 | 21051.7 | 4114.19 |
| left Primary somatosensory area, nose, layer 4 | 6484.03 | 1327.97 | 16837 | 5280.24 |
| left Primary somatosensory area, nose, layer 5 | 6547.55 | 1518.71 | 14488.9 | 3556.25 |
| left Primary somatosensory area, nose, layer 6a | 6908.13 | 1252.47 | 12867.6 | 2349.98 |
| left Primary somatosensory area, nose, layer 6b | 3781.93 | 808.742 | 7370.72 | 2327.99 |
| left Primary somatosensory area, barrel field | 18827.4 | 2499.62 | 22927.2 | 2419.87 |
| left Primary somatosensory area, barrel field, layer 1 | 7690.01 | 1362.63 | 8449.48 | 864.744 |
| left Primary somatosensory area, barrel field, layer 2/3 | 21725.7 | 2803.02 | 32631.1 | 3559.96 |
| left Primary somatosensory area, barrel field, layer 4 | 23105.6 | 3505.73 | 28699.5 | 4160.23 |
| left Primary somatosensory area, barrel field, layer 5 | 16113.8 | 2055.52 | 17217.9 | 2396.05 |
| left Primary somatosensory area, barrel field, layer 6a | 21046.9 | 2789.04 | 21256.5 | 2618.58 |
| left Primary somatosensory area, barrel field, layer 6b | 17863.3 | 2574.78 | 16778.6 | 2539.24 |
| left Primary somatosensory area, lower limb | 15504.7 | 2347.22 | 15520.1 | 2202.03 |
| left Primary somatosensory area, lower limb, layer 1 | 4565.69 | 656.232 | 4433.93 | 1537.17 |
| left Primary somatosensory area, lower limb, layer 2/3 | 17368.4 | 3003.47 | 19943.8 | 2979.38 |
| left Primary somatosensory area, lower limb, layer 4 | 15063.6 | 3010.21 | 15695.2 | 2435.16 |
| left Primary somatosensory area, lower limb, layer 5 | 14740.4 | 1898.2 | 14595.3 | 2573.63 |
| left Primary somatosensory area, lower limb, layer 6a | 20815.4 | 3060.7 | 17716.9 | 2636.27 |
| left Primary somatosensory area, lower limb, layer 6b | 11631.6 | 1691.77 | 10725.7 | 1269.36 |
| left Primary somatosensory area, mouth | 3728.32 | 518.767 | 6327.08 | 1236.24 |
| left Primary somatosensory area, mouth, layer 1 | 2918.09 | 680.41 | 2339.68 | 318.568 |
| left Primary somatosensory area, mouth, layer 2/3 | 5068.56 | 750.187 | 7087.53 | 1384.28 |
| left Primary somatosensory area, mouth, layer 4 | 3837.27 | 590.224 | 6312.18 | 1728.46 |
| left Primary somatosensory area, mouth, layer 5 | 3490.12 | 516.931 | 6100.79 | 1609.22 |
| left Primary somatosensory area, mouth, layer 6a | 3108.2 | 455.969 | 7733.38 | 1865.61 |
| left Primary somatosensory area, mouth, layer 6b | 2299.63 | 391.657 | 5808.67 | 1738.86 |
| left Primary somatosensory area, upper limb | 11690.4 | 1360.4 | 14321.6 | 2414.12 |
| left Primary somatosensory area, upper limb, layer 1 | 5083.67 | 757.765 | 5067.05 | 829.212 |
| left Primary somatosensory area, upper limb, layer 2/3 | 13501.6 | 1661.41 | 18264 | 2554.05 |
| left Primary somatosensory area, upper limb, layer 4 | 11412 | 1592.69 | 14132.6 | 2550.64 |
| left Primary somatosensory area, upper limb, layer 5 | 11874.9 | 1122.57 | 14522.6 | 3457.07 |
| left Primary somatosensory area, upper limb, layer 6a | 13332.4 | 1883.44 | 15142 | 2835.46 |
| left Primary somatosensory area, upper limb, layer 6b | 8050.57 | 1293.79 | 8652.9 | 1625.57 |
| left Primary somatosensory area, trunk | 13993 | 1760.32 | 17147.2 | 2494.25 |
| left Primary somatosensory area, trunk, layer 1 | 3376.31 | 663.161 | 3386.09 | 622.546 |
| left Primary somatosensory area, trunk, layer 2/3 | 14941.5 | 2223.98 | 21641.9 | 3499.36 |
| left Primary somatosensory area, trunk, layer 4 | 16800.8 | 2516.79 | 20414.1 | 3647.74 |
| left Primary somatosensory area, trunk, layer 5 | 13624.7 | 1728.04 | 16068.9 | 2551.02 |
| left Primary somatosensory area, trunk, layer 6a | 19507.8 | 2487.86 | 20394 | 2917.08 |
| left Primary somatosensory area, trunk, layer 6b | 19539.4 | 2341.43 | 18210.9 | 2173.07 |
| left Primary somatosensory area, unassigned | 13187.7 | 1530.1 | 18957.5 | 3275.72 |
| left Primary somatosensory area, unassigned, layer 1 | 7419.77 | 1403.12 | 9363.9 | 1316.25 |
| left Primary somatosensory area, unassigned, layer 2/3 | 14708.6 | 2100.53 | 24898.6 | 4270.59 |
| left Primary somatosensory area, unassigned, layer 4 | 13346.6 | 1606.07 | 18352.3 | 2639.81 |
| left Primary somatosensory area, unassigned, layer 5 | 12371.7 | 1217.41 | 17517.9 | 3803.2 |
| left Primary somatosensory area, unassigned, layer 6a | 15493.6 | 1971.95 | 20035.9 | 4150.12 |
| left Primary somatosensory area, unassigned, layer 6b | 11551.1 | 1214.73 | 10881.1 | 2618.36 |
| left Supplemental somatosensory area | 8797.78 | 725.754 | 11674 | 1083.7 |
| left Supplemental somatosensory area, layer 1 | 4885.68 | 1045.07 | 3542.47 | 434.804 |
| left Supplemental somatosensory area, layer 2/3 | 10487.6 | 1004.02 | 14819 | 1427.5 |
| left Supplemental somatosensory area, layer 4 | 9253.17 | 906.916 | 14584.2 | 2056.27 |
| left Supplemental somatosensory area, layer 5 | 8396.22 | 724.388 | 11340.7 | 1270.37 |
| left Supplemental somatosensory area, layer 6a | 9538.53 | 891.654 | 12090 | 1523.19 |
| left Supplemental somatosensory area, layer 6b | 9122.89 | 1055.29 | 11055 | 1551.68 |
| left Gustatory areas | 5616.15 | 613.089 | 7270.46 | 1183.1 |
| left Gustatory areas, layer 1 | 3501.6 | 808.687 | 1835.14 | 448.688 |
| left Gustatory areas, layer 2/3 | 6568.25 | 847.677 | 7735.2 | 1613.48 |
| left Gustatory areas, layer 4 | 5639.46 | 571.561 | 8183.67 | 1897.21 |
| left Gustatory areas, layer 5 | 5348.32 | 738.01 | 8236.84 | 1489.95 |
| left Gustatory areas, layer 6a | 6145.16 | 711.525 | 7888.57 | 1138.43 |
| left Gustatory areas, layer 6b | 4137.07 | 945.217 | 5865.01 | 674.144 |
| left Visceral area | 7428.56 | 939.867 | 8076.78 | 908.905 |
| left Visceral area, layer 1 | 4333.3 | 1214.27 | 2627.57 | 411.587 |
| left Visceral area, layer 2/3 | 7751.26 | 925.372 | 8877.63 | 1186.47 |
| left Visceral area, layer 4 | 6970.01 | 977.094 | 9298.21 | 1540.44 |
| left Visceral area, layer 5 | 7231.05 | 1125.82 | 8156.81 | 1044.12 |
| left Visceral area, layer 6a | 9228.51 | 1078.13 | 10014 | 1263.83 |
| left Visceral area, layer 6b | 10464.8 | 1214.02 | 10592.6 | 1223.06 |
| left Auditory areas | 17303.7 | 1480.38 | 21209.5 | 1685.65 |
| left Dorsal auditory area | 13890.1 | 1986.73 | 20667.9 | 2132.27 |
| left Dorsal auditory area, layer 1 | 2527.39 | 593.27 | 2282.8 | 467.44 |
| left Dorsal auditory area, layer 2/3 | 14268.3 | 2036.83 | 23336.4 | 1885.69 |
| left Dorsal auditory area, layer 4 | 17334 | 3098.77 | 29806.1 | 4858.51 |
| left Dorsal auditory area, layer 5 | 15047.5 | 2248.48 | 20718.7 | 2450.54 |
| left Dorsal auditory area, layer 6a | 19657 | 2583.3 | 27543.7 | 3077.13 |
| left Dorsal auditory area, layer 6b | 20097.1 | 2903.14 | 28499.1 | 3023.94 |
| left Primary auditory area | 17321.5 | 1712.09 | 20620.4 | 1720.15 |
| left Primary auditory area, layer 1 | 3480.93 | 757.114 | 3850.53 | 594.322 |
| left Primary auditory area, layer 2/3 | 18323.6 | 1309.63 | 24427.2 | 1376.72 |
| left Primary auditory area, layer 4 | 27167.9 | 4008.39 | 32052.5 | 3679.53 |
| left Primary auditory area, layer 5 | 19805.3 | 2325.23 | 21096.9 | 2424.59 |
| left Primary auditory area, layer 6a | 19025.5 | 2026.63 | 24703.5 | 3942.17 |
| left Primary auditory area, layer 6b | 20758.1 | 1633.42 | 26688.1 | 5004.05 |
| left Posterior auditory area | 15217.3 | 2101.49 | 18824.3 | 2169.6 |
| left Posterior auditory area, layer 1 | 2148.62 | 533.793 | 2889.32 | 274.832 |
| left Posterior auditory area, layer 2/3 | 16556.2 | 2559.43 | 23542.2 | 1641.48 |
| left Posterior auditory area, layer 4 | 24308.2 | 4797.82 | 28127.3 | 5147.47 |
| left Posterior auditory area, layer 5 | 16972.2 | 2241.87 | 18473.7 | 2780.38 |
| left Posterior auditory area, layer 6a | 17269.6 | 1820.81 | 22643 | 3337.51 |
| left Posterior auditory area, layer 6b | 22196.2 | 2251.79 | 28897.3 | 4388 |
| left Ventral auditory area | 20254.7 | 1008.68 | 23065.8 | 2971.82 |
| left Ventral auditory area, layer 1 | 4688.58 | 893.424 | 2544.74 | 393.828 |
| left Ventral auditory area, layer 2/3 | 22127.5 | 1787.5 | 26274.7 | 2350.86 |
| left Ventral auditory area, layer 4 | 31590.9 | 2673.96 | 37014.8 | 3546.79 |
| left Ventral auditory area, layer 5 | 21987.4 | 1294.25 | 24726.6 | 4317.7 |
| left Ventral auditory area, layer 6a | 21468.6 | 1531.08 | 25981.4 | 4638.47 |
| left Ventral auditory area, layer 6b | 20331.2 | 1016.34 | 21553.5 | 4583.82 |
| left Visual areas | 29834.6 | 2406.38 | 34825 | 3469.19 |
| left Anterolateral visual area | 20127.9 | 3160.01 | 28213.8 | 3385.32 |
| left Anterolateral visual area, layer 1 | 1668.88 | 541.05 | 2195 | 248.037 |
| left Anterolateral visual area, layer 2/3 | 24560.4 | 4306.94 | 38280.2 | 3132.45 |
| left Anterolateral visual area, layer 4 | 23975 | 3149.94 | 38102.6 | 6093.52 |
| left Anterolateral visual area, layer 5 | 22101.4 | 3183.8 | 28331.4 | 4351.29 |
| left Anterolateral visual area, layer 6a | 22978.9 | 3930.61 | 27314.8 | 3966.3 |
| left Anterolateral visual area, layer 6b | 27771.9 | 4846.48 | 33364.9 | 4821.1 |
| left Anteromedial visual area | 24536.5 | 2543.87 | 29342.6 | 4213.36 |
| left Anteromedial visual area, layer 1 | 2373.14 | 434.357 | 2069.02 | 547.379 |
| left Anteromedial visual area, layer 2/3 | 29859 | 2328.34 | 34501.6 | 4317.65 |
| left Anteromedial visual area, layer 4 | 33662.7 | 3572.44 | 41749.7 | 5934.42 |
| left Anteromedial visual area, layer 5 | 26044 | 4206.4 | 34432.2 | 6432.66 |
| left Anteromedial visual area, layer 6a | 29915.3 | 3853.51 | 32585.1 | 5565.15 |
| left Anteromedial visual area, layer 6b | 33797.6 | 3296.55 | 38370.9 | 5317.56 |
| left Lateral visual area | 28276.8 | 2699.17 | 32666.6 | 3281.75 |
| left Lateral visual area, layer 1 | 1282.15 | 155.867 | 1860.38 | 305.915 |
| left Lateral visual area, layer 2/3 | 24790.4 | 2430.68 | 37888.3 | 2291.36 |
| left Lateral visual area, layer 4 | 35980.7 | 3807.6 | 44183.8 | 4623.41 |
| left Lateral visual area, layer 5 | 37400.2 | 3710.94 | 36656.1 | 4483.19 |
| left Lateral visual area, layer 6a | 34269.8 | 4008.98 | 35374.2 | 5031.12 |
| left Lateral visual area, layer 6b | 43000 | 4407.98 | 39521.4 | 5547.29 |
| left Primary visual area | 33165 | 3440.69 | 37674.6 | 3868.89 |
| left Primary visual area, layer 1 | 3466.04 | 386.562 | 4240.23 | 790.131 |
| left Primary visual area, layer 2/3 | 35936.3 | 4281.38 | 46750.1 | 4252.34 |
| left Primary visual area, layer 4 | 43714.1 | 3845.71 | 47676.6 | 4837.85 |
| left Primary visual area, layer 5 | 37859.7 | 4848.96 | 40514.7 | 5359.15 |
| left Primary visual area, layer 6a | 42971 | 5539.22 | 44594.5 | 5927.44 |
| left Primary visual area, layer 6b | 52831.6 | 6845.87 | 49272.1 | 6678.08 |
| left Posterolateral visual area | 32854.4 | 4852.89 | 32824.6 | 2119.56 |
| left Posterolateral visual area, layer 1 | 7857.8 | 4160.66 | 5870.53 | 1650.31 |
| left Posterolateral visual area, layer 2/3 | 31980.8 | 7391.73 | 31065.9 | 3169.41 |
| left Posterolateral visual area, layer 4 | 46158.1 | 4731.06 | 48659.8 | 2978.95 |
| left Posterolateral visual area, layer 5 | 46077.5 | 4953.64 | 45835 | 3504.95 |
| left Posterolateral visual area, layer 6a | 43271.3 | 3827.71 | 47841.9 | 4921.78 |
| left Posterolateral visual area, layer 6b | 45020.2 | 6031.09 | 47411.6 | 6394.24 |
| left posteromedial visual area | 33679.7 | 3388.64 | 41977.2 | 5311.31 |
| left posteromedial visual area, layer 1 | 2755.38 | 514.798 | 4591.71 | 1806.93 |
| left posteromedial visual area, layer 2/3 | 40664 | 5505.73 | 49061 | 5914.14 |
| left posteromedial visual area, layer 4 | 46075.8 | 4035.32 | 57080.4 | 9802.61 |
| left posteromedial visual area, layer 5 | 38395.6 | 4440.7 | 47012.6 | 5786.06 |
| left posteromedial visual area, layer 6a | 39891.1 | 4688.98 | 52470.8 | 7955.46 |
| left posteromedial visual area, layer 6b | 42678.3 | 4328.95 | 57304.2 | 7902.07 |
| left Laterointermediate area | 25838.5 | 2374.71 | 31120.8 | 3700.05 |
| left Laterointermediate area, layer 1 | 1593.31 | 286.8 | 2097.63 | 441.443 |
| left Laterointermediate area, layer 2/3 | 23081.8 | 1468.26 | 33989.7 | 3599.71 |
| left Laterointermediate area, layer 4 | 35131.1 | 3376.24 | 43794.1 | 5139.61 |
| left Laterointermediate area, layer 5 | 34426.7 | 3835.81 | 37538.4 | 5264.95 |
| left Laterointermediate area, layer 6a | 27758.4 | 3260.25 | 30962.6 | 4801.79 |
| left Laterointermediate area, layer 6b | 37386.6 | 4696.88 | 42027.9 | 6295.8 |
| left Postrhinal area | 18493.1 | 2362.48 | 25296.7 | 3037.87 |
| left Postrhinal area, layer 1 | 1280.37 | 332.627 | 1208.43 | 202.29 |
| left Postrhinal area, layer 2/3 | 14038.3 | 1386.8 | 21976.1 | 2204.07 |
| left Postrhinal area, layer 4 | 24556.5 | 2774.14 | 32275.5 | 4281.93 |
| left Postrhinal area, layer 5 | 27118.4 | 4614.84 | 36258.7 | 4405.92 |
| left Postrhinal area, layer 6a | 28845.3 | 3382.67 | 37153.2 | 5976.26 |
| left Postrhinal area, layer 6b | 31334.6 | 5013.33 | 40291.7 | 5154.39 |
| left Anterior cingulate area | 21563 | 2468.88 | 21373.1 | 2072.85 |
| left Anterior cingulate area, dorsal part | 24190.6 | 2546.73 | 23981.7 | 2916.76 |
| left Anterior cingulate area, dorsal part, layer 1 | 16396.5 | 1943.7 | 17774.1 | 1092.71 |
| left Anterior cingulate area, dorsal part, layer 2/3 | 30899 | 3216.11 | 30418.4 | 4419.27 |
| left Anterior cingulate area, dorsal part, layer 5 | 24185.9 | 2919.06 | 24484 | 3657.11 |
| left Anterior cingulate area, dorsal part, layer 6a | 25204.2 | 2277.23 | 23052 | 2665.57 |
| left Anterior cingulate area, dorsal part, layer 6b | 13231.6 | 1762.41 | 12905.9 | 1695.29 |
| left Anterior cingulate area, ventral part | 18148.8 | 2386.89 | 17983.7 | 1628.63 |
| left Anterior cingulate area, ventral part, layer 1 | 10408.2 | 2014.73 | 12573.8 | 1644.64 |
| left Anterior cingulate area, ventral part, layer 2/3 | 24379.4 | 3210.53 | 23772.8 | 2490.19 |
| left Anterior cingulate area, ventral part, layer 5 | 19768.4 | 2486.9 | 19234.5 | 1678.71 |
| left Anterior cingulate area, ventral part, 6a | 17037.8 | 2033.04 | 15617.9 | 1409.43 |
| left Anterior cingulate area, ventral part, 6b | 6956.52 | 1262.48 | 6050.56 | 1198.99 |
| left Prelimbic area | 21239.1 | 2520.11 | 19813.9 | 1538.9 |
| left Prelimbic area, layer 1 | 5652.4 | 877.358 | 5257.49 | 668.618 |
| left Prelimbic area, layer 2/3 | 24742.5 | 3249.02 | 20999.2 | 1940.28 |
| left Prelimbic area, layer 5 | 26184.5 | 3380.82 | 24833.8 | 2061.18 |
| left Prelimbic area, layer 6a | 25376.1 | 2290.23 | 24517.9 | 2144.8 |
| left Prelimbic area, layer 6b | 6758.15 | 1512.25 | 10043.8 | 1844.42 |
| left Infralimbic area | 17631.5 | 2092.38 | 16835.9 | 1616.79 |
| left Infralimbic area, layer 1 | 2811.93 | 1137.36 | 1750.05 | 333.814 |
| left Infralimbic area, layer 2/3 | 12860.6 | 2681.15 | 10415.1 | 1790.52 |
| left Infralimbic area, layer 5 | 23830 | 3152.63 | 22659.8 | 2098.68 |
| left Infralimbic area, layer 6a | 21839.4 | 2102.06 | 22828.8 | 2439.37 |
| left Infralimbic area, layer 6b | 9537.25 | 1869.59 | 12015.7 | 1974.93 |
| left Orbital area | 25056.9 | 2500.59 | 28259.2 | 2313.06 |
| left Orbital area, lateral part | 25280.3 | 2214.76 | 29781.5 | 2992.7 |
| left Orbital area, lateral part, layer 1 | 4771.7 | 865.176 | 10961.2 | 2347.21 |
| left Orbital area, lateral part, layer 2/3 | 29210.5 | 4425.19 | 37449.7 | 3210.7 |
| left Orbital area, lateral part, layer 5 | 33482.7 | 3019.4 | 37981.6 | 4494.52 |
| left Orbital area, lateral part, layer 6a | 16995.7 | 1532.94 | 15839.4 | 1553.95 |
| left Orbital area, lateral part, layer 6b | 10203.5 | 2297.35 | 9108.2 | 1232.9 |
| left Orbital area, medial part | 19719.7 | 2649.28 | 19937.5 | 2054.17 |
| left Orbital area, medial part, layer 1 | 2601.61 | 534.023 | 4857.85 | 1495.88 |
| left Orbital area, medial part, layer 2/3 | 18055.5 | 3817.64 | 21017.3 | 3631.82 |
| left Orbital area, medial part, layer 5 | 31996.5 | 5142.23 | 29560.8 | 4157.49 |
| left Orbital area, medial part, layer 6a | 29387.4 | 1640.61 | 27109.2 | 2848.21 |
| left Orbital area, medial part, layer 6b | 11644.4 | 2108.81 | 14755.6 | 3169.4 |
| left Orbital area, ventrolateral part | 28887.3 | 3305 | 32430.3 | 2956.26 |
| left Orbital area, ventrolateral part, layer 1 | 5150.86 | 1645.54 | 9027.84 | 1352.49 |
| left Orbital area, ventrolateral part, layer 2/3 | 25334.8 | 5490.83 | 34084.4 | 4301.77 |
| left Orbital area, ventrolateral part, layer 5 | 44594.4 | 5767.5 | 46563 | 5914.98 |
| left Orbital area, ventrolateral part, layer 6a | 34724.1 | 2329.5 | 30701.7 | 3743 |
| left Orbital area, ventrolateral part, layer 6b | 17562.9 | 4356.4 | 13534.7 | 2449.99 |
| left Agranular insular area | 10477.8 | 1141.18 | 12456.5 | 1574.18 |
| left Agranular insular area, dorsal part | 9221.84 | 1192.69 | 10012 | 1535.2 |
| left Agranular insular area, dorsal part, layer 1 | 4891.24 | 714.665 | 3347.17 | 551.601 |
| left Agranular insular area, dorsal part, layer 2/3 | 11141.3 | 1705.31 | 10788.7 | 1633.35 |
| left Agranular insular area, dorsal part, layer 5 | 10264 | 1341.34 | 11593.5 | 2019.78 |
| left Agranular insular area, dorsal part, layer 6a | 7913.62 | 828.021 | 10245.6 | 1386.27 |
| left Agranular insular area, dorsal part, layer 6b | 2623.71 | 450.121 | 5710.04 | 1009.49 |
| left Agranular insular area, posterior part | 8077.39 | 1112.74 | 10718.8 | 1228.12 |
| left Agranular insular area, posterior part, layer 1 | 2074.71 | 498.024 | 1287.02 | 96.5998 |
| left Agranular insular area, posterior part, layer 2/3 | 9777.29 | 1388.03 | 13818.7 | 2316.07 |
| left Agranular insular area, posterior part, layer 5 | 7277.92 | 1050.35 | 10178.6 | 1316.27 |
| left Agranular insular area, posterior part, layer 6a | 11931.3 | 1462.22 | 14923.7 | 1857.64 |
| left Agranular insular area, posterior part, layer 6b | 12355.1 | 2291.95 | 13735.2 | 2982.72 |
| left Agranular insular area, ventral part | 16534.7 | 1342.66 | 20134.1 | 2617.1 |
| left Agranular insular area, ventral part, layer 1 | 5058.36 | 577.595 | 4271.98 | 697.955 |
| left Agranular insular area, ventral part, layer 2/3 | 15463.7 | 1053.11 | 20201.3 | 2712.3 |
| left Agranular insular area, ventral part, layer 5 | 21614.8 | 2165.11 | 25417.2 | 3710.03 |
| left Agranular insular area, ventral part, layer 6a | 15862.5 | 1578.97 | 21091.9 | 1917.72 |
| left Agranular insular area, ventral part, layer 6b | 5222.22 | 1443.73 | 9888.89 | 2687.04 |
| left Retrosplenial area | 21510.9 | 1342.27 | 24132.8 | 1859.06 |
| left Retrosplenial area, lateral agranular part | 24516.9 | 1640.39 | 29360.9 | 3131.85 |
| left Retrosplenial area, lateral agranular part, layer 1 | 4054.72 | 536.109 | 5319.18 | 1608.81 |
| left Retrosplenial area, lateral agranular part, layer 2/3 | 29351.6 | 2753.54 | 35472.2 | 4328.16 |
| left Retrosplenial area, lateral agranular part, layer 5 | 29816.8 | 2501.86 | 35048.1 | 3010.02 |
| left Retrosplenial area, lateral agranular part, layer 6a | 32096.5 | 2185.73 | 38353.8 | 5037.7 |
| left Retrosplenial area, lateral agranular part, layer 6b | 27984 | 2576.56 | 35412.2 | 5217.8 |
| left Retrosplenial area, dorsal part | 22095 | 1811.46 | 24179.4 | 2106.73 |
| left Retrosplenial area, dorsal part, layer 1 | 7092.39 | 797.126 | 5819.64 | 986.64 |
| left Retrosplenial area, dorsal part, layer 2/3 | 24224.4 | 2183.41 | 27395.2 | 2948.98 |
| left Retrosplenial area, dorsal part, layer 5 | 25773.8 | 2209.42 | 28199.8 | 2375.58 |
| left Retrosplenial area, dorsal part, layer 6a | 29804 | 2711.05 | 33734.8 | 4055.9 |
| left Retrosplenial area, dorsal part, layer 6b | 26077.5 | 2036.76 | 25785.6 | 3055.14 |
| left Retrosplenial area, ventral part | 19364.2 | 1083.08 | 21249 | 1683.2 |
| left Retrosplenial area, ventral part, layer 1 | 11816 | 1185.79 | 13059 | 898.256 |
| left Retrosplenial area, ventral part, layer 2/3 | 21446.7 | 1270.75 | 25867.2 | 1295.95 |
| left Retrosplenial area, ventral part, layer 5 | 22510 | 1286.58 | 23745.2 | 3010.49 |
| left Retrosplenial area, ventral part, layer 6a | 18877.9 | 1767.08 | 19803.4 | 2141.53 |
| left Retrosplenial area, ventral part, layer 6b | 17299.7 | 1796.79 | 13744.1 | 1544 |
| left Posterior parietal association areas | 24029.6 | 3204.69 | 29398.3 | 3936.26 |
| left Anterior area | 22322.6 | 3180.55 | 28137.6 | 4031.45 |
| left Anterior area, layer 1 | 3712.58 | 952.371 | 3604.63 | 666.937 |
| left Anterior area, layer 2/3 | 25801.5 | 4162.02 | 35015.7 | 5308.71 |
| left Anterior area, layer 4 | 30335 | 5152.71 | 39074.6 | 5809.23 |
| left Anterior area, layer 5 | 22535.5 | 3395.12 | 28984.2 | 4437.55 |
| left Anterior area, layer 6a | 27894.7 | 3014.9 | 30846.7 | 4410.43 |
| left Anterior area, layer 6b | 31685.8 | 3605.79 | 35448.1 | 5348.12 |
| left Rostrolateral visual area | 26445.6 | 3931.73 | 31182.5 | 3963.3 |
| left Rostrolateral area, layer 1 | 3216.12 | 676.483 | 5121.02 | 1158.81 |
| left Rostrolateral area, layer 2/3 | 32130.5 | 4893.71 | 41904.1 | 4788.52 |
| left Rostrolateral area, layer 4 | 32408.8 | 4354.05 | 37691 | 6147 |
| left Rostrolateral area, layer 5 | 26680.4 | 3911.51 | 29654.3 | 4835.67 |
| left Rostrolateral area, layer 6a | 34074.3 | 6088.01 | 35915.2 | 4789.97 |
| left Rostrolateral area, layer 6b | 36213.9 | 5604.68 | 36776.5 | 6338.03 |
| left Temporal association areas | 17798.3 | 597.11 | 20992.7 | 1665.7 |
| left Temporal association areas, layer 1 | 3410.94 | 847.111 | 2404.86 | 212.261 |
| left Temporal association areas, layer 2/3 | 16709.2 | 651.538 | 20675.5 | 1500.97 |
| left Temporal association areas, layer 4 | 24706.4 | 1328.97 | 29163.7 | 2026.96 |
| left Temporal association areas, layer 5 | 20524.4 | 807.042 | 24030.2 | 2302.62 |
| left Temporal association areas, layer 6a | 24076 | 1238.68 | 29074.3 | 3388.95 |
| left Temporal association areas, layer 6b | 22744.5 | 1265.25 | 26743.6 | 3172.49 |
| left Perirhinal area | 10403.8 | 561.849 | 14152.1 | 1998.37 |
| left Perirhinal area, layer 1 | 3984.28 | 603.816 | 3855.51 | 938.423 |
| left Perirhinal area, layer 2/3 | 13521.2 | 558.558 | 19058.9 | 2802.88 |
| left Perirhinal area, layer 5 | 10704.3 | 942.873 | 15462.7 | 2340.24 |
| left Perirhinal area, layer 6a | 14186.9 | 1289.75 | 19435.9 | 2699.62 |
| left Perirhinal area, layer 6b | 20882.6 | 2866.51 | 25965.6 | 1914.61 |
| left Ectorhinal area | 13260.6 | 606.235 | 16313.4 | 1341.86 |
| left Ectorhinal area/Layer 1 | 4544.96 | 972.286 | 3710.58 | 550.447 |
| left Ectorhinal area/Layer 2/3 | 12906.9 | 848.334 | 16833.8 | 2049.48 |
| left Ectorhinal area/Layer 5 | 13577.3 | 626.559 | 17235.3 | 1895.53 |
| left Ectorhinal area/Layer 6a | 18389.5 | 1287.82 | 21969.2 | 2098.85 |
| left Ectorhinal area/Layer 6b | 18880.9 | 1510.54 | 23574.7 | 2882.8 |
| left Olfactory areas | 14226.2 | 533.521 | 13283.6 | 731.592 |
| left Main olfactory bulb | 16714.6 | 300.258 | 11507.8 | 1433.51 |
| left Accessory olfactory bulb | 20480.6 | 2344.28 | 13325.7 | 2184.53 |
| left Accessory olfactory bulb, glomerular layer | 5671.79 | 1104.37 | 7103.21 | 2599.18 |
| left Accessory olfactory bulb, granular layer | 27640.1 | 5522.71 | 16869.1 | 1528.86 |
| left Accessory olfactory bulb, mitral layer | 23058.6 | 3378.21 | 13982.1 | 3321.61 |
| left Anterior olfactory nucleus | 21707.9 | 1498.57 | 23467 | 1923.97 |
| left Taenia tecta | 9116.91 | 1293.38 | 10244.4 | 1152.91 |
| left Taenia tecta, dorsal part | 12863.1 | 2023.54 | 14361 | 1505.22 |
| left Taenia tecta, ventral part | 5359.32 | 818.182 | 6115.19 | 971.168 |
| left Dorsal peduncular area | 15086.2 | 1458.96 | 15723.8 | 1535.19 |
| left Piriform area | 11564.8 | 1081.19 | 14693.4 | 1320.78 |
| left Nucleus of the lateral olfactory tract | 12930.1 | 1133.01 | 17690.3 | 2228.37 |
| left Nucleus of the lateral olfactory tract, molecular layer | 5153.86 | 570.929 | 6876.8 | 820.284 |
| left Nucleus of the lateral olfactory tract, pyramidal layer | 18735.8 | 1863.96 | 25498.3 | 3229.22 |
| left Nucleus of the lateral olfactory tract, layer 3 | 12906.2 | 1565.75 | 18343.4 | 2583.53 |
| left Cortical amygdalar area | 8907.3 | 1199.8 | 10900.4 | 1384.82 |
| left Cortical amygdalar area, anterior part | 14151.3 | 1678.34 | 19512.6 | 2460.42 |
| left Cortical amygdalar area, posterior part | 7295.63 | 1218.82 | 8253.52 | 1442.85 |
| left Cortical amygdalar area, posterior part, lateral zone | 5274.41 | 742.846 | 6519.52 | 1254.98 |
| left Cortical amygdalar area, posterior part, medial zone | 9171.57 | 1748.91 | 9862.88 | 1632.51 |
| left Piriform-amygdalar area | 4917.07 | 644.169 | 6896.1 | 832.531 |
| left Postpiriform transition area | 4464.44 | 823.986 | 5177.85 | 528.584 |
| left Hippocampal formation | 8862.63 | 755.244 | 11787.6 | 1264.57 |
| left Hippocampal region | 5767.77 | 578.924 | 8140.85 | 1276.34 |
| left Ammon's horn | 6006.75 | 634.101 | 8558.06 | 1431.98 |
| left Field CA1 | 7272.18 | 840.251 | 9510.94 | 1621.27 |
| left Field CA2 | 4047.38 | 674.64 | 6969.59 | 1792.58 |
| left Field CA3 | 4093.06 | 338.551 | 7125.62 | 1163.71 |
| left Dentate gyrus | 5202.01 | 590.723 | 7140.77 | 935.953 |
| left Dentate gyrus, molecular layer | 3636.4 | 580.173 | 5372.67 | 990.908 |
| left Dentate gyrus, polymorph layer | 7621.06 | 903.819 | 10149.9 | 1404.02 |
| left Dentate gyrus, granule cell layer | 8375.78 | 616.167 | 10606.6 | 1164.07 |
| left Fasciola cinerea | 6569.14 | 1254.25 | 8980.73 | 2306.7 |
| left Induseum griseum | 2748.35 | 914.763 | 3821.88 | 1159.29 |
| left Retrohippocampal region | 12702.3 | 1036.74 | 16407.1 | 1353.89 |
| left Entorhinal area | 11190.4 | 856.04 | 16248.8 | 1454.46 |
| left Entorhinal area, lateral part | 11467.3 | 977.373 | 17603.5 | 2007.4 |
| left Entorhinal area, lateral part, layer 1 | 1659.15 | 260.391 | 2592.56 | 676.135 |
| left Entorhinal area, lateral part, layer 2 | 10538.2 | 1184.91 | 18225.1 | 2212.78 |
| left Entorhinal area, lateral part, layer 3 | 14880.2 | 1055.67 | 23274.5 | 2473.99 |
| left Entorhinal area, lateral part, layer 5 | 11256.3 | 1047.79 | 16920.4 | 2545.75 |
| left Entorhinal area, lateral part, layer 6a | 18062.9 | 2101.14 | 24496.8 | 3551.83 |
| left Entorhinal area, medial part, dorsal zone | 10839.3 | 1004.93 | 14531.6 | 1272.69 |
| left Entorhinal area, medial part, dorsal zone, layer 1 | 3616.59 | 967.887 | 4417.54 | 822.882 |
| left Entorhinal area, medial part, dorsal zone, layer 2 | 7183.52 | 1177.51 | 10598 | 1073.5 |
| left Entorhinal area, medial part, dorsal zone, layer 3 | 13163.9 | 1240.04 | 18160 | 1010.36 |
| left Entorhinal area, medial part, dorsal zone, layer 5 | 16007.1 | 1521.03 | 21715.7 | 2342.77 |
| left Entorhinal area, medial part, dorsal zone, layer 6 | 21349.8 | 2395.35 | 26358.7 | 2903.91 |
| left Parasubiculum | 11688.9 | 1457.89 | 15485.8 | 1878.21 |
| left Postsubiculum | 21282.1 | 3646.02 | 19710 | 2816.5 |
| left Presubiculum | 18599.4 | 2685.9 | 19857.2 | 2267.07 |
| left Subiculum | 14307.4 | 1363.83 | 15991.9 | 1648.59 |
| left Prosubiculum | 10466.9 | 1329.28 | 12220 | 1653.19 |
| left Hippocampo-amygdalar transition area | 8210.84 | 1009.59 | 9957.56 | 2236.99 |
| left Area prostriata | 27919.6 | 4201.2 | 31766.2 | 4903.01 |
| left Cortical subplate | 7368.81 | 830.716 | 10054.2 | 1262.31 |
| left Claustrum | 12104.4 | 1624.19 | 15747.9 | 1538.08 |
| left Endopiriform nucleus | 9639.67 | 1008.24 | 12158 | 1501.44 |
| left Endopiriform nucleus, dorsal part | 11843.5 | 1225.45 | 14430.3 | 1767.72 |
| left Endopiriform nucleus, ventral part | 5521.47 | 716.896 | 7911.76 | 1056.31 |
| left Lateral amygdalar nucleus | 3966.31 | 1116.02 | 5241.56 | 858.543 |
| left Basolateral amygdalar nucleus | 4252.18 | 730.631 | 6727.32 | 1464.65 |
| left Basolateral amygdalar nucleus, anterior part | 5512.62 | 890.68 | 8516.12 | 2511.8 |
| left Basolateral amygdalar nucleus, posterior part | 3326.74 | 631.142 | 5580.5 | 961.278 |
| left Basolateral amygdalar nucleus, ventral part | 3504.89 | 667.08 | 5383.87 | 858.247 |
| left Basomedial amygdalar nucleus | 7890 | 910.671 | 10754.2 | 1342.23 |
| left Basomedial amygdalar nucleus, anterior part | 11050.4 | 1154.1 | 13537.7 | 1643.24 |
| left Basomedial amygdalar nucleus, posterior part | 4505.08 | 783.889 | 7773 | 1161.07 |
| left Posterior amygdalar nucleus | 6736.22 | 667.428 | 10901.6 | 1340.6 |
| left Cerebral nuclei | 6971.31 | 700.739 | 8028.03 | 645.545 |
| left Striatum | 7547.95 | 758.532 | 8648.2 | 731.598 |
| left Striatum dorsal region | 6410.04 | 642.868 | 7399.35 | 767.296 |
| left Caudoputamen | 6410.04 | 642.868 | 7399.35 | 767.296 |
| left Striatum ventral region | 8838.67 | 1234.34 | 9710.9 | 992.313 |
| left Nucleus accumbens | 10383.9 | 1474.28 | 11315.8 | 1468.16 |
| left Fundus of striatum | 5883.33 | 1355.97 | 7116 | 965.322 |
| left Olfactory tubercle | 7392.94 | 1157.26 | 8155.56 | 557.064 |
| left Lateral septal complex | 12660.8 | 1262.26 | 13035 | 366.382 |
| left Lateral septal nucleus | 13802.5 | 1320.62 | 14000.6 | 419.432 |
| left Lateral septal nucleus, caudal (caudodorsal) part | 6033.18 | 1084.07 | 6249.2 | 526.118 |
| left Lateral septal nucleus, rostral (rostroventral) part | 13790.3 | 1503.89 | 14116.5 | 324.612 |
| left Lateral septal nucleus, ventral part | 21048.9 | 1548.28 | 20834.2 | 1297.63 |
| left Septofimbrial nucleus | 5559.54 | 964.262 | 7176.52 | 804.082 |
| left Septohippocampal nucleus | 8287.39 | 1992.56 | 7494.29 | 1268.04 |
| left Striatum-like amygdalar nuclei | 7865.89 | 738.112 | 11086.2 | 1430.19 |
| left Anterior amygdalar area | 6433.05 | 1042.06 | 9939.59 | 1663.28 |
| left Bed nucleus of the accessory olfactory tract | 6307.25 | 1787.17 | 6214.49 | 753.398 |
| left Central amygdalar nucleus | 4291 | 388.515 | 6653.1 | 1464.49 |
| left Central amygdalar nucleus, capsular part | 5134.45 | 438.743 | 7418.7 | 2308.16 |
| left Central amygdalar nucleus, lateral part | 2657.37 | 414.657 | 4763.53 | 1363.37 |
| left Central amygdalar nucleus, medial part | 4510 | 416.088 | 6993.14 | 1409.55 |
| left Intercalated amygdalar nucleus | 7937.69 | 590.197 | 10799.5 | 1574.21 |
| left Medial amygdalar nucleus | 10553.3 | 1036.74 | 14335.6 | 1603.01 |
| left Pallidum | 4197.46 | 508.355 | 5044.78 | 443.535 |
| left Pallidum, dorsal region | 1173.09 | 184.64 | 1656.45 | 322.116 |
| left Globus pallidus, external segment | 1245.11 | 195.266 | 1849.21 | 387.956 |
| left Globus pallidus, internal segment | 907.316 | 154.859 | 945.088 | 154.739 |
| left Pallidum, ventral region | 4627.47 | 770.826 | 6199.66 | 907.766 |
| left Substantia innominata | 4870.64 | 776.469 | 6374.7 | 936.789 |
| left Magnocellular nucleus | 2608.63 | 892.638 | 4746.53 | 722.945 |
| left Pallidum, medial region | 4108.36 | 617.539 | 5005.46 | 413.227 |
| left Medial septal complex | 3455.32 | 658.562 | 3974.52 | 414.14 |
| left Medial septal nucleus | 3757.43 | 745.374 | 3881.05 | 365.726 |
| left Diagonal band nucleus | 3289.78 | 794.896 | 4025.74 | 682.792 |
| left Triangular nucleus of septum | 6212.29 | 670.699 | 8326.89 | 1053.89 |
| left Pallidum, caudal region | 9329.35 | 870.035 | 8912.92 | 370.897 |
| left Bed nuclei of the stria terminalis | 9370.11 | 873.311 | 8937.92 | 374.511 |
| left Bed nucleus of the anterior commissure | 2044.44 | 667.16 | 4444.44 | 1370.83 |
| left Brain stem | 3804.94 | 182.309 | 4785.79 | 589.163 |
| left Interbrain | 5454.99 | 273.069 | 6302.61 | 643.8 |
| left Thalamus | 4906.93 | 262.342 | 5641.65 | 713.428 |
| left Thalamus, sensory-motor cortex related | 3794.92 | 493.86 | 4418.77 | 599.905 |
| left Ventral group of the dorsal thalamus | 3418.87 | 698.421 | 4078 | 667.141 |
| left Ventral anterior-lateral complex of the thalamus | 4591.56 | 1230.98 | 4077.9 | 1337.17 |
| left Ventral medial nucleus of the thalamus | 4207.48 | 703.935 | 4182.27 | 1108.05 |
| left Ventral posterior complex of the thalamus | 2738.85 | 661.779 | 3992.96 | 932.622 |
| left Ventral posterolateral nucleus of the thalamus | 1272.13 | 172.065 | 2428.02 | 676.151 |
| left Ventral posterolateral nucleus of the thalamus, parvicellular part | 2212.95 | 557.267 | 2584.02 | 376.966 |
| left Ventral posteromedial nucleus of the thalamus | 3268.81 | 1007.56 | 4874.53 | 1396.38 |
| left Ventral posteromedial nucleus of the thalamus, parvicellular part | 4925.11 | 618.906 | 4287.86 | 700.854 |
| left Posterior triangular thalamic nucleus | 4269.77 | 1073.32 | 4595.46 | 979.941 |
| left Subparafascicular nucleus | 9268.32 | 717.154 | 9196.36 | 1000.01 |
| left Subparafascicular nucleus, magnocellular part | 15668.2 | 1095.85 | 17245.6 | 2285.01 |
| left Subparafascicular nucleus, parvicellular part | 5970.26 | 686.567 | 5048.33 | 658.054 |
| left Subparafascicular area | 8323.13 | 506.939 | 10132.2 | 837.201 |
| left Peripeduncular nucleus | 4291.75 | 1012.6 | 5246.08 | 1690.62 |
| left Geniculate group, dorsal thalamus | 3860.64 | 648.85 | 4341.46 | 875.037 |
| left Medial geniculate complex | 4054.93 | 654.755 | 4503.47 | 862.991 |
| left Medial geniculate complex, dorsal part | 4040.43 | 716.63 | 5638.4 | 1045.89 |
| left Medial geniculate complex, ventral part | 3347.84 | 567.618 | 4618.17 | 1032.2 |
| left Medial geniculate complex, medial part | 4848.24 | 1029.98 | 3595.54 | 638.807 |
| left Dorsal part of the lateral geniculate complex | 3672.77 | 784.289 | 4184.81 | 990.832 |
| left Dorsal part of the lateral geniculate complex, shell | 4045.48 | 1038.2 | 4790.78 | 1174.33 |
| left Dorsal part of the lateral geniculate complex, core | 3798.91 | 722.165 | 3980.04 | 799.694 |
| left Dorsal part of the lateral geniculate complex, ipsilateral zone | 2309.6 | 585.15 | 3765.73 | 1614.25 |
| left Thalamus, polymodal association cortex related | 5466.45 | 327.208 | 6321.2 | 881.218 |
| left Lateral group of the dorsal thalamus | 3701.29 | 548.41 | 5070.6 | 872.323 |
| left Lateral posterior nucleus of the thalamus | 3505.87 | 724.078 | 5642.25 | 1046.68 |
| left Posterior complex of the thalamus | 3206.41 | 722.086 | 4143.64 | 803.868 |
| left Posterior limiting nucleus of the thalamus | 6200.42 | 694.28 | 7133.32 | 920.334 |
| left Suprageniculate nucleus | 6749.87 | 1256.18 | 6064.07 | 875.519 |
| left Ethmoid nucleus of the thalamus | 2802.1 | 585.949 | 4516.47 | 1015.84 |
| left Anterior group of the dorsal thalamus | 6247.47 | 602.168 | 5978.59 | 1361.79 |
| left Anteroventral nucleus of thalamus | 9322.51 | 1434.83 | 7468.27 | 1711.53 |
| left Anteromedial nucleus | 11583.9 | 1282.24 | 8472.62 | 1955.94 |
| left Anteromedial nucleus, dorsal part | 9959.19 | 1230.19 | 7909.01 | 2112.22 |
| left Anteromedial nucleus, ventral part | 13975.4 | 1767.21 | 9302.23 | 1787.41 |
| left Anterodorsal nucleus | 4803.53 | 1101.62 | 3770.94 | 718.103 |
| left Interanteromedial nucleus of the thalamus | 9686.27 | 942.053 | 10588.2 | 2339.91 |
| left Interanterodorsal nucleus of the thalamus | 13827.1 | 832.546 | 14054.7 | 2077.8 |
| left Lateral dorsal nucleus of thalamus | 2077.05 | 354.131 | 3621.48 | 1311.04 |
| left Medial group of the dorsal thalamus | 5685.11 | 604.415 | 7611.88 | 1207.28 |
| left Intermediodorsal nucleus of the thalamus | 9245.62 | 1599.59 | 10825.4 | 2071.83 |
| left Mediodorsal nucleus of thalamus | 5273.08 | 519.375 | 7895.47 | 1228.62 |
| left Submedial nucleus of the thalamus | 5544.37 | 935.391 | 4579.8 | 1350.15 |
| left Perireunensis nucleus | 5720.41 | 618.603 | 7574.43 | 1904.58 |
| left Midline group of the dorsal thalamus | 13346.4 | 1122.14 | 13914.9 | 1215.55 |
| left Paraventricular nucleus of the thalamus | 17425 | 1837.01 | 19487.4 | 871.857 |
| left Parataenial nucleus | 13664.8 | 1409.59 | 13322.8 | 1932.85 |
| left Nucleus of reuniens | 8980.41 | 623.004 | 8484.36 | 1857.67 |
| left Xiphoid thalamic nucleus | 13251.1 | 2151.48 | 13921.2 | 2009 |
| left Intralaminar nuclei of the dorsal thalamus | 6090.95 | 496.691 | 6732.46 | 1119.85 |
| left Rhomboid nucleus | 10187.6 | 1794.71 | 9959.3 | 2316.61 |
| left Central medial nucleus of the thalamus | 8706.62 | 1331.09 | 7876.9 | 1376.32 |
| left Paracentral nucleus | 6202.85 | 1030.3 | 5669.38 | 1509.84 |
| left Central lateral nucleus of the thalamus | 5796.08 | 541.318 | 7520.51 | 1539.17 |
| left Parafascicular nucleus | 4055.21 | 535.275 | 5670.07 | 1002.47 |
| left Posterior intralaminar thalamic nucleus | 6489.87 | 1059.96 | 6268.04 | 1013.01 |
| left Reticular nucleus of the thalamus | 1100.67 | 215.361 | 1640.41 | 487.136 |
| left Geniculate group, ventral thalamus | 7085.36 | 1000.14 | 4466.26 | 677.915 |
| left Intergeniculate leaflet of the lateral geniculate complex | 8454.93 | 1769.19 | 5536.6 | 957.1 |
| left Intermediate geniculate nucleus | 10651.7 | 551.376 | 5639.16 | 1270.78 |
| left Ventral part of the lateral geniculate complex | 6944.26 | 987.689 | 4385.41 | 712.542 |
| left Subgeniculate nucleus | 2691.71 | 507.75 | 1940.53 | 559.168 |
| left Epithalamus | 3331.65 | 247.15 | 6640.28 | 1390.17 |
| left Medial habenula | 2375.69 | 322.789 | 5513.8 | 1390.93 |
| left Lateral habenula | 4217.14 | 312.71 | 7683.74 | 1429.99 |
| left Hypothalamus | 6187.21 | 421.443 | 7185.66 | 802.993 |
| left Periventricular zone | 7215.17 | 291.567 | 8148.25 | 1005.15 |
| left Supraoptic nucleus | 4000 | 828.88 | 6007.17 | 1555.16 |
| left Accessory supraoptic group | 13565.8 | 2986.75 | 14823.9 | 3613.92 |
| left Paraventricular hypothalamic nucleus | 13134.4 | 1069.1 | 13716.3 | 2213.3 |
| left Periventricular hypothalamic nucleus, anterior part | 15014.6 | 3040.74 | 12041.5 | 2980.35 |
| left Periventricular hypothalamic nucleus, intermediate part | 4808.19 | 155.284 | 5968.85 | 1124.94 |
| left Arcuate hypothalamic nucleus | 4021.43 | 369.508 | 5523.6 | 531.639 |
| left Periventricular region | 8166.01 | 687.459 | 8719.96 | 721.059 |
| left Anterodorsal preoptic nucleus | 9432.97 | 1183.19 | 10511.2 | 2589.63 |
| left Anteroventral preoptic nucleus | 5371.44 | 1118.43 | 5004.88 | 1241.04 |
| left Anteroventral periventricular nucleus | 4841.27 | 621.056 | 5910.33 | 712.361 |
| left Dorsomedial nucleus of the hypothalamus | 12736.1 | 616.004 | 15586.3 | 1653.52 |
| left Median preoptic nucleus | 4775.3 | 1374.22 | 5909.66 | 1181.53 |
| left Medial preoptic area | 5138.48 | 811.349 | 5804.92 | 1064.73 |
| left Vascular organ of the lamina terminalis | 3036.2 | 792.315 | 3955.06 | 1654.68 |
| left Posterodorsal preoptic nucleus | 5000.9 | 386.569 | 8036.13 | 1532.7 |
| left Parastrial nucleus | 6846.54 | 1763.92 | 8050.9 | 1640.31 |
| left Periventricular hypothalamic nucleus, posterior part | 4884.59 | 647.552 | 7052.87 | 767.42 |
| left Periventricular hypothalamic nucleus, preoptic part | 12251.9 | 1359.62 | 8249.83 | 1116.19 |
| left Subparaventricular zone | 14079.4 | 1276.85 | 12245.1 | 3112.64 |
| left Suprachiasmatic nucleus | 11106.1 | 2819.13 | 8000 | 1776.28 |
| left Subfornical organ | 3230.38 | 1222.67 | 5421.29 | 879.051 |
| left Ventromedial preoptic nucleus | 11914.5 | 1739.32 | 8210.74 | 1261.48 |
| left Ventrolateral preoptic nucleus | 7636.14 | 1052.52 | 8558.26 | 2395.26 |
| left Hypothalamic medial zone | 7241.09 | 490.073 | 8571.47 | 1195.24 |
| left Anterior hypothalamic nucleus | 8387.2 | 1015.76 | 8581.26 | 2032.8 |
| left Mammillary body | 4701.96 | 247.771 | 5332.67 | 1063.8 |
| left Lateral mammillary nucleus | 2462.86 | 311.167 | 1754.89 | 251.906 |
| left Medial mammillary nucleus | 2823.11 | 182.874 | 3418.43 | 718.661 |
| left Medial mammillary nucleus, median part | 3040 | 253.45 | 3248.89 | 809.216 |
| left Medial mammillary nucleus, lateral part | 2589.65 | 360.115 | 3446.61 | 818.907 |
| left Medial mammillary nucleus, medial part | 4216.32 | 378.025 | 4485.18 | 914.462 |
| left Medial mammillary nucleus, posterior part | 1573.47 | 248.743 | 1857.06 | 452.884 |
| left Medial mammillary nucleus, dorsal part | 1352.11 | 332.195 | 2333.06 | 681.393 |
| left Supramammillary nucleus | 7617.17 | 841.917 | 8273.64 | 1862.53 |
| left Tuberomammillary nucleus | 7800.16 | 832.56 | 9245.07 | 1737.01 |
| left Tuberomammillary nucleus, dorsal part | 8964.23 | 1465.94 | 11638.1 | 2689.19 |
| left Tuberomammillary nucleus, ventral part | 7427.91 | 807.916 | 8479.82 | 1609.73 |
| left Medial preoptic nucleus | 8207.36 | 1173.91 | 6980.19 | 1452.68 |
| left Dorsal premammillary nucleus | 4570.34 | 619.734 | 6326.61 | 1090.18 |
| left Ventral premammillary nucleus | 8297.05 | 1359.2 | 17053.7 | 2907.53 |
| left Paraventricular hypothalamic nucleus, descending division | 8580.26 | 1092.3 | 9980.55 | 1626.19 |
| left Ventromedial hypothalamic nucleus | 3401.23 | 345.566 | 4773.14 | 650.59 |
| left Posterior hypothalamic nucleus | 12126.9 | 999.084 | 14877 | 1921.25 |
| left Hypothalamic lateral zone | 4536.18 | 529.041 | 5542.59 | 749.621 |
| left Lateral hypothalamic area | 5166.35 | 631.835 | 6389.58 | 898.927 |
| left Lateral preoptic area | 3577.8 | 814.28 | 3887.37 | 730.921 |
| left Preparasubthalamic nucleus | 5604.52 | 1051.91 | 6947.54 | 1524.65 |
| left Parasubthalamic nucleus | 6890.06 | 1255.76 | 11946.9 | 3203 |
| left Perifornical nucleus | 7351.92 | 940.916 | 8147.52 | 1210.15 |
| left Retrochiasmatic area | 3099.46 | 631.054 | 2653.14 | 742.794 |
| left Subthalamic nucleus | 1868.04 | 391.314 | 2743.79 | 1042.94 |
| left Tuberal nucleus | 5661.85 | 514.078 | 9553.14 | 1318.51 |
| left Zona incerta | 3625.21 | 465.736 | 3539.88 | 514.3 |
| left Fields of Forel | 3127.93 | 608.976 | 3245.06 | 689.084 |
| left Median eminence | 1141.17 | 566.548 | 1257.44 | 315.252 |
| left Midbrain | 5418.12 | 537.084 | 6663.29 | 944.431 |
| left Midbrain, sensory related | 8688.11 | 1036.42 | 10751.8 | 1277.45 |
| left Superior colliculus, sensory related | 16371.6 | 2000.13 | 16915.8 | 1803.02 |
| left Superior colliculus, optic layer | 19993.8 | 2755.57 | 18194.2 | 2086.01 |
| left Superior colliculus, superficial gray layer | 16215.2 | 2131.1 | 18042.5 | 2179.52 |
| left Superior colliculus, zonal layer | 10438 | 1088.05 | 11059.4 | 1689.93 |
| left Inferior colliculus | 5281.16 | 801.01 | 8080.73 | 1213.42 |
| left Inferior colliculus, central nucleus | 5538.95 | 1090.08 | 9016.7 | 1555.7 |
| left Inferior colliculus, dorsal nucleus | 6241.31 | 936.48 | 8909.93 | 1326.57 |
| left Inferior colliculus, external nucleus | 4499.89 | 624.226 | 7003.65 | 987.033 |
| left Nucleus of the brachium of the inferior colliculus | 4980.57 | 1172.9 | 7558.57 | 1421.48 |
| left Nucleus sagulum | 2506.8 | 547.807 | 4302.72 | 1319.79 |
| left Parabigeminal nucleus | 1011.62 | 164.88 | 2945.61 | 871.976 |
| left Midbrain trigeminal nucleus | 3878.79 | 665.56 | 6580.09 | 1339.2 |
| left Subcommissural organ | 2555.04 | 561.757 | 3299.22 | 622.054 |
| left Midbrain, motor related | 5022.51 | 557.47 | 6085.23 | 974.375 |
| left Substantia nigra, reticular part | 1411.31 | 136.659 | 2140.87 | 255.152 |
| left Ventral tegmental area | 2216.28 | 230.426 | 3054.3 | 717.917 |
| left Paranigral nucleus | 2105.67 | 292.767 | 2136.41 | 536.589 |
| left Midbrain reticular nucleus, retrorubral area | 3328.66 | 730.031 | 4499.36 | 816.875 |
| left Midbrain reticular nucleus | 2660.15 | 395.202 | 3507.41 | 656.516 |
| left Superior colliculus, motor related | 8055.09 | 1230.99 | 8173.63 | 1411.06 |
| left Superior colliculus, motor related, deep gray layer | 4969.77 | 755.241 | 6321.75 | 1511.49 |
| left Superior colliculus, motor related, deep white layer | 4953.75 | 710.375 | 7056.07 | 2027.33 |
| left Superior colliculus, motor related, intermediate white layer | 7225.25 | 1064.1 | 7310.89 | 1531.54 |
| left Superior colliculus, motor related, intermediate gray layer | 11224.1 | 1883.36 | 10329.5 | 1492.78 |
| left Periaqueductal gray | 5849.47 | 486.792 | 8323.59 | 1362.82 |
| left Precommissural nucleus | 8902.17 | 547.569 | 12883.9 | 1334.92 |
| left Interstitial nucleus of Cajal | 1866.98 | 273.087 | 1643.62 | 246.318 |
| left Nucleus of Darkschewitsch | 3988.66 | 251.209 | 2994.22 | 714.729 |
| left Supraoculomotor periaqueductal gray | 9648.52 | 1452.32 | 10125.8 | 2302.49 |
| left Pretectal region | 6470.18 | 611.587 | 7178.6 | 931.512 |
| left Anterior pretectal nucleus | 4174.23 | 574.15 | 5252.91 | 933.145 |
| left Medial pretectal area | 15477.3 | 2179.84 | 17522.7 | 3611.1 |
| left Nucleus of the optic tract | 10008.8 | 1552.37 | 9986.13 | 2371.63 |
| left Nucleus of the posterior commissure | 10002.2 | 638.793 | 10295.1 | 1864.73 |
| left Olivary pretectal nucleus | 7666.31 | 769.948 | 9029.59 | 2199.14 |
| left Posterior pretectal nucleus | 12275.8 | 1786.48 | 10842.6 | 2649.97 |
| left Retroparafascicular nucleus | 4614.77 | 479.559 | 5461.08 | 896.016 |
| left Cuneiform nucleus | 1859.54 | 295.091 | 5231.4 | 1578.58 |
| left Red nucleus | 1296.2 | 246.079 | 2657.99 | 780.189 |
| left Oculomotor nucleus | 6218.79 | 2141.45 | 4159.59 | 1566.39 |
| left Medial accesory oculomotor nucleus | 7573.13 | 1620.39 | 3274.33 | 1643.63 |
| left Edinger-Westphal nucleus | 18620.2 | 1740.23 | 20263.6 | 3812.78 |
| left Trochlear nucleus | 4536.95 | 904.682 | 5260.94 | 1450.24 |
| left Paratrochlear nucleus | 4128.36 | 960.521 | 3122.46 | 1078.95 |
| left Ventral tegmental nucleus | 1872.91 | 500.556 | 1453.73 | 306.732 |
| left Anterior tegmental nucleus | 1513.88 | 354.065 | 2232.2 | 743.935 |
| left Lateral terminal nucleus of the accessory optic tract | 2975.44 | 790.74 | 2938.01 | 819.306 |
| left Dorsal terminal nucleus of the accessory optic tract | 5876.95 | 1935.68 | 5465.56 | 1187.81 |
| left Medial terminal nucleus of the accessory optic tract | 1531.54 | 440.967 | 2459.75 | 488.352 |
| left Midbrain, behavioral state related | 2833.73 | 375.968 | 4072.86 | 1006.91 |
| left Substantia nigra, compact part | 1725.56 | 389.917 | 2899.08 | 684.995 |
| left Pedunculopontine nucleus | 1970.66 | 419.172 | 4399.47 | 1294.62 |
| left Midbrain raphe nuclei | 4212.36 | 628.045 | 3989.29 | 858.41 |
| left Interfascicular nucleus raphe | 6646.61 | 1610.66 | 5272.66 | 824.396 |
| left Interpeduncular nucleus | 2383.78 | 471.366 | 2117.75 | 415.047 |
| left Interpeduncular nucleus, rostral | 2992.5 | 793.353 | 1770.63 | 574.056 |
| left Interpeduncular nucleus, caudal | 1000.63 | 356.745 | 568.889 | 128.432 |
| left Interpeduncular nucleus, apical | 7696.73 | 1758.85 | 8449.67 | 1432.17 |
| left Interpeduncular nucleus, lateral | 1946.24 | 400.432 | 1758.06 | 533.411 |
| left Interpeduncular nucleus, intermediate | 966.95 | 149.398 | 1017.31 | 240.651 |
| left Interpeduncular nucleus, dorsomedial | 4457.2 | 1223.38 | 3861.78 | 1045.78 |
| left Interpeduncular nucleus, dorsolateral | 1121.41 | 247.057 | 1827.81 | 433.256 |
| left Interpeduncular nucleus, rostrolateral | 1464.73 | 333.624 | 1688.75 | 389.664 |
| left Rostral linear nucleus raphe | 5905.55 | 1374.51 | 5823.36 | 1498.71 |
| left Central linear nucleus raphe | 4187.29 | 807.136 | 4717.79 | 1575.9 |
| left Dorsal nucleus raphe | 6610.52 | 431.854 | 6688.84 | 1490.02 |
| left Hindbrain | 1334.83 | 105.665 | 2211.08 | 321.932 |
| left Pons | 1799.41 | 269.825 | 2600.37 | 623.793 |
| left Pons, sensory related | 1171.06 | 121.388 | 2047.9 | 351.455 |
| left Nucleus of the lateral lemniscus | 443.753 | 109.636 | 719.774 | 158.265 |
| left Principal sensory nucleus of the trigeminal | 478.775 | 124.861 | 1391.89 | 368.919 |
| left Parabrachial nucleus | 2775.95 | 260.856 | 4502.78 | 832.688 |
| left Koelliker-Fuse subnucleus | 3509.32 | 602.723 | 4364.33 | 1220.8 |
| left Superior olivary complex | 598.484 | 95.493 | 854.713 | 170.148 |
| left Superior olivary complex, periolivary region | 1138.28 | 170.519 | 1136.45 | 277.91 |
| left Superior olivary complex, medial part | 161.487 | 42.9644 | 639.15 | 138.717 |
| left Superior olivary complex, lateral part | 271.512 | 70.9792 | 676.811 | 308.074 |
| left Pons, motor related | 2289.46 | 392.797 | 2927.56 | 777.769 |
| left Barrington's nucleus | 6517.31 | 916.037 | 14207.7 | 2368.85 |
| left Dorsal tegmental nucleus | 2051.03 | 330.222 | 3108.7 | 1026.64 |
| left Posterodorsal tegmental nucleus | 2052.36 | 555.665 | 3357.77 | 958.453 |
| left Pontine central gray | 3531.99 | 499.408 | 5215.91 | 937.558 |
| left Pontine gray | 5720.79 | 1115.66 | 4075.86 | 1077.8 |
| left Pontine reticular nucleus, caudal part | 558.594 | 137.002 | 1468.53 | 482.282 |
| left Supragenual nucleus | 3238.79 | 944.314 | 4286.06 | 1508.6 |
| left Supratrigeminal nucleus | 1592.19 | 485.122 | 3000.08 | 961.815 |
| left Tegmental reticular nucleus | 4788.2 | 1297.73 | 5100.19 | 2100.22 |
| left Motor nucleus of trigeminal | 341.814 | 72.2235 | 1726.91 | 780.867 |
| left Peritrigeminal zone | 477.643 | 140.575 | 2292.27 | 971.406 |
| left Accessory trigeminal nucleus | 92.1526 | 53.2043 | 1935.21 | 855.83 |
| left Parvicellular motor 5 nucleus | 723.084 | 318.981 | 3166.45 | 820.69 |
| left Intertrigeminal nucleus | 256.483 | 91.5651 | 1870.82 | 900.169 |
| left Pons, behavioral state related | 1992.14 | 419.756 | 3271.64 | 925.768 |
| left Superior central nucleus raphe | 2274.56 | 691.441 | 3087.99 | 1007.16 |
| left Locus ceruleus | 6618.34 | 1078.22 | 11303.5 | 3824.88 |
| left Laterodorsal tegmental nucleus | 7479.4 | 944.672 | 8737.66 | 705.346 |
| left Nucleus incertus | 6073.23 | 989.609 | 6647.81 | 1992.22 |
| left Pontine reticular nucleus | 1133.03 | 332.841 | 2553.41 | 908.429 |
| left Nucleus raphe pontis | 3789.07 | 758.126 | 3671.74 | 952.926 |
| left Subceruleus nucleus | 2454.73 | 578.922 | 4871.49 | 1426.67 |
| left Sublaterodorsal nucleus | 3372.5 | 476.632 | 6259.19 | 1411.21 |
| left Medulla | 1084.19 | 64.7141 | 2001.06 | 289.065 |
| left Medulla, sensory related | 1271.99 | 189.32 | 2509.27 | 591.237 |
| left Area postrema | 1442.83 | 532.457 | 5797.1 | 2109.89 |
| left Cochlear nuclei | 1004.02 | 283.734 | 2756.67 | 671.928 |
| left Dorsal cochlear nucleus | 1102.94 | 335.534 | 4478.76 | 1584.3 |
| left Ventral cochlear nucleus | 944.639 | 257.56 | 1722.85 | 456.664 |
| left Dorsal column nuclei | 1862.15 | 499.374 | 5850.42 | 1536.2 |
| left Cuneate nucleus | 1262.95 | 280.59 | 4609.51 | 1262.41 |
| left Gracile nucleus | 4314.06 | 1518.85 | 10928.1 | 2891.14 |
| left External cuneate nucleus | 555.589 | 111.892 | 732.663 | 242.272 |
| left Nucleus of the trapezoid body | 189.63 | 58.5927 | 1006.33 | 341.077 |
| left Nucleus of the solitary tract | 2405.38 | 353.692 | 4385.15 | 918.516 |
| left Spinal nucleus of the trigeminal, caudal part | 1644.32 | 279.942 | 2596.63 | 1035.11 |
| left Spinal nucleus of the trigeminal, interpolar part | 1242.7 | 381.828 | 1678.37 | 563.643 |
| left Spinal nucleus of the trigeminal, oral part | 275.381 | 37.8832 | 1110.55 | 334.723 |
| left Paratrigeminal nucleus | 867.686 | 292.84 | 815.494 | 561.054 |
| left Medulla, motor related | 951.592 | 86.5915 | 1917.86 | 244.526 |
| left Abducens nucleus | 3164.96 | 1146.34 | 3931.55 | 2164.58 |
| left Facial motor nucleus | 350.019 | 47.4203 | 868.323 | 238.542 |
| left Accessory facial motor nucleus | 180.791 | 112.743 | 3314.5 | 1151.07 |
| left Nucleus ambiguus | 625.181 | 143.465 | 1165.46 | 355.781 |
| left Nucleus ambiguus, dorsal division | 627.451 | 128.717 | 1050.98 | 237.29 |
| left Nucleus ambiguus, ventral division | 622.982 | 176.437 | 1276.35 | 603.977 |
| left Dorsal motor nucleus of the vagus nerve | 2976.48 | 283.655 | 5100.62 | 952.991 |
| left Gigantocellular reticular nucleus | 457.285 | 109.835 | 765.713 | 211.659 |
| left Infracerebellar nucleus | 522.273 | 185.845 | 866.359 | 255.381 |
| left Inferior olivary complex | 338.293 | 81.6404 | 556.42 | 152.313 |
| left Intermediate reticular nucleus | 787.646 | 116.217 | 2234.09 | 413.466 |
| left Inferior salivatory nucleus | 695.652 | 224.82 | 2643.48 | 1522.33 |
| left Linear nucleus of the medulla | 378.059 | 74.7063 | 1857.89 | 520.988 |
| left Lateral reticular nucleus | 1435.83 | 181.386 | 945.133 | 147.551 |
| left Lateral reticular nucleus, magnocellular part | 1349.91 | 221.399 | 989.195 | 134.052 |
| left Lateral reticular nucleus, parvicellular part | 2208.11 | 575.148 | 549.106 | 407.543 |
| left Magnocellular reticular nucleus | 409.485 | 138.976 | 842.798 | 212.241 |
| left Medullary reticular nucleus | 2575.75 | 300.177 | 5390.17 | 1174.07 |
| left Medullary reticular nucleus, dorsal part | 2278.59 | 173.782 | 4685.95 | 1460.17 |
| left Medullary reticular nucleus, ventral part | 2921.18 | 522.721 | 6208.8 | 1637.2 |
| left Parvicellular reticular nucleus | 724.735 | 123.437 | 2072.32 | 547.764 |
| left Parasolitary nucleus | 1890.75 | 457.088 | 1700.48 | 668.766 |
| left Paragigantocellular reticular nucleus | 886.663 | 190.287 | 1302.76 | 337.584 |
| left Paragigantocellular reticular nucleus, dorsal part | 1897.72 | 431.575 | 1991.35 | 682.426 |
| left Paragigantocellular reticular nucleus, lateral part | 546.972 | 131.158 | 1071.41 | 233.034 |
| left Perihypoglossal nuclei | 1103.7 | 330.362 | 1018.52 | 346.875 |
| left Nucleus of Roller | 1190.08 | 263.543 | 517.906 | 114.957 |
| left Nucleus prepositus | 1092.81 | 351.868 | 1081.68 | 394.354 |
| left Parapyramidal nucleus | 200.594 | 73.9449 | 503.244 | 256.318 |
| left Vestibular nuclei | 739.938 | 54.4432 | 1276.55 | 271.645 |
| left Lateral vestibular nucleus | 457.558 | 147.443 | 960.406 | 266.781 |
| left Medial vestibular nucleus | 780.09 | 96.2513 | 1359.29 | 368.544 |
| left Spinal vestibular nucleus | 977.511 | 78.1469 | 1327.99 | 270.78 |
| left Superior vestibular nucleus | 260.645 | 36.0133 | 999.294 | 256.931 |
| left Nucleus x | 1578.08 | 293.869 | 1619 | 501.316 |
| left Hypoglossal nucleus | 1654.42 | 336.704 | 1990.95 | 747.04 |
| left Nucleus y | 1068.09 | 426.643 | 1837.12 | 566.346 |
| left Medulla, behavioral state related | 1348.83 | 312.443 | 1307.51 | 275.105 |
| left Nucleus raphe magnus | 1786.03 | 483.858 | 1844.84 | 522.187 |
| left Nucleus raphe pallidus | 1055.13 | 432.991 | 730.895 | 270.722 |
| left Nucleus raphe obscurus | 470.294 | 176.415 | 620.388 | 111.125 |
| left Cerebellum | 1902.55 | 303.244 | 4662.15 | 974.681 |
| left Cerebellar cortex | 1975.71 | 316.74 | 4795.55 | 982.734 |
| left Vermal regions | 1625.52 | 209.514 | 4890.52 | 912.085 |
| left Lingula (I) | 525.959 | 143.459 | 3152.97 | 1198.91 |
| left Central lobule | 804.786 | 279.557 | 5992.66 | 1667.62 |
| left Lobule II | 955.731 | 249.139 | 5540.27 | 1713.42 |
| left Lobule III | 731.467 | 308.05 | 6212.4 | 1808.83 |
| left Culmen | 1368.2 | 233.11 | 5430.44 | 1513.72 |
| left Lobules IV-V | 1368.2 | 233.11 | 5430.44 | 1513.72 |
| left Declive (VI) | 2839.68 | 358.811 | 3130.93 | 625.78 |
| left Folium-tuber vermis (VII) | 2339.05 | 459.355 | 2373.61 | 477.704 |
| left Pyramus (VIII) | 1923.25 | 495.513 | 5634.51 | 2420.09 |
| left Uvula (IX) | 1853.25 | 379.125 | 5165.29 | 761.645 |
| left Nodulus (X) | 1317.87 | 269.018 | 4285.77 | 837.493 |
| left Hemispheric regions | 2204.14 | 403.805 | 4733.61 | 1076.18 |
| left Simple lobule | 1833.45 | 524.576 | 6456.75 | 1815.38 |
| left Ansiform lobule | 2591.32 | 600.032 | 5000.33 | 1253.34 |
| left Crus 1 | 3046.98 | 665.437 | 5795.69 | 1435.64 |
| left Crus 2 | 2083.52 | 538.407 | 4113.96 | 1143.47 |
| left Paramedian lobule | 1164.88 | 279.526 | 3649.3 | 1152.99 |
| left Copula pyramidis | 1257.8 | 466.329 | 4164.6 | 1209.09 |
| left Paraflocculus | 2950.93 | 548.163 | 3485.62 | 650.084 |
| left Flocculus | 3025.93 | 584.368 | 5609.96 | 1403.22 |
| left Cerebellar nuclei | 280.628 | 45.4712 | 2066.03 | 1068.94 |
| left Fastigial nucleus | 251.34 | 106.029 | 1047.92 | 534.278 |
| left Interposed nucleus | 352.886 | 64.3734 | 1731.54 | 1067.3 |
| left Dentate nucleus | 188.492 | 95.8322 | 4856.15 | 2288.41 |
| left Vestibulocerebellar nucleus | 85.9585 | 32.534 | 437.607 | 118.854 |
| left fiber tracts | 3748 | 307.285 | 4839.94 | 490.042 |
| left cranial nerves | 2859.99 | 203.474 | 2719.68 | 248.078 |
| left vomeronasal nerve | 3187.74 | 508.085 | 4291.19 | 1987.09 |
| left olfactory nerve | 4591.28 | 370.995 | 3897.25 | 422.769 |
| left olfactory nerve layer of main olfactory bulb | 3152.64 | 394.39 | 3032.05 | 478.496 |
| left lateral olfactory tract, general | 7530.82 | 611.369 | 4728.43 | 966.873 |
| left lateral olfactory tract, body | 3758.33 | 288.535 | 2458.39 | 660.69 |
| left dorsal limb | 29277 | 2871.26 | 17813.9 | 3106.59 |
| left anterior commissure, olfactory limb | 8312.1 | 1103.14 | 7444.84 | 1000.76 |
| left optic nerve | 1925.54 | 153.285 | 2622.62 | 499.879 |
| left brachium of the superior colliculus | 2877.4 | 513.775 | 3924.8 | 913.119 |
| left superior colliculus commissure | 4824.5 | 655.007 | 8995.68 | 2188.1 |
| left optic chiasm | 1941.09 | 526.94 | 3432.8 | 1341.25 |
| left optic tract | 1662.33 | 155.485 | 1771.5 | 283.58 |
| left oculomotor nerve | 3142.68 | 257.654 | 3144.3 | 499.255 |
| left medial longitudinal fascicle | 2125.16 | 306.202 | 2138.78 | 604.806 |
| left posterior commissure | 6325.35 | 485.523 | 6089.63 | 790.762 |
| left trochlear nerve | 3123.12 | 784.866 | 5285.29 | 1254.17 |
| left trigeminal nerve | 665.906 | 170.577 | 753.098 | 301.717 |
| left motor root of the trigeminal nerve | 1361.94 | 646.789 | 982.383 | 295.533 |
| left sensory root of the trigeminal nerve | 648.005 | 160.654 | 747.201 | 306.573 |
| left spinal tract of the trigeminal nerve | 397.816 | 66.5551 | 629.704 | 260.314 |
| left facial nerve | 1684.37 | 334.99 | 2399.61 | 975.227 |
| left genu of the facial nerve | 4766.45 | 1074.95 | 5472.16 | 2746.01 |
| left vestibulocochlear nerve | 941.642 | 88.3243 | 1678.35 | 254.018 |
| left vestibular nerve | 496.836 | 99.4893 | 764.569 | 138.716 |
| left cochlear nerve | 1020.47 | 116.351 | 1840.29 | 302.365 |
| left trapezoid body | 256.196 | 72.183 | 331.941 | 91.4028 |
| left dorsal acoustic stria | 773.748 | 272.629 | 801.382 | 189.112 |
| left lateral lemniscus | 759.934 | 28.6843 | 1657.45 | 458.59 |
| left inferior colliculus commissure | 5714.95 | 1186.85 | 5975.57 | 924.329 |
| left brachium of the inferior colliculus | 2482.61 | 595.426 | 4013.86 | 682.913 |
| left vagus nerve | 1955.56 | 485.998 | 6542.22 | 3015.17 |
| left solitary tract | 1955.56 | 485.998 | 6542.22 | 3015.17 |
| left dorsal roots | 1976.84 | 441.782 | 1850.71 | 425.543 |
| left cervicothalamic tract | 1976.84 | 441.782 | 1850.71 | 425.543 |
| left dorsal column | 521.415 | 286.883 | 804.469 | 654.816 |
| left cuneate fascicle | 521.415 | 286.883 | 804.469 | 654.816 |
| left medial lemniscus | 2025.82 | 459.797 | 1885.92 | 445.474 |
| left cerebellum related fiber tracts | 684.292 | 99.24 | 2865.62 | 801.516 |
| left cerebellar commissure | 203.267 | 88.1833 | 2758.62 | 1479.63 |
| left cerebellar peduncles | 760.933 | 104.321 | 1084.36 | 182.52 |
| left superior cerebelar peduncles | 1154.02 | 160.385 | 1573.59 | 283.821 |
| left superior cerebellar peduncle decussation | 656.07 | 146.476 | 1208.94 | 389.602 |
| left uncinate fascicle | 275.218 | 89.253 | 790.269 | 377.309 |
| left ventral spinocerebellar tract | 1087.84 | 202.837 | 907.702 | 134.723 |
| left middle cerebellar peduncle | 725.402 | 243.459 | 591.237 | 86.02 |
| left inferior cerebellar peduncle | 420.171 | 36.9505 | 1224.13 | 399.216 |
| left dorsal spinocerebellar tract | 311.585 | 116.477 | 377.541 | 208.463 |
| left arbor vitae | 657.849 | 109.263 | 3592.68 | 1084.89 |
| left supra-callosal cerebral white matter | 8045.18 | 813.804 | 10660.1 | 1483.8 |
| left lateral forebrain bundle system | 6086.31 | 685.262 | 7372.65 | 1061.21 |
| left corpus callosum | 5990.93 | 787.261 | 7787.24 | 1154.34 |
| left corpus callosum, anterior forceps | 8796.88 | 815.218 | 10453.5 | 793.841 |
| left external capsule | 12904.2 | 1327.22 | 13809.7 | 1506.41 |
| left corpus callosum, extreme capsule | 9781.94 | 1959.05 | 14137.8 | 1989.57 |
| left genu of corpus callosum | 3147.14 | 551.799 | 5776.54 | 990.353 |
| left corpus callosum, posterior forceps | 11518 | 2294.46 | 11963.6 | 2962.25 |
| left corpus callosum, body | 2673.23 | 393.292 | 4938.63 | 906.969 |
| left corpus callosum, splenium | 5551.66 | 907.659 | 6986.61 | 1623.48 |
| left corticospinal tract | 904.723 | 133.776 | 1293.4 | 283.565 |
| left internal capsule | 871.455 | 120.3 | 1297.5 | 271.725 |
| left cerebal peduncle | 1113.04 | 266.705 | 1634.6 | 520.989 |
| left pyramid | 443.131 | 116.209 | 281.344 | 74.7775 |
| left pyramidal decussation | 3381.87 | 1291.57 | 4950.05 | 1417.94 |
| left thalamus related | 14488.6 | 1602.59 | 15495.9 | 2443.78 |
| left external medullary lamina of the thalamus | 390.909 | 87.4861 | 1230.3 | 368.519 |
| left optic radiation | 16614.6 | 1917.98 | 17696 | 2784.13 |
| left auditory radiation | 8726.98 | 880.026 | 9446.46 | 1846.15 |
| left extrapyramidal fiber systems | 1202.78 | 178.223 | 1512.38 | 309.879 |
| left cerebral nuclei related | 3097.73 | 533.424 | 3716.09 | 1234.34 |
| left nigrostriatal tract | 3097.73 | 533.424 | 3716.09 | 1234.34 |
| left tectospinal pathway | 854.097 | 160.979 | 1222.87 | 339.408 |
| left doral tegmental decussation | 7471.58 | 1544.55 | 5898.62 | 1439.83 |
| left crossed tectospinal pathway | 764.241 | 161.424 | 1159.38 | 329.726 |
| left rubrospinal tract | 1143.5 | 208.886 | 1346.72 | 215.671 |
| left ventral tegmental decussation | 2716.98 | 586.441 | 2643.8 | 460.58 |
| left medial forebrain bundle system | 5167.59 | 541.177 | 6532.05 | 579.363 |
| left cerebrum related | 5502.11 | 618.53 | 6925.53 | 604.039 |
| left amygdalar capsule | 6890.22 | 1507.63 | 7403.19 | 1508.19 |
| left anterior commissure, temporal limb | 3015.28 | 597.573 | 4843.63 | 716.733 |
| left cingulum bundle | 12285.7 | 1402.82 | 13266.1 | 1000.26 |
| left fornix system | 3800.05 | 449.279 | 5348.88 | 602.372 |
| left alveus | 4677.45 | 444.062 | 6824.03 | 1146.06 |
| left dorsal fornix | 1465.65 | 389.346 | 4885.5 | 1822.69 |
| left fimbria | 1471.77 | 154.501 | 2822.49 | 374.361 |
| left postcommissural fornix | 3192.78 | 508.795 | 4143.87 | 409.584 |
| left medial corticohypothalamic tract | 7979.95 | 3718.49 | 6375.94 | 2302.71 |
| left columns of the fornix | 3044.6 | 495.568 | 4074.78 | 393.143 |
| left hippocampal commissures | 6309.75 | 1256.63 | 7426.26 | 884.701 |
| left dorsal hippocampal commissure | 6586.78 | 1334.56 | 7565.17 | 900.439 |
| left ventral hippocampal commissure | 2268.2 | 327.665 | 5399.74 | 1337.38 |
| left stria terminalis | 2448.57 | 220.714 | 4144.63 | 471.167 |
| left commissural branch of stria terminalis | 7823.48 | 626.013 | 9960.58 | 1394.16 |
| left hypothalamus related | 2630.64 | 243.62 | 3547.98 | 621.381 |
| left medial forebrain bundle | 4689.31 | 641.487 | 6238.99 | 1780.02 |
| left supraoptic commissures | 1935.94 | 374.394 | 1784.1 | 423.741 |
| left mammillary related | 2916.73 | 289.984 | 3186.51 | 677.864 |
| left principal mammillary tract | 5449.61 | 1249.83 | 6187.68 | 1263.3 |
| left mammillothalamic tract | 3125.2 | 304.666 | 3089.34 | 811.575 |
| left mammillotegmental tract | 2434.67 | 411.814 | 3048.33 | 791.912 |
| left mammillary peduncle | 468.293 | 96.225 | 780.488 | 306.506 |
| left epithalamus related | 2301.78 | 226.358 | 3563.37 | 606.568 |
| left stria medullaris | 2002.55 | 235.198 | 3380.85 | 739.176 |
| left fasciculus retroflexus | 2787.31 | 272.117 | 3906.64 | 877.185 |
| left habenular commissure | 2298.85 | 315.447 | 3395.23 | 686.738 |
| left ventricular systems | 2718.28 | 155.518 | 3778.27 | 223.292 |
| left lateral ventricle | 3037.63 | 217.592 | 3969.48 | 344.59 |
| left subependymal zone | 4320.33 | 804.106 | 5433.03 | 458.138 |
| left choroid plexus | 1417.65 | 130.708 | 2502.57 | 314.665 |
| left third ventricle | 4006.68 | 459.026 | 5231.99 | 548.228 |
| left cerebral aqueduct | 1124.28 | 202.414 | 3052.58 | 462.401 |
| left fourth ventricle | 977.63 | 174.677 | 2025.49 | 403.363 |
| left lateral recess | 975.978 | 240.564 | 2375.89 | 643.628 |
| left central canal, spinal cord/medulla | 0 | 0 | 0 | 0 |
| right root | 9067.16 | 749.136 | 9496.88 | 578.744 |
| right Basic cell groups and regions | 9694.06 | 804.646 | 10076.3 | 616 |
| right Cerebrum | 13419.7 | 1239.04 | 13482.8 | 724.637 |
| right Cerebral cortex | 14908.9 | 1438.31 | 14647.1 | 764.164 |
| right Cortical plate | 15134.4 | 1460.24 | 14772.6 | 752.403 |
| right Isocortex | 18878.9 | 2188.1 | 16848.5 | 982.989 |
| right Frontal pole, cerebral cortex | 14478.9 | 2144.54 | 11901.6 | 1120.23 |
| right Frontal pole, layer 1 | 2593.31 | 653.966 | 5760.29 | 1761.32 |
| right Frontal pole, layer 2/3 | 11881.8 | 2233.82 | 16360.4 | 1691.34 |
| right Frontal pole, layer 5 | 25763.1 | 4409.61 | 15122.5 | 3148.78 |
| right Frontal pole, layer 6a | 9452.79 | 1712.76 | 5808.2 | 259.939 |
| right Frontal pole, layer 6b | 6274.51 | 1640.2 | 12235.3 | 4408.93 |
| right Somatomotor areas | 13169.2 | 1305.55 | 10790 | 1885 |
| right Primary motor area | 11423.2 | 1150.69 | 9315.39 | 2027.29 |
| right Primary motor area, Layer 1 | 2108.23 | 536.392 | 1631.78 | 508.209 |
| right Primary motor area, Layer 2/3 | 11151.4 | 1192.16 | 9945.12 | 2594.38 |
| right Primary motor area, Layer 5 | 13235.5 | 1425.4 | 9971.92 | 2259.2 |
| right Primary motor area, Layer 6a | 13974.9 | 1323.46 | 11198.9 | 1895.51 |
| right Primary motor area, Layer 6b | 12114.1 | 1315.08 | 11139.4 | 1505.18 |
| right Secondary motor area | 14682.3 | 1747.99 | 12068 | 1883.04 |
| right Secondary motor area, layer 1 | 3375.27 | 507.405 | 2608.07 | 623.229 |
| right Secondary motor area, layer 2/3 | 15550.4 | 2361.74 | 14918.4 | 2791.01 |
| right Secondary motor area, layer 5 | 17905.3 | 2234.98 | 13773.7 | 2133.5 |
| right Secondary motor area, layer 6a | 18127.3 | 1501.13 | 13443.4 | 1740.51 |
| right Secondary motor area, layer 6b | 16815.7 | 1965.67 | 13566.4 | 1388.46 |
| right Somatosensory areas | 14811.7 | 2451.16 | 12002.5 | 1226.88 |
| right Primary somatosensory area | 16596.3 | 2775.71 | 13127.1 | 1782.98 |
| right Primary somatosensory area, nose | 12962 | 4193.22 | 9220.88 | 1352.01 |
| right Primary somatosensory area, nose, layer 1 | 3344.6 | 959.164 | 2472.24 | 1028.94 |
| right Primary somatosensory area, nose, layer 2/3 | 12636.5 | 3678.61 | 9915.47 | 1522.97 |
| right Primary somatosensory area, nose, layer 4 | 16880 | 6329.47 | 9819.81 | 1882.52 |
| right Primary somatosensory area, nose, layer 5 | 13802.3 | 5145.39 | 8907.8 | 1918.4 |
| right Primary somatosensory area, nose, layer 6a | 15084 | 4563.08 | 11794.5 | 2710.31 |
| right Primary somatosensory area, nose, layer 6b | 8725.58 | 1498.74 | 10449.6 | 2320.17 |
| right Primary somatosensory area, barrel field | 23666 | 4498.99 | 17664.8 | 2285.37 |
| right Primary somatosensory area, barrel field, layer 1 | 7515.86 | 1102.52 | 4132.61 | 1085.98 |
| right Primary somatosensory area, barrel field, layer 2/3 | 24588 | 5495.98 | 21456.4 | 2786.71 |
| right Primary somatosensory area, barrel field, layer 4 | 25558.6 | 5967.84 | 18864.7 | 3483.61 |
| right Primary somatosensory area, barrel field, layer 5 | 22280.1 | 4390.23 | 15862.1 | 3528.45 |
| right Primary somatosensory area, barrel field, layer 6a | 31978.9 | 5764.03 | 21924.7 | 3517.12 |
| right Primary somatosensory area, barrel field, layer 6b | 27877 | 4442.29 | 22590.7 | 3259.06 |
| right Primary somatosensory area, lower limb | 21973.9 | 4145.71 | 16869.5 | 3216.38 |
| right Primary somatosensory area, lower limb, layer 1 | 3049.52 | 631.453 | 2181.57 | 555.666 |
| right Primary somatosensory area, lower limb, layer 2/3 | 23598.2 | 5198.99 | 19650.4 | 3854.37 |
| right Primary somatosensory area, lower limb, layer 4 | 25224.2 | 6168.12 | 17853.2 | 4615.71 |
| right Primary somatosensory area, lower limb, layer 5 | 22598.5 | 4084.5 | 16168.1 | 3418.56 |
| right Primary somatosensory area, lower limb, layer 6a | 28922.3 | 5136.98 | 21943.7 | 3480.71 |
| right Primary somatosensory area, lower limb, layer 6b | 16500.6 | 3101.58 | 17902.4 | 3386.2 |
| right Primary somatosensory area, mouth | 5997.96 | 798.374 | 5316.33 | 1024.08 |
| right Primary somatosensory area, mouth, layer 1 | 1601.15 | 187.639 | 1386.98 | 472.075 |
| right Primary somatosensory area, mouth, layer 2/3 | 5729.82 | 835.792 | 5790.51 | 1310.1 |
| right Primary somatosensory area, mouth, layer 4 | 7541.19 | 1303.25 | 6118.22 | 1712.38 |
| right Primary somatosensory area, mouth, layer 5 | 6335.39 | 817.782 | 4853.51 | 1092.36 |
| right Primary somatosensory area, mouth, layer 6a | 7342.67 | 1146.63 | 6562.39 | 1023.21 |
| right Primary somatosensory area, mouth, layer 6b | 3891.73 | 438.564 | 6350.59 | 1429.54 |
| right Primary somatosensory area, upper limb | 17706.5 | 2436.94 | 15152.2 | 3167.47 |
| right Primary somatosensory area, upper limb, layer 1 | 3170.96 | 509.778 | 2949.71 | 838.996 |
| right Primary somatosensory area, upper limb, layer 2/3 | 18618.5 | 2549.28 | 19334.4 | 3760.19 |
| right Primary somatosensory area, upper limb, layer 4 | 20854.3 | 3599.02 | 16379.3 | 4007.31 |
| right Primary somatosensory area, upper limb, layer 5 | 18186.3 | 2651.03 | 13624 | 3573.78 |
| right Primary somatosensory area, upper limb, layer 6a | 21731.5 | 2855.29 | 17296.8 | 3258.88 |
| right Primary somatosensory area, upper limb, layer 6b | 17913.7 | 2186.76 | 16515.4 | 2660.37 |
| right Primary somatosensory area, trunk | 21968.2 | 3829.93 | 19934.5 | 3481.75 |
| right Primary somatosensory area, trunk, layer 1 | 3399.95 | 702.656 | 3411.37 | 1144.54 |
| right Primary somatosensory area, trunk, layer 2/3 | 22594.1 | 3890.85 | 22749.3 | 4351 |
| right Primary somatosensory area, trunk, layer 4 | 27404 | 5947.94 | 25609.6 | 4260.53 |
| right Primary somatosensory area, trunk, layer 5 | 22702.4 | 4146.98 | 20535.7 | 3582.33 |
| right Primary somatosensory area, trunk, layer 6a | 31806.4 | 5788.95 | 23947.3 | 4675.83 |
| right Primary somatosensory area, trunk, layer 6b | 26281.9 | 4382.6 | 24859.7 | 5064.73 |
| right Primary somatosensory area, unassigned | 22947.3 | 3988.46 | 17738.6 | 3224.98 |
| right Primary somatosensory area, unassigned, layer 1 | 7801.74 | 1795.95 | 4659.77 | 872.059 |
| right Primary somatosensory area, unassigned, layer 2/3 | 25943.5 | 5049.13 | 23820.2 | 3457.41 |
| right Primary somatosensory area, unassigned, layer 4 | 26838.9 | 6301.53 | 17660 | 4154.86 |
| right Primary somatosensory area, unassigned, layer 5 | 21332 | 3882.16 | 15825.3 | 4004.67 |
| right Primary somatosensory area, unassigned, layer 6a | 26931.8 | 3783.42 | 19712.1 | 3460.83 |
| right Primary somatosensory area, unassigned, layer 6b | 24218.8 | 3276.09 | 20977.8 | 3010.51 |
| right Supplemental somatosensory area | 10002.5 | 1612.91 | 8971.91 | 1084.69 |
| right Supplemental somatosensory area, layer 1 | 855.985 | 113.517 | 1216.61 | 513.145 |
| right Supplemental somatosensory area, layer 2/3 | 8183.79 | 1416.78 | 7606.86 | 1038.35 |
| right Supplemental somatosensory area, layer 4 | 12144.4 | 2329.56 | 10284.2 | 1187.25 |
| right Supplemental somatosensory area, layer 5 | 11669.6 | 2158.32 | 11257.2 | 2055.17 |
| right Supplemental somatosensory area, layer 6a | 13824.1 | 2040.55 | 11398.4 | 1703.37 |
| right Supplemental somatosensory area, layer 6b | 16046.9 | 2089.11 | 14042.1 | 2079.62 |
| right Gustatory areas | 5650.42 | 1189.48 | 6850.31 | 570.41 |
| right Gustatory areas, layer 1 | 679.309 | 101.346 | 1082.17 | 399.875 |
| right Gustatory areas, layer 2/3 | 4462.34 | 993.455 | 6215.93 | 965.115 |
| right Gustatory areas, layer 4 | 6983.13 | 1761.16 | 8268.19 | 1051.23 |
| right Gustatory areas, layer 5 | 6395.47 | 1499.46 | 7672.24 | 858.766 |
| right Gustatory areas, layer 6a | 7614.37 | 1510.77 | 8640.95 | 510.754 |
| right Gustatory areas, layer 6b | 5635.87 | 806.874 | 5705.02 | 822.678 |
| right Visceral area | 4473.04 | 683.587 | 4420.62 | 443.194 |
| right Visceral area, layer 1 | 236.66 | 28.0221 | 442.364 | 242.602 |
| right Visceral area, layer 2/3 | 2561.38 | 249.525 | 3125.79 | 889.217 |
| right Visceral area, layer 4 | 4961.07 | 739.061 | 4790.94 | 742.98 |
| right Visceral area, layer 5 | 5164.67 | 1092.49 | 4807.04 | 594.857 |
| right Visceral area, layer 6a | 7528.79 | 1111.46 | 6916.21 | 540.98 |
| right Visceral area, layer 6b | 8986.26 | 1402.46 | 11250.9 | 1128.63 |
| right Auditory areas | 17521.5 | 2594.15 | 11617 | 1562.66 |
| right Dorsal auditory area | 20025.2 | 3314.39 | 13985.6 | 2053.48 |
| right Dorsal auditory area, layer 1 | 2628.68 | 344.651 | 1153.11 | 524.115 |
| right Dorsal auditory area, layer 2/3 | 17227.2 | 2881.26 | 12282.8 | 2159.58 |
| right Dorsal auditory area, layer 4 | 25144.1 | 5033.7 | 19217.1 | 3081.5 |
| right Dorsal auditory area, layer 5 | 25057.6 | 4631.31 | 20360.9 | 4433.2 |
| right Dorsal auditory area, layer 6a | 28205.1 | 4406.49 | 14704 | 2864.53 |
| right Dorsal auditory area, layer 6b | 34614.1 | 5279.65 | 16648 | 3163.95 |
| right Primary auditory area | 18416.1 | 2797.99 | 11215.3 | 1822.07 |
| right Primary auditory area, layer 1 | 2304.52 | 413.171 | 1930 | 1098.86 |
| right Primary auditory area, layer 2/3 | 17420.5 | 2507.57 | 9623.05 | 2066.22 |
| right Primary auditory area, layer 4 | 23325.8 | 4041.2 | 14687.1 | 2172.68 |
| right Primary auditory area, layer 5 | 22565 | 3815.13 | 14309 | 2635.36 |
| right Primary auditory area, layer 6a | 24929.9 | 3169.49 | 14242 | 2962.12 |
| right Primary auditory area, layer 6b | 27466.2 | 3732.12 | 17246.8 | 3532.44 |
| right Posterior auditory area | 24942.2 | 3845.33 | 11420.5 | 2638.67 |
| right Posterior auditory area, layer 1 | 6123.24 | 1479.44 | 1505.57 | 441.572 |
| right Posterior auditory area, layer 2/3 | 27579.1 | 4044.06 | 9745.16 | 2605.49 |
| right Posterior auditory area, layer 4 | 37503.6 | 7604.48 | 18223 | 5110 |
| right Posterior auditory area, layer 5 | 27907.9 | 5270.2 | 14696.3 | 3411.19 |
| right Posterior auditory area, layer 6a | 26938 | 4023.59 | 13422.4 | 3259.63 |
| right Posterior auditory area, layer 6b | 30492.5 | 5275.3 | 17192.6 | 4037.88 |
| right Ventral auditory area | 12304.4 | 1679.07 | 10577.6 | 1251.23 |
| right Ventral auditory area, layer 1 | 856.153 | 142.37 | 1901.79 | 1306.97 |
| right Ventral auditory area, layer 2/3 | 7500.71 | 919.795 | 6267.67 | 1711.79 |
| right Ventral auditory area, layer 4 | 13051.5 | 1838.76 | 11639.9 | 1012.01 |
| right Ventral auditory area, layer 5 | 15705.6 | 2545.97 | 12932.6 | 1709.25 |
| right Ventral auditory area, layer 6a | 20444.5 | 2585.01 | 17365.1 | 2515.84 |
| right Ventral auditory area, layer 6b | 25897.6 | 2986.2 | 23250.6 | 3221.2 |
| right Visual areas | 36773.3 | 4524.45 | 34365.7 | 1616.91 |
| right Anterolateral visual area | 28891.7 | 5129.36 | 17857.9 | 1729.07 |
| right Anterolateral visual area, layer 1 | 7384.62 | 1898.28 | 663.272 | 121.541 |
| right Anterolateral visual area, layer 2/3 | 35423.2 | 5468.2 | 17407 | 3314.08 |
| right Anterolateral visual area, layer 4 | 37384.6 | 8600.2 | 27166.1 | 3530.92 |
| right Anterolateral visual area, layer 5 | 27073.8 | 6138.37 | 21673.1 | 3774.55 |
| right Anterolateral visual area, layer 6a | 33595.6 | 5964.39 | 19412.6 | 3253.49 |
| right Anterolateral visual area, layer 6b | 36991.9 | 7314.04 | 25961.1 | 4943.13 |
| right Anteromedial visual area | 45764.4 | 9727.95 | 40648.7 | 3938.35 |
| right Anteromedial visual area, layer 1 | 3849.18 | 1195.45 | 3479.07 | 799.483 |
| right Anteromedial visual area, layer 2/3 | 52293.2 | 12127.1 | 48164.5 | 3554.39 |
| right Anteromedial visual area, layer 4 | 61377.6 | 18152.3 | 55712.3 | 6174.04 |
| right Anteromedial visual area, layer 5 | 49220.6 | 11059.4 | 47643.1 | 6704.88 |
| right Anteromedial visual area, layer 6a | 62797.2 | 11049.4 | 45338.8 | 5413.31 |
| right Anteromedial visual area, layer 6b | 56297.8 | 8071.96 | 53901.9 | 5225 |
| right Lateral visual area | 37310.4 | 4392.99 | 27271.9 | 1714.09 |
| right Lateral visual area, layer 1 | 10406.9 | 3190.54 | 2011.05 | 629.311 |
| right Lateral visual area, layer 2/3 | 43744.5 | 5955.55 | 29547.3 | 3687.36 |
| right Lateral visual area, layer 4 | 46734.3 | 6681.61 | 40419.9 | 2725.07 |
| right Lateral visual area, layer 5 | 39719.6 | 5405.62 | 30496.8 | 3658.44 |
| right Lateral visual area, layer 6a | 41340.2 | 4581.49 | 29504.6 | 4131.94 |
| right Lateral visual area, layer 6b | 30685.3 | 7167.66 | 31176.1 | 5439.4 |
| right Primary visual area | 40290.5 | 4204.8 | 40310.2 | 1875.04 |
| right Primary visual area, layer 1 | 11787.4 | 2094.31 | 5219.25 | 862.667 |
| right Primary visual area, layer 2/3 | 47127.3 | 4536.63 | 51767.9 | 2666.63 |
| right Primary visual area, layer 4 | 45572.8 | 5460.4 | 51272.9 | 3182.18 |
| right Primary visual area, layer 5 | 43199.9 | 6018.85 | 40839.4 | 2750.64 |
| right Primary visual area, layer 6a | 50579.5 | 4946.93 | 47363.6 | 4311.57 |
| right Primary visual area, layer 6b | 48766.9 | 4734.58 | 51295.1 | 4699.79 |
| right Posterolateral visual area | 17680.8 | 2831.35 | 21568.8 | 1810.62 |
| right Posterolateral visual area, layer 1 | 2269.69 | 709.341 | 2234.37 | 807.108 |
| right Posterolateral visual area, layer 2/3 | 13624.3 | 2910.19 | 17312.5 | 2824.17 |
| right Posterolateral visual area, layer 4 | 37697 | 7852.59 | 40345 | 3320.12 |
| right Posterolateral visual area, layer 5 | 25581.9 | 4055.17 | 31798.3 | 2477.55 |
| right Posterolateral visual area, layer 6a | 27307.3 | 3484.64 | 33548.7 | 1893.75 |
| right Posterolateral visual area, layer 6b | 28049.4 | 6141.06 | 32948.1 | 5442.97 |
| right posteromedial visual area | 50015.9 | 10035.6 | 46810.2 | 3510.04 |
| right posteromedial visual area, layer 1 | 5675.23 | 2358.92 | 2837.61 | 763.774 |
| right posteromedial visual area, layer 2/3 | 55924.9 | 12331.7 | 54207.8 | 5085.32 |
| right posteromedial visual area, layer 4 | 62280.9 | 13976.3 | 64893.4 | 7825.06 |
| right posteromedial visual area, layer 5 | 55580.4 | 11980.5 | 57125 | 4076.49 |
| right posteromedial visual area, layer 6a | 70703.3 | 11216.2 | 52943.2 | 5044.66 |
| right posteromedial visual area, layer 6b | 75687.2 | 8502.59 | 63829.9 | 7857.79 |
| right Laterointermediate area | 29027.2 | 5021.79 | 16614.3 | 1538.38 |
| right Laterointermediate area, layer 1 | 5369.07 | 2461.06 | 1916.25 | 718.77 |
| right Laterointermediate area, layer 2/3 | 30314.4 | 5568.56 | 12278.1 | 1741.61 |
| right Laterointermediate area, layer 4 | 38311.3 | 8303.62 | 24054 | 1609.94 |
| right Laterointermediate area, layer 5 | 30692.2 | 5754.89 | 19446.8 | 2866.06 |
| right Laterointermediate area, layer 6a | 38733.1 | 5947.54 | 24765.2 | 4494.9 |
| right Laterointermediate area, layer 6b | 33297.8 | 7967.37 | 23523.4 | 4145.59 |
| right Postrhinal area | 19936.9 | 3700.05 | 18793.8 | 1965.49 |
| right Postrhinal area, layer 1 | 924.356 | 263.344 | 790.025 | 284.895 |
| right Postrhinal area, layer 2/3 | 13613.8 | 3039.73 | 10993 | 2531.91 |
| right Postrhinal area, layer 4 | 28798.9 | 7479.57 | 26848.5 | 3032.36 |
| right Postrhinal area, layer 5 | 28282.3 | 5820.1 | 28781.7 | 3266.44 |
| right Postrhinal area, layer 6a | 36304.8 | 5444.34 | 33648.3 | 4266.03 |
| right Postrhinal area, layer 6b | 33028 | 6785.8 | 31531.8 | 6332.84 |
| right Anterior cingulate area | 25215.8 | 3222.03 | 21129.4 | 1466.68 |
| right Anterior cingulate area, dorsal part | 29135.9 | 3699.48 | 22718.4 | 1521.51 |
| right Anterior cingulate area, dorsal part, layer 1 | 3134.63 | 251.737 | 3546.67 | 813.574 |
| right Anterior cingulate area, dorsal part, layer 2/3 | 33757 | 5525.43 | 29529.7 | 2564.96 |
| right Anterior cingulate area, dorsal part, layer 5 | 36349.9 | 5057.87 | 27168 | 2313.55 |
| right Anterior cingulate area, dorsal part, layer 6a | 35745.1 | 3588.58 | 26089.4 | 2278.1 |
| right Anterior cingulate area, dorsal part, layer 6b | 22523 | 3400.7 | 17964.9 | 2132.57 |
| right Anterior cingulate area, ventral part | 20127.8 | 2635.81 | 19067 | 1926.47 |
| right Anterior cingulate area, ventral part, layer 1 | 2201.01 | 418.021 | 3291.42 | 587.143 |
| right Anterior cingulate area, ventral part, layer 2/3 | 23053.7 | 3607.93 | 21519.2 | 2113.2 |
| right Anterior cingulate area, ventral part, layer 5 | 25585.8 | 3252.13 | 24236.7 | 2933.51 |
| right Anterior cingulate area, ventral part, 6a | 26056.1 | 3129.17 | 23492.9 | 2349.88 |
| right Anterior cingulate area, ventral part, 6b | 11385.6 | 2844.62 | 11154.9 | 2036.37 |
| right Prelimbic area | 24497.5 | 3082.8 | 19556.9 | 1046.54 |
| right Prelimbic area, layer 1 | 5610.62 | 1084.25 | 4937.57 | 732.975 |
| right Prelimbic area, layer 2/3 | 27204.8 | 4238.12 | 24350.7 | 1761.05 |
| right Prelimbic area, layer 5 | 33855.9 | 4192.28 | 25311.3 | 2027.42 |
| right Prelimbic area, layer 6a | 24622.8 | 2589.23 | 20302.4 | 1636.15 |
| right Prelimbic area, layer 6b | 5753.16 | 1433.16 | 6017.49 | 1687.16 |
| right Infralimbic area | 18435.6 | 1586.71 | 17797.5 | 1580.07 |
| right Infralimbic area, layer 1 | 4854.98 | 729.841 | 5162.34 | 835.681 |
| right Infralimbic area, layer 2/3 | 16891.4 | 2169.43 | 17800.3 | 2118.5 |
| right Infralimbic area, layer 5 | 24268.8 | 2244.33 | 23567.9 | 2482.46 |
| right Infralimbic area, layer 6a | 20889.4 | 1690.21 | 18640.2 | 1804.21 |
| right Infralimbic area, layer 6b | 5776.5 | 1094.97 | 3698.19 | 1564.07 |
| right Orbital area | 26801.8 | 2817.46 | 28109.2 | 1794.79 |
| right Orbital area, lateral part | 24011.3 | 3081.3 | 25871.8 | 2610.01 |
| right Orbital area, lateral part, layer 1 | 6011.96 | 1820.01 | 14300.5 | 4450.14 |
| right Orbital area, lateral part, layer 2/3 | 27008.2 | 5017 | 36840.5 | 4016.5 |
| right Orbital area, lateral part, layer 5 | 32789.1 | 4444.85 | 30908 | 2816.17 |
| right Orbital area, lateral part, layer 6a | 13717.5 | 1951.33 | 9667.41 | 1530.42 |
| right Orbital area, lateral part, layer 6b | 6231.39 | 909.857 | 7726.92 | 938.24 |
| right Orbital area, medial part | 24271.1 | 2630.76 | 24499.6 | 1544.47 |
| right Orbital area, medial part, layer 1 | 5947.92 | 857.481 | 9630.77 | 1805.18 |
| right Orbital area, medial part, layer 2/3 | 24206.9 | 4982.66 | 33867.1 | 3452.28 |
| right Orbital area, medial part, layer 5 | 39770.2 | 4468.15 | 34036.6 | 3029.98 |
| right Orbital area, medial part, layer 6a | 27925.4 | 1982.21 | 19445.3 | 988.718 |
| right Orbital area, medial part, layer 6b | 7024.39 | 1122.36 | 4509.49 | 1653.01 |
| right Orbital area, ventrolateral part | 33088.4 | 2911.44 | 34400.8 | 1758.75 |
| right Orbital area, ventrolateral part, layer 1 | 10965.3 | 3276.13 | 16326.7 | 5175.23 |
| right Orbital area, ventrolateral part, layer 2/3 | 25304.4 | 3316.25 | 36958.8 | 3550.73 |
| right Orbital area, ventrolateral part, layer 5 | 53610.7 | 6203.83 | 49248.6 | 5429.86 |
| right Orbital area, ventrolateral part, layer 6a | 33693.9 | 3452.58 | 20702.5 | 2845.98 |
| right Orbital area, ventrolateral part, layer 6b | 10261.6 | 1734.16 | 5873.42 | 1616.73 |
| right Agranular insular area | 8743.83 | 792.377 | 10918 | 902.619 |
| right Agranular insular area, dorsal part | 8979.41 | 848.703 | 10586.6 | 1302.03 |
| right Agranular insular area, dorsal part, layer 1 | 758.129 | 107.682 | 1480.05 | 669.031 |
| right Agranular insular area, dorsal part, layer 2/3 | 8221.43 | 954.228 | 9617.01 | 1301 |
| right Agranular insular area, dorsal part, layer 5 | 11310.5 | 1148.25 | 13680.6 | 1536.18 |
| right Agranular insular area, dorsal part, layer 6a | 10517.7 | 939.942 | 11444.6 | 1649.71 |
| right Agranular insular area, dorsal part, layer 6b | 3955.29 | 857.816 | 5175.54 | 955.87 |
| right Agranular insular area, posterior part | 5412.43 | 634.562 | 6365.6 | 438.408 |
| right Agranular insular area, posterior part, layer 1 | 382.329 | 109.793 | 561.693 | 109.956 |
| right Agranular insular area, posterior part, layer 2/3 | 4332.83 | 504.52 | 5914.63 | 787.653 |
| right Agranular insular area, posterior part, layer 5 | 6972.6 | 1190.46 | 7800.99 | 630.351 |
| right Agranular insular area, posterior part, layer 6a | 9405.73 | 911.739 | 10227.5 | 617.982 |
| right Agranular insular area, posterior part, layer 6b | 9481.48 | 1115.31 | 9193.19 | 1182.51 |
| right Agranular insular area, ventral part | 12916.9 | 1136.88 | 18023.9 | 1115.77 |
| right Agranular insular area, ventral part, layer 1 | 3422.77 | 724.752 | 2928.47 | 632.141 |
| right Agranular insular area, ventral part, layer 2/3 | 10321.8 | 788.972 | 16660.1 | 930.398 |
| right Agranular insular area, ventral part, layer 5 | 17744.3 | 1791.82 | 23527.2 | 1758.83 |
| right Agranular insular area, ventral part, layer 6a | 15748.4 | 1605.6 | 22034.2 | 1870.36 |
| right Agranular insular area, ventral part, layer 6b | 3591.84 | 958.194 | 4244.9 | 1529.63 |
| right Retrosplenial area | 27911.1 | 3545.62 | 26639.9 | 1363.26 |
| right Retrosplenial area, lateral agranular part | 31983.4 | 4289.51 | 32959.4 | 1681.3 |
| right Retrosplenial area, lateral agranular part, layer 1 | 3284.94 | 712.137 | 2739.32 | 552.789 |
| right Retrosplenial area, lateral agranular part, layer 2/3 | 35342.6 | 4726.08 | 40821.1 | 2667.49 |
| right Retrosplenial area, lateral agranular part, layer 5 | 40873.9 | 6094.93 | 41445.9 | 2218.09 |
| right Retrosplenial area, lateral agranular part, layer 6a | 44613.7 | 5869.71 | 41089.8 | 4078.72 |
| right Retrosplenial area, lateral agranular part, layer 6b | 47958.1 | 5953.05 | 45973.8 | 2952.02 |
| right Retrosplenial area, dorsal part | 29610.7 | 3881.36 | 28627.7 | 1803.27 |
| right Retrosplenial area, dorsal part, layer 1 | 7763.2 | 1002.73 | 6419.81 | 479.758 |
| right Retrosplenial area, dorsal part, layer 2/3 | 30051.8 | 4184.32 | 32597.3 | 2285.82 |
| right Retrosplenial area, dorsal part, layer 5 | 36178.3 | 5271.71 | 34201.7 | 2689.51 |
| right Retrosplenial area, dorsal part, layer 6a | 41920.1 | 5641.83 | 38445.3 | 3832.89 |
| right Retrosplenial area, dorsal part, layer 6b | 40260.4 | 5120.67 | 37625 | 4032.77 |
| right Retrosplenial area, ventral part | 24239.2 | 3180.84 | 21506 | 1576.44 |
| right Retrosplenial area, ventral part, layer 1 | 9034.65 | 1446.95 | 7556.33 | 1197.74 |
| right Retrosplenial area, ventral part, layer 2/3 | 24064.3 | 2651.56 | 18223.9 | 1402.84 |
| right Retrosplenial area, ventral part, layer 5 | 30674.1 | 4231.66 | 29049.3 | 2624.29 |
| right Retrosplenial area, ventral part, layer 6a | 30239.7 | 5033.42 | 27844.7 | 2219.87 |
| right Retrosplenial area, ventral part, layer 6b | 27509.9 | 3708.07 | 22103.7 | 2660.83 |
| right Posterior parietal association areas | 33808.1 | 5619.07 | 32736.4 | 3440.91 |
| right Anterior area | 34158.1 | 6906.26 | 35471.9 | 3938.19 |
| right Anterior area, layer 1 | 4913.67 | 1579.73 | 4102.14 | 841.018 |
| right Anterior area, layer 2/3 | 34595.6 | 6687.39 | 44449.8 | 5925.34 |
| right Anterior area, layer 4 | 45541.6 | 10851.6 | 47458.9 | 5402.42 |
| right Anterior area, layer 5 | 37131.4 | 8991.47 | 38106.4 | 4064.48 |
| right Anterior area, layer 6a | 49916 | 9010.32 | 37879.7 | 5092.61 |
| right Anterior area, layer 6b | 46804.5 | 8900.71 | 44871.2 | 5995.23 |
| right Rostrolateral visual area | 33312.8 | 4608.7 | 28864.9 | 3056.08 |
| right Rostrolateral area, layer 1 | 13539.5 | 4170.53 | 2584.19 | 475.506 |
| right Rostrolateral area, layer 2/3 | 37092.7 | 4026.88 | 35957.4 | 2747.32 |
| right Rostrolateral area, layer 4 | 35859.7 | 6892.2 | 35544.9 | 6007.78 |
| right Rostrolateral area, layer 5 | 32703 | 6579.3 | 30464.5 | 5365.36 |
| right Rostrolateral area, layer 6a | 45531.5 | 7576.52 | 34613.8 | 4909.84 |
| right Rostrolateral area, layer 6b | 41854.3 | 8038.66 | 38334.1 | 4671.98 |
| right Temporal association areas | 13437.9 | 1886.76 | 10680.7 | 1333.05 |
| right Temporal association areas, layer 1 | 1353.06 | 346.801 | 1065.78 | 438.231 |
| right Temporal association areas, layer 2/3 | 8769.56 | 1116.6 | 5883.7 | 1019.37 |
| right Temporal association areas, layer 4 | 17660.9 | 2940.8 | 13225.4 | 1650.01 |
| right Temporal association areas, layer 5 | 16614.3 | 2601.4 | 12974.2 | 1461.43 |
| right Temporal association areas, layer 6a | 21153.1 | 2839.18 | 18355.3 | 3184.52 |
| right Temporal association areas, layer 6b | 27533.2 | 3735.82 | 26743.4 | 4304.99 |
| right Perirhinal area | 8259.49 | 713.22 | 8886.5 | 1233.68 |
| right Perirhinal area, layer 1 | 3180.47 | 778.324 | 2575.02 | 815.927 |
| right Perirhinal area, layer 2/3 | 10027.2 | 893.034 | 9909.12 | 2247.13 |
| right Perirhinal area, layer 5 | 9793.65 | 1968.14 | 13132.9 | 2129.09 |
| right Perirhinal area, layer 6a | 11821 | 1794.81 | 13845.7 | 2314.98 |
| right Perirhinal area, layer 6b | 15015.2 | 2605.7 | 14912.9 | 3510.16 |
| right Ectorhinal area | 8485.99 | 1190.92 | 10185.6 | 1224.75 |
| right Ectorhinal area/Layer 1 | 1346.63 | 277.083 | 1168.9 | 417.936 |
| right Ectorhinal area/Layer 2/3 | 5579.8 | 580.674 | 6680 | 1121.07 |
| right Ectorhinal area/Layer 5 | 10972.4 | 2001.4 | 12370.7 | 1821.65 |
| right Ectorhinal area/Layer 6a | 12177.9 | 1828.79 | 15474.2 | 2909.86 |
| right Ectorhinal area/Layer 6b | 16468.9 | 2572.87 | 23361.6 | 4660.93 |
| right Olfactory areas | 9970.92 | 360.535 | 12184.7 | 584.122 |
| right Main olfactory bulb | 6669.29 | 835.703 | 7774.45 | 1409.82 |
| right Accessory olfactory bulb | 15767.8 | 1716.05 | 14172.8 | 1087.04 |
| right Accessory olfactory bulb, glomerular layer | 18602.9 | 4141.01 | 12040.5 | 3627.89 |
| right Accessory olfactory bulb, granular layer | 10612.2 | 2509.14 | 13599.4 | 2897.69 |
| right Accessory olfactory bulb, mitral layer | 18292.4 | 1085.25 | 15817.1 | 1985.3 |
| right Anterior olfactory nucleus | 19186 | 1605.46 | 23734.9 | 1661.33 |
| right Taenia tecta | 12464.3 | 1456.23 | 12553.7 | 1114.66 |
| right Taenia tecta, dorsal part | 17198.3 | 2045.04 | 17051 | 1538.84 |
| right Taenia tecta, ventral part | 7679.09 | 994.163 | 8007.86 | 949.539 |
| right Dorsal peduncular area | 17418.8 | 1244.66 | 16579.8 | 1587.79 |
| right Piriform area | 11909.1 | 909.876 | 15569.6 | 473.664 |
| right Nucleus of the lateral olfactory tract | 10839.3 | 1730.93 | 12397.9 | 2508.05 |
| right Nucleus of the lateral olfactory tract, molecular layer | 5547.27 | 828.372 | 4245.72 | 707.514 |
| right Nucleus of the lateral olfactory tract, pyramidal layer | 15557 | 2577.15 | 17873.8 | 3473.43 |
| right Nucleus of the lateral olfactory tract, layer 3 | 8853.39 | 1617.72 | 13959.3 | 3880.7 |
| right Cortical amygdalar area | 8947.57 | 1314.89 | 10978.3 | 1642.63 |
| right Cortical amygdalar area, anterior part | 10698.8 | 1665.46 | 14886.4 | 1996.08 |
| right Cortical amygdalar area, posterior part | 8408.85 | 1422.14 | 9776.09 | 1705.57 |
| right Cortical amygdalar area, posterior part, lateral zone | 6038.94 | 1204.69 | 8678.09 | 1534.63 |
| right Cortical amygdalar area, posterior part, medial zone | 10608.2 | 2224.56 | 10795.1 | 2118 |
| right Piriform-amygdalar area | 5750.93 | 909.352 | 9982.85 | 1592.46 |
| right Postpiriform transition area | 6113.49 | 820.27 | 8580.2 | 1693.93 |
| right Hippocampal formation | 9958.81 | 988.585 | 11602.6 | 762.368 |
| right Hippocampal region | 7578.59 | 680.114 | 8773.69 | 543.67 |
| right Ammon's horn | 8123.95 | 791.006 | 9623.08 | 688.44 |
| right Field CA1 | 10063.7 | 1095.4 | 11240.9 | 782.319 |
| right Field CA2 | 4929.72 | 501.924 | 6723.87 | 962.646 |
| right Field CA3 | 5204.59 | 398.733 | 7206.02 | 640.993 |
| right Dentate gyrus | 6303.67 | 643.886 | 6699.35 | 510.042 |
| right Dentate gyrus, molecular layer | 4657.28 | 622.236 | 4433.15 | 537.855 |
| right Dentate gyrus, polymorph layer | 9203.1 | 1107.11 | 10754.4 | 782.563 |
| right Dentate gyrus, granule cell layer | 9500.43 | 673.576 | 11072.2 | 816.946 |
| right Fasciola cinerea | 5761.71 | 1443.14 | 7726.91 | 2166.65 |
| right Induseum griseum | 1932.73 | 416.738 | 3703.09 | 983.258 |
| right Retrohippocampal region | 12935.2 | 1401.33 | 15173.8 | 1150.35 |
| right Entorhinal area | 10810.3 | 1025.82 | 13917.1 | 1669.99 |
| right Entorhinal area, lateral part | 10298.9 | 1087.18 | 11909.3 | 2212.13 |
| right Entorhinal area, lateral part, layer 1 | 840.729 | 275.913 | 360.604 | 85.1414 |
| right Entorhinal area, lateral part, layer 2 | 7035.47 | 1145.72 | 7448.76 | 2204.38 |
| right Entorhinal area, lateral part, layer 3 | 14267.2 | 1746.64 | 17210.4 | 3328.99 |
| right Entorhinal area, lateral part, layer 5 | 12109.9 | 1281.43 | 13902.5 | 2268.96 |
| right Entorhinal area, lateral part, layer 6a | 16680.3 | 2318.83 | 20152.6 | 4013.24 |
| right Entorhinal area, medial part, dorsal zone | 11459 | 1127.35 | 16463.6 | 1814.09 |
| right Entorhinal area, medial part, dorsal zone, layer 1 | 1797.31 | 504.486 | 2373.67 | 455.41 |
| right Entorhinal area, medial part, dorsal zone, layer 2 | 6290.01 | 1096.75 | 8739.81 | 1108.13 |
| right Entorhinal area, medial part, dorsal zone, layer 3 | 13387 | 2102.57 | 20131.1 | 2463.55 |
| right Entorhinal area, medial part, dorsal zone, layer 5 | 19277.4 | 1424.48 | 27244.3 | 2959.65 |
| right Entorhinal area, medial part, dorsal zone, layer 6 | 26602.2 | 2904.38 | 38482.9 | 6184.32 |
| right Parasubiculum | 9481.52 | 2153.47 | 12764.1 | 1524.6 |
| right Postsubiculum | 20812.6 | 2794.46 | 21285.8 | 933.437 |
| right Presubiculum | 15840.6 | 3246.14 | 18235.6 | 1382.18 |
| right Subiculum | 18532.4 | 2594.62 | 18851.1 | 1913.15 |
| right Prosubiculum | 15705 | 2368.14 | 14811.5 | 1134.83 |
| right Hippocampo-amygdalar transition area | 10026.8 | 2073.07 | 10620.8 | 2354.83 |
| right Area prostriata | 19494.3 | 3100.04 | 20765.4 | 1708.56 |
| right Cortical subplate | 9524.83 | 977.522 | 11649.7 | 1326.58 |
| right Claustrum | 19009 | 1746.96 | 19079.3 | 2046.41 |
| right Endopiriform nucleus | 11647.1 | 1218.83 | 14848 | 1333.53 |
| right Endopiriform nucleus, dorsal part | 14072.2 | 1427.53 | 18220.6 | 1715.12 |
| right Endopiriform nucleus, ventral part | 7119.33 | 1149.08 | 8551.08 | 916.929 |
| right Lateral amygdalar nucleus | 7948.93 | 1154.89 | 8420.99 | 1377.71 |
| right Basolateral amygdalar nucleus | 6471.18 | 908.5 | 8116.35 | 1491.92 |
| right Basolateral amygdalar nucleus, anterior part | 7104.04 | 1075.58 | 9174.27 | 1979.97 |
| right Basolateral amygdalar nucleus, posterior part | 6258.29 | 684.101 | 7916.41 | 1523.62 |
| right Basolateral amygdalar nucleus, ventral part | 5668.23 | 1463.71 | 6509.27 | 873.954 |
| right Basomedial amygdalar nucleus | 8130.33 | 937.166 | 10493 | 1663.7 |
| right Basomedial amygdalar nucleus, anterior part | 9482.25 | 1162.28 | 11459.7 | 2032.85 |
| right Basomedial amygdalar nucleus, posterior part | 6683.04 | 756.353 | 9458.16 | 1461.67 |
| right Posterior amygdalar nucleus | 7874.87 | 1029.91 | 10379.1 | 1799.89 |
| right Cerebral nuclei | 7350.63 | 726.319 | 8738.01 | 609.266 |
| right Striatum | 7873.03 | 721.88 | 9372.68 | 732.606 |
| right Striatum dorsal region | 6831.48 | 756.9 | 8852.38 | 806.832 |
| right Caudoputamen | 6831.48 | 756.9 | 8852.38 | 806.832 |
| right Striatum ventral region | 8461.27 | 949.685 | 9144.65 | 723.115 |
| right Nucleus accumbens | 10009.5 | 1034.84 | 10427.8 | 1238.1 |
| right Fundus of striatum | 6473.4 | 1433.07 | 6917.69 | 653.503 |
| right Olfactory tubercle | 6903.42 | 895.822 | 7918.93 | 702.457 |
| right Lateral septal complex | 13779.6 | 1201.73 | 13772.9 | 698.343 |
| right Lateral septal nucleus | 14886.6 | 1252.02 | 14727.1 | 790.648 |
| right Lateral septal nucleus, caudal (caudodorsal) part | 7616.34 | 1140.37 | 6907.99 | 841.777 |
| right Lateral septal nucleus, rostral (rostroventral) part | 15755.2 | 1477.8 | 15636.5 | 767.352 |
| right Lateral septal nucleus, ventral part | 18918.9 | 1301.04 | 19140.9 | 1657.9 |
| right Septofimbrial nucleus | 7043.46 | 1007.82 | 8157.92 | 446.636 |
| right Septohippocampal nucleus | 10017.5 | 2599.27 | 8033.81 | 1086.85 |
| right Striatum-like amygdalar nuclei | 7362.2 | 768.153 | 9341.34 | 1550.73 |
| right Anterior amygdalar area | 6878.07 | 1543.72 | 10954.9 | 1820.2 |
| right Bed nucleus of the accessory olfactory tract | 6937.34 | 1065.11 | 5761.07 | 1159.36 |
| right Central amygdalar nucleus | 3174.75 | 255.6 | 5050.7 | 1002.6 |
| right Central amygdalar nucleus, capsular part | 3805.22 | 403.414 | 6075.87 | 1175.71 |
| right Central amygdalar nucleus, lateral part | 2340.15 | 241.186 | 4296.69 | 1024.74 |
| right Central amygdalar nucleus, medial part | 3203.57 | 309.865 | 4888.03 | 1003.68 |
| right Intercalated amygdalar nucleus | 6985.04 | 453.428 | 10363.4 | 2362.41 |
| right Medial amygdalar nucleus | 10238.4 | 1124.3 | 11685.8 | 2306.08 |
| right Pallidum | 4848.96 | 857.213 | 5698.68 | 408.067 |
| right Pallidum, dorsal region | 1182.71 | 194.757 | 1641.81 | 224.475 |
| right Globus pallidus, external segment | 1276.8 | 241.262 | 1721.63 | 220.634 |
| right Globus pallidus, internal segment | 834.748 | 70.3978 | 1346.62 | 337.786 |
| right Pallidum, ventral region | 5769.97 | 1320.33 | 6134.69 | 639.227 |
| right Substantia innominata | 5977.47 | 1391.81 | 6153.71 | 638.605 |
| right Magnocellular nucleus | 4046.07 | 786.557 | 5976.74 | 779.252 |
| right Pallidum, medial region | 5848.77 | 721.322 | 7619.7 | 754.842 |
| right Medial septal complex | 5327.5 | 699.63 | 7178.14 | 903.5 |
| right Medial septal nucleus | 7537.57 | 1085.32 | 8201.41 | 965.897 |
| right Diagonal band nucleus | 4057.05 | 731.551 | 6589.91 | 1214.03 |
| right Triangular nucleus of septum | 7544.38 | 1136.22 | 9056.03 | 1197.46 |
| right Pallidum, caudal region | 8586.99 | 1584.14 | 10339.3 | 359.716 |
| right Bed nuclei of the stria terminalis | 8588.88 | 1590.08 | 10345.4 | 364.137 |
| right Bed nucleus of the anterior commissure | 8252.4 | 2428.72 | 9262 | 812.763 |
| right Brain stem | 4473.83 | 255.325 | 5031.35 | 542.32 |
| right Interbrain | 5694.35 | 210.808 | 7331.53 | 705.123 |
| right Thalamus | 5084.01 | 283.519 | 6183.46 | 788.027 |
| right Thalamus, sensory-motor cortex related | 3758.82 | 270.984 | 4361.24 | 720.969 |
| right Ventral group of the dorsal thalamus | 3042.36 | 278.583 | 3519.97 | 740.369 |
| right Ventral anterior-lateral complex of the thalamus | 2645.44 | 377.572 | 3573.38 | 980.894 |
| right Ventral medial nucleus of the thalamus | 2949.12 | 474.985 | 3248.92 | 583.817 |
| right Ventral posterior complex of the thalamus | 3163.52 | 380.913 | 3395.17 | 896.285 |
| right Ventral posterolateral nucleus of the thalamus | 1576.22 | 338.142 | 2905.08 | 1199.25 |
| right Ventral posterolateral nucleus of the thalamus, parvicellular part | 3617.36 | 774.325 | 2221.66 | 937.518 |
| right Ventral posteromedial nucleus of the thalamus | 3801.91 | 530.783 | 3604.18 | 933.928 |
| right Ventral posteromedial nucleus of the thalamus, parvicellular part | 4538.38 | 432.368 | 4333.43 | 690.26 |
| right Posterior triangular thalamic nucleus | 3281.59 | 406.59 | 5523.64 | 1314.45 |
| right Subparafascicular nucleus | 10208.9 | 323.964 | 10477 | 929.442 |
| right Subparafascicular nucleus, magnocellular part | 19468.7 | 912.822 | 19157.9 | 2707.47 |
| right Subparafascicular nucleus, parvicellular part | 5396.69 | 687.763 | 5965.68 | 575.793 |
| right Subparafascicular area | 11712.8 | 758.846 | 13435.6 | 1488.29 |
| right Peripeduncular nucleus | 2407.56 | 313.617 | 4604.59 | 1749.36 |
| right Geniculate group, dorsal thalamus | 4589.08 | 698.062 | 5490.81 | 1115.29 |
| right Medial geniculate complex | 3282.71 | 755.556 | 3426.04 | 680.134 |
| right Medial geniculate complex, dorsal part | 2940.39 | 1047.47 | 2890.1 | 810.222 |
| right Medial geniculate complex, ventral part | 3215.63 | 816.952 | 2857.27 | 472.835 |
| right Medial geniculate complex, medial part | 3592.37 | 607.3 | 4423.93 | 1047.63 |
| right Dorsal part of the lateral geniculate complex | 5856.92 | 887.405 | 7494.67 | 1681.39 |
| right Dorsal part of the lateral geniculate complex, shell | 4974.24 | 743.206 | 5151.37 | 1268.08 |
| right Dorsal part of the lateral geniculate complex, core | 6319.04 | 976.782 | 8380.5 | 1782.14 |
| right Dorsal part of the lateral geniculate complex, ipsilateral zone | 5746.53 | 1659.72 | 8687.5 | 2412.65 |
| right Thalamus, polymodal association cortex related | 5739 | 470.644 | 7141.39 | 873.485 |
| right Lateral group of the dorsal thalamus | 4631.91 | 829.018 | 5092.06 | 1222.59 |
| right Lateral posterior nucleus of the thalamus | 4684.39 | 939.683 | 5261.81 | 1342.81 |
| right Posterior complex of the thalamus | 4868.91 | 1099.54 | 4786.38 | 1291.32 |
| right Posterior limiting nucleus of the thalamus | 5022.37 | 433.341 | 6117.22 | 879.105 |
| right Suprageniculate nucleus | 3772.58 | 449.028 | 5019.72 | 1066.37 |
| right Ethmoid nucleus of the thalamus | 3433.74 | 932.417 | 5009.36 | 1275.27 |
| right Anterior group of the dorsal thalamus | 6438.57 | 970.275 | 7846.55 | 1155.27 |
| right Anteroventral nucleus of thalamus | 8333.48 | 1470.35 | 11251.2 | 1561.07 |
| right Anteromedial nucleus | 12004.5 | 1949.03 | 10395.7 | 740.655 |
| right Anteromedial nucleus, dorsal part | 12509.5 | 2008.06 | 10647.6 | 916.747 |
| right Anteromedial nucleus, ventral part | 11259.1 | 2099.6 | 10024 | 686.174 |
| right Anterodorsal nucleus | 4280.02 | 1372.8 | 6284.45 | 1577.14 |
| right Interanteromedial nucleus of the thalamus | 11600.6 | 2443 | 12310.7 | 1950.59 |
| right Interanterodorsal nucleus of the thalamus | 12017.3 | 1894.46 | 15161 | 1594.07 |
| right Lateral dorsal nucleus of thalamus | 2908.76 | 569.324 | 4661.05 | 1665.65 |
| right Medial group of the dorsal thalamus | 6146.37 | 746.482 | 7993.73 | 943.619 |
| right Intermediodorsal nucleus of the thalamus | 8296.5 | 2157.53 | 10633.7 | 1624.24 |
| right Mediodorsal nucleus of thalamus | 6007.22 | 757.094 | 8645.96 | 1131.94 |
| right Submedial nucleus of the thalamus | 4935.4 | 1081.17 | 2601.97 | 437.006 |
| right Perireunensis nucleus | 7288.63 | 486.384 | 9841.41 | 1838.22 |
| right Midline group of the dorsal thalamus | 13039.3 | 1653.95 | 16387.2 | 710.665 |
| right Paraventricular nucleus of the thalamus | 18561.2 | 2685.79 | 24327.9 | 1099.09 |
| right Parataenial nucleus | 11798 | 1611.98 | 16690.5 | 1037.21 |
| right Nucleus of reuniens | 8288.34 | 918.381 | 8800.43 | 945.311 |
| right Xiphoid thalamic nucleus | 10987 | 1405.79 | 11729.7 | 1910.08 |
| right Intralaminar nuclei of the dorsal thalamus | 6194.3 | 767.134 | 7303.86 | 818.323 |
| right Rhomboid nucleus | 13144.5 | 2571.91 | 10677.4 | 2576.14 |
| right Central medial nucleus of the thalamus | 9914.77 | 1633.69 | 9690.78 | 1156 |
| right Paracentral nucleus | 4896.34 | 889.122 | 4989.88 | 695.281 |
| right Central lateral nucleus of the thalamus | 5715.51 | 1353.96 | 8140.13 | 1345.85 |
| right Parafascicular nucleus | 4427.28 | 995.712 | 6153.72 | 1329.27 |
| right Posterior intralaminar thalamic nucleus | 4795.63 | 334.808 | 6699.44 | 938.024 |
| right Reticular nucleus of the thalamus | 883.132 | 150.729 | 2179.62 | 725.987 |
| right Geniculate group, ventral thalamus | 5961.61 | 704.195 | 7319.9 | 1534.27 |
| right Intergeniculate leaflet of the lateral geniculate complex | 8384.74 | 993.155 | 11477.2 | 2246.32 |
| right Intermediate geniculate nucleus | 9487.95 | 1002.86 | 6766.71 | 1555.74 |
| right Ventral part of the lateral geniculate complex | 5604.59 | 752.724 | 6949.76 | 1569.65 |
| right Subgeniculate nucleus | 2162.66 | 572.247 | 2749.14 | 746.173 |
| right Epithalamus | 3441.04 | 241.085 | 5293.52 | 861.255 |
| right Medial habenula | 2885.22 | 234.658 | 3905.31 | 511.833 |
| right Lateral habenula | 3955.4 | 345.445 | 6578.2 | 1204.85 |
| right Hypothalamus | 6507.72 | 518.093 | 8861.52 | 936.746 |
| right Periventricular zone | 8980.79 | 1029.86 | 9714.92 | 837.045 |
| right Supraoptic nucleus | 4181.9 | 1621.36 | 6647.24 | 1361.23 |
| right Accessory supraoptic group | 10390.3 | 981.423 | 11329.9 | 2506.52 |
| right Paraventricular hypothalamic nucleus | 14586.1 | 1586.93 | 14926.7 | 977.001 |
| right Periventricular hypothalamic nucleus, anterior part | 15372.9 | 2967.03 | 14600.7 | 3828.38 |
| right Periventricular hypothalamic nucleus, intermediate part | 9307.84 | 1583.12 | 8207.6 | 2129.33 |
| right Arcuate hypothalamic nucleus | 4661.11 | 678.003 | 6950.7 | 718.619 |
| right Periventricular region | 8076.54 | 1023.41 | 11169.2 | 933.021 |
| right Anterodorsal preoptic nucleus | 8003.38 | 1428.59 | 12097.9 | 872.711 |
| right Anteroventral preoptic nucleus | 4733.13 | 1192.57 | 10619.8 | 1715.65 |
| right Anteroventral periventricular nucleus | 4181.1 | 865.183 | 8513.2 | 1063.16 |
| right Dorsomedial nucleus of the hypothalamus | 15635.8 | 1491.91 | 20486.6 | 3613.71 |
| right Median preoptic nucleus | 5717.75 | 1138.11 | 6658.27 | 602.696 |
| right Medial preoptic area | 5158.04 | 918.607 | 8954.51 | 1346.41 |
| right Vascular organ of the lamina terminalis | 3494.88 | 1127.97 | 4841.87 | 637.694 |
| right Posterodorsal preoptic nucleus | 7016.04 | 1244.49 | 10153.3 | 1397.6 |
| right Parastrial nucleus | 8427.43 | 2398.45 | 12089.1 | 1503.8 |
| right Periventricular hypothalamic nucleus, posterior part | 3907.9 | 534.827 | 5699.13 | 1156.33 |
| right Periventricular hypothalamic nucleus, preoptic part | 8978.1 | 1130.93 | 8167.13 | 1267.36 |
| right Subparaventricular zone | 14000.8 | 1727.66 | 13646 | 2358.91 |
| right Suprachiasmatic nucleus | 10535.7 | 1836.57 | 6565.27 | 1284.18 |
| right Subfornical organ | 1685.78 | 440.241 | 8384.15 | 2408.45 |
| right Ventromedial preoptic nucleus | 5323.35 | 926.184 | 7100.02 | 1847.38 |
| right Ventrolateral preoptic nucleus | 4810.17 | 1475.23 | 7577.25 | 1502.05 |
| right Hypothalamic medial zone | 7173.39 | 606.512 | 10129.6 | 1378.87 |
| right Anterior hypothalamic nucleus | 7984.77 | 974.868 | 10049.1 | 1522.04 |
| right Mammillary body | 3905.84 | 281.595 | 5542.62 | 1078.88 |
| right Lateral mammillary nucleus | 1081.81 | 226.151 | 1380.39 | 269.572 |
| right Medial mammillary nucleus | 2682.57 | 242.408 | 3228.88 | 697.43 |
| right Medial mammillary nucleus, median part | 3847.81 | 1019.68 | 2875.47 | 866.591 |
| right Medial mammillary nucleus, lateral part | 2247.83 | 457.41 | 3548.8 | 618.742 |
| right Medial mammillary nucleus, medial part | 3612.82 | 653.153 | 3941.71 | 1042.18 |
| right Medial mammillary nucleus, posterior part | 2400.44 | 1715.54 | 1492.64 | 516.812 |
| right Medial mammillary nucleus, dorsal part | 1173.64 | 333.07 | 2312.63 | 629.08 |
| right Supramammillary nucleus | 7295.03 | 944.572 | 9265.18 | 1810.65 |
| right Tuberomammillary nucleus | 4016.53 | 520.512 | 9912.55 | 2073.41 |
| right Tuberomammillary nucleus, dorsal part | 7017.05 | 1239.53 | 14128.2 | 4433.93 |
| right Tuberomammillary nucleus, ventral part | 3033.98 | 399.746 | 8532.1 | 1801.24 |
| right Medial preoptic nucleus | 5894.8 | 647.795 | 8313.5 | 1232.36 |
| right Dorsal premammillary nucleus | 4524.1 | 627.586 | 7470.21 | 1856.72 |
| right Ventral premammillary nucleus | 6236.13 | 1344.48 | 22300.9 | 3841.58 |
| right Paraventricular hypothalamic nucleus, descending division | 11025.9 | 1341.33 | 13066.5 | 1816.06 |
| right Ventromedial hypothalamic nucleus | 3954.65 | 429.651 | 6919.13 | 1515.14 |
| right Posterior hypothalamic nucleus | 14332.1 | 1214.19 | 16940.1 | 2290.48 |
| right Hypothalamic lateral zone | 4974.12 | 477.679 | 6897.59 | 846.623 |
| right Lateral hypothalamic area | 5748.17 | 694.555 | 7546.12 | 904.636 |
| right Lateral preoptic area | 5283.52 | 848 | 8255.95 | 1771.93 |
| right Preparasubthalamic nucleus | 3070.31 | 467.021 | 8575.68 | 3365.73 |
| right Parasubthalamic nucleus | 8072.4 | 1350.55 | 12321.8 | 2474.97 |
| right Perifornical nucleus | 6883.62 | 705.776 | 9673.99 | 1195.54 |
| right Retrochiasmatic area | 1865.59 | 577.437 | 2017.1 | 308.63 |
| right Subthalamic nucleus | 1771.22 | 466.022 | 3041.51 | 878.407 |
| right Tuberal nucleus | 3972.13 | 710.247 | 10686.4 | 1176.75 |
| right Zona incerta | 4381.19 | 752.835 | 4606.13 | 944.409 |
| right Fields of Forel | 4240.06 | 1033.67 | 3793.89 | 960.718 |
| right Median eminence | 791.795 | 228.378 | 804.103 | 298.709 |
| right Midbrain | 6944.24 | 772.474 | 6777.29 | 710.503 |
| right Midbrain, sensory related | 10976.1 | 1785.73 | 9604.69 | 1152.95 |
| right Superior colliculus, sensory related | 19987.5 | 2884.23 | 15535.5 | 2209.02 |
| right Superior colliculus, optic layer | 25345.2 | 3680.97 | 19332.1 | 2581.71 |
| right Superior colliculus, superficial gray layer | 19385 | 3359.39 | 15119.9 | 2316.21 |
| right Superior colliculus, zonal layer | 12438.1 | 1726.83 | 10149.9 | 1388.76 |
| right Inferior colliculus | 7039.97 | 1362.85 | 7020.11 | 1129.4 |
| right Inferior colliculus, central nucleus | 8211.46 | 1793.85 | 7770.63 | 1471.7 |
| right Inferior colliculus, dorsal nucleus | 8041.53 | 1770.99 | 7748.87 | 1162.79 |
| right Inferior colliculus, external nucleus | 5717.22 | 878.181 | 6115.04 | 1125.19 |
| right Nucleus of the brachium of the inferior colliculus | 2558.48 | 553.266 | 4331.2 | 535.902 |
| right Nucleus sagulum | 4202.33 | 1333.78 | 5031.96 | 866.517 |
| right Parabigeminal nucleus | 992.948 | 172.194 | 1985.9 | 655.523 |
| right Midbrain trigeminal nucleus | 7123.04 | 826.244 | 9843.4 | 2566.93 |
| right Subcommissural organ | 4378.95 | 1253.02 | 4042.11 | 949.021 |
| right Midbrain, motor related | 6830.51 | 682.448 | 6622.86 | 692.548 |
| right Substantia nigra, reticular part | 878.824 | 92.4935 | 2063.58 | 449.942 |
| right Ventral tegmental area | 2059.74 | 297.675 | 3386.64 | 1015.48 |
| right Paranigral nucleus | 1797.75 | 422.674 | 3535.58 | 932.376 |
| right Midbrain reticular nucleus, retrorubral area | 5334.54 | 1069.1 | 4568.33 | 408.614 |
| right Midbrain reticular nucleus | 3620.45 | 542.352 | 3895.45 | 667.47 |
| right Superior colliculus, motor related | 11470.1 | 1317.63 | 8315.69 | 1228.8 |
| right Superior colliculus, motor related, deep gray layer | 7917.1 | 794.719 | 5745.24 | 1083.96 |
| right Superior colliculus, motor related, deep white layer | 7043.25 | 881.569 | 6004.43 | 780.719 |
| right Superior colliculus, motor related, intermediate white layer | 10986.1 | 1296.29 | 7959.32 | 1309.47 |
| right Superior colliculus, motor related, intermediate gray layer | 14775.8 | 1905.52 | 10567.7 | 1418.55 |
| right Periaqueductal gray | 8641.18 | 752.489 | 9497.99 | 1170.15 |
| right Precommissural nucleus | 11011.3 | 768.909 | 16670.7 | 1652.02 |
| right Interstitial nucleus of Cajal | 3110.24 | 522.217 | 3632.09 | 989.3 |
| right Nucleus of Darkschewitsch | 5129.6 | 755.57 | 4438.38 | 1104.88 |
| right Supraoculomotor periaqueductal gray | 13053.8 | 2185.96 | 11951.4 | 2462.67 |
| right Pretectal region | 6506.21 | 964.608 | 8420.45 | 1039.55 |
| right Anterior pretectal nucleus | 4353.85 | 672.835 | 5649.03 | 1020.45 |
| right Medial pretectal area | 10349.6 | 1840.51 | 15709.4 | 2308.43 |
| right Nucleus of the optic tract | 9174.37 | 2156.96 | 12704 | 1942.02 |
| right Nucleus of the posterior commissure | 11334.1 | 1625.02 | 12638 | 1365.52 |
| right Olivary pretectal nucleus | 7460.37 | 1792.71 | 11302.2 | 1861.49 |
| right Posterior pretectal nucleus | 11251 | 1619.27 | 16353.4 | 1886.74 |
| right Retroparafascicular nucleus | 4713.93 | 1001.08 | 5180.49 | 875.834 |
| right Cuneiform nucleus | 2816.18 | 412.951 | 5516.26 | 1040.52 |
| right Red nucleus | 2534.27 | 471.061 | 2971.3 | 725.831 |
| right Oculomotor nucleus | 5180.66 | 1536.35 | 2880.41 | 596.166 |
| right Medial accesory oculomotor nucleus | 1320.24 | 251.037 | 4296.01 | 1947.55 |
| right Edinger-Westphal nucleus | 18109.3 | 1624.22 | 15978.1 | 3082.61 |
| right Trochlear nucleus | 5447.78 | 1038.74 | 6317.6 | 1291.6 |
| right Paratrochlear nucleus | 3328.17 | 590.807 | 2439.28 | 577.826 |
| right Ventral tegmental nucleus | 3989.09 | 1684.62 | 2959.08 | 561.47 |
| right Anterior tegmental nucleus | 3031.01 | 1167.4 | 2465.12 | 472.169 |
| right Lateral terminal nucleus of the accessory optic tract | 3124.21 | 780.564 | 2828.42 | 358.227 |
| right Dorsal terminal nucleus of the accessory optic tract | 4007.37 | 984.944 | 3624.31 | 795.853 |
| right Medial terminal nucleus of the accessory optic tract | 1614.75 | 384.051 | 2561.86 | 892.399 |
| right Midbrain, behavioral state related | 3032.51 | 369.256 | 3951.25 | 624.03 |
| right Substantia nigra, compact part | 1230.12 | 179.693 | 3031.49 | 631.282 |
| right Pedunculopontine nucleus | 2784.89 | 651.249 | 3427.01 | 598.329 |
| right Midbrain raphe nuclei | 3796.35 | 296.884 | 4809.97 | 758.342 |
| right Interfascicular nucleus raphe | 5220.42 | 905.078 | 7478.71 | 1356.82 |
| right Interpeduncular nucleus | 2126.89 | 365.873 | 2715.9 | 476.255 |
| right Interpeduncular nucleus, rostral | 2516.26 | 437.268 | 2353.66 | 940.752 |
| right Interpeduncular nucleus, caudal | 1508.32 | 442.601 | 1145.38 | 303.555 |
| right Interpeduncular nucleus, apical | 4557.35 | 762.206 | 9040.8 | 1474.94 |
| right Interpeduncular nucleus, lateral | 1909.56 | 257.006 | 1635.23 | 345.49 |
| right Interpeduncular nucleus, intermediate | 2031.26 | 439.133 | 1622.97 | 477.623 |
| right Interpeduncular nucleus, dorsomedial | 2507.67 | 696.231 | 6232.82 | 1693.84 |
| right Interpeduncular nucleus, dorsolateral | 1218.3 | 232.362 | 2717.06 | 633.207 |
| right Interpeduncular nucleus, rostrolateral | 1181.27 | 328.348 | 1727.82 | 381.957 |
| right Rostral linear nucleus raphe | 4481.26 | 694.892 | 5720.64 | 977.506 |
| right Central linear nucleus raphe | 3993.1 | 437.152 | 5282.09 | 985.885 |
| right Dorsal nucleus raphe | 6402.47 | 422.176 | 7460.29 | 1504.87 |
| right Hindbrain | 1660.78 | 126.274 | 1973.1 | 336.662 |
| right Pons | 2277.9 | 307.292 | 2287.08 | 422.574 |
| right Pons, sensory related | 1646.8 | 152.897 | 2109.77 | 396.752 |
| right Nucleus of the lateral lemniscus | 748.691 | 160.02 | 935.519 | 121.643 |
| right Principal sensory nucleus of the trigeminal | 308.977 | 64.7933 | 1073.56 | 374.048 |
| right Parabrachial nucleus | 4578.26 | 438.784 | 5005.57 | 965.403 |
| right Koelliker-Fuse subnucleus | 3600.57 | 536.813 | 3851.64 | 966.41 |
| right Superior olivary complex | 353.368 | 88.0753 | 713.41 | 128.159 |
| right Superior olivary complex, periolivary region | 435.263 | 108.354 | 932.446 | 186.645 |
| right Superior olivary complex, medial part | 386.263 | 141.283 | 712.706 | 197.221 |
| right Superior olivary complex, lateral part | 245.653 | 58.7271 | 476.508 | 146.495 |
| right Pons, motor related | 2742.06 | 422.297 | 2320.72 | 450.314 |
| right Barrington's nucleus | 14080.8 | 2408.3 | 17452.6 | 3309.97 |
| right Dorsal tegmental nucleus | 3631.62 | 738.475 | 4775.74 | 1363.94 |
| right Posterodorsal tegmental nucleus | 2732.72 | 470.146 | 3664.49 | 902.21 |
| right Pontine central gray | 5214.58 | 608.893 | 5592.87 | 1157.58 |
| right Pontine gray | 4834.44 | 1279.38 | 2840.23 | 645.105 |
| right Pontine reticular nucleus, caudal part | 961.515 | 152.958 | 1100.35 | 226.659 |
| right Supragenual nucleus | 5685.09 | 1727.51 | 5894.24 | 2320.39 |
| right Supratrigeminal nucleus | 2129.22 | 602.857 | 1917.33 | 463.718 |
| right Tegmental reticular nucleus | 6195.67 | 1354.8 | 3952.44 | 1277.24 |
| right Motor nucleus of trigeminal | 625.901 | 158.583 | 1083.33 | 208.116 |
| right Peritrigeminal zone | 566.096 | 135 | 799.378 | 136.013 |
| right Accessory trigeminal nucleus | 402.069 | 143.54 | 614.93 | 311.382 |
| right Parvicellular motor 5 nucleus | 789.238 | 156.135 | 1937.22 | 414.229 |
| right Intertrigeminal nucleus | 197.39 | 63.2155 | 485.884 | 163.065 |
| right Pons, behavioral state related | 2761.52 | 423.485 | 2933.05 | 553.421 |
| right Superior central nucleus raphe | 2256.26 | 360.608 | 2803.1 | 605.716 |
| right Locus ceruleus | 12721.5 | 2904.58 | 10435.8 | 3478.36 |
| right Laterodorsal tegmental nucleus | 10814 | 893.662 | 10735.4 | 1352.77 |
| right Nucleus incertus | 6910.32 | 789.112 | 7348.97 | 1659.35 |
| right Pontine reticular nucleus | 1885.16 | 460.259 | 1975.02 | 447.173 |
| right Nucleus raphe pontis | 2915.72 | 444.408 | 3158.69 | 735.047 |
| right Subceruleus nucleus | 3985.9 | 885.42 | 4186.45 | 1251.63 |
| right Sublaterodorsal nucleus | 4835.56 | 550.702 | 5347.56 | 879.864 |
| right Medulla | 1327.84 | 96.4069 | 1803.7 | 325.072 |
| right Medulla, sensory related | 1679.47 | 128.619 | 2339.84 | 366.454 |
| right Area postrema | 1562.32 | 570.331 | 5183.23 | 1599.07 |
| right Cochlear nuclei | 883.816 | 101.764 | 1041.85 | 261.747 |
| right Dorsal cochlear nucleus | 1127.86 | 136.799 | 1608.82 | 499 |
| right Ventral cochlear nucleus | 737.103 | 119.029 | 701.003 | 136.6 |
| right Dorsal column nuclei | 2186.51 | 469.459 | 3482.99 | 818.085 |
| right Cuneate nucleus | 1524.24 | 375.331 | 3128.54 | 835.454 |
| right Gracile nucleus | 4879.56 | 1877.52 | 4924.33 | 1301.27 |
| right External cuneate nucleus | 936.015 | 119.237 | 1085.52 | 481.67 |
| right Nucleus of the trapezoid body | 295.517 | 143.972 | 1046.17 | 347.995 |
| right Nucleus of the solitary tract | 3862.59 | 408.278 | 4318.44 | 735.758 |
| right Spinal nucleus of the trigeminal, caudal part | 2809.55 | 391.78 | 4048.64 | 803.827 |
| right Spinal nucleus of the trigeminal, interpolar part | 1190 | 189.359 | 1664.59 | 538.213 |
| right Spinal nucleus of the trigeminal, oral part | 420.864 | 53.1057 | 1179.42 | 410.145 |
| right Paratrigeminal nucleus | 353.072 | 111.026 | 320.68 | 119.17 |
| right Medulla, motor related | 1245.39 | 119.419 | 1725.58 | 342.965 |
| right Abducens nucleus | 3599.32 | 933.116 | 2286.5 | 674.649 |
| right Facial motor nucleus | 577.274 | 89.0172 | 774.905 | 132.654 |
| right Accessory facial motor nucleus | 545.455 | 306.046 | 1030.3 | 432.106 |
| right Nucleus ambiguus | 2162.68 | 500.405 | 3429.4 | 898.629 |
| right Nucleus ambiguus, dorsal division | 2749.14 | 531.068 | 4382.92 | 1187.02 |
| right Nucleus ambiguus, ventral division | 1595.44 | 513.818 | 2507.12 | 926.689 |
| right Dorsal motor nucleus of the vagus nerve | 3933.45 | 750.823 | 3362.78 | 579.683 |
| right Gigantocellular reticular nucleus | 766.321 | 136.406 | 660.863 | 157.151 |
| right Infracerebellar nucleus | 792.877 | 212 | 991.096 | 471.069 |
| right Inferior olivary complex | 244.955 | 76.8715 | 525.932 | 107.678 |
| right Intermediate reticular nucleus | 1260.54 | 133.224 | 1696.37 | 469.741 |
| right Inferior salivatory nucleus | 973.913 | 182.586 | 2875.36 | 1335.57 |
| right Linear nucleus of the medulla | 1454.3 | 379.785 | 1913.84 | 1092.64 |
| right Lateral reticular nucleus | 1303.27 | 331.403 | 1758.56 | 311.395 |
| right Lateral reticular nucleus, magnocellular part | 1170.07 | 317.658 | 1619.75 | 340.868 |
| right Lateral reticular nucleus, parvicellular part | 2492.37 | 675.967 | 2997.82 | 1105.5 |
| right Magnocellular reticular nucleus | 667.876 | 134.782 | 1033.02 | 223.641 |
| right Medullary reticular nucleus | 2571.89 | 285.954 | 5044.37 | 1489.91 |
| right Medullary reticular nucleus, dorsal part | 2681.77 | 372.378 | 5275.99 | 1407.77 |
| right Medullary reticular nucleus, ventral part | 2444.04 | 237.676 | 4774.85 | 1851.92 |
| right Parvicellular reticular nucleus | 1032.72 | 78.6099 | 1585.33 | 448.765 |
| right Parasolitary nucleus | 1803.66 | 653.241 | 1982.83 | 1066.59 |
| right Paragigantocellular reticular nucleus | 1600.58 | 335.913 | 1634.65 | 421.072 |
| right Paragigantocellular reticular nucleus, dorsal part | 2422.47 | 267.789 | 2594.1 | 709.508 |
| right Paragigantocellular reticular nucleus, lateral part | 1324.12 | 377.259 | 1311.92 | 345.961 |
| right Perihypoglossal nuclei | 1249.21 | 311.722 | 1234.49 | 278.751 |
| right Nucleus of Roller | 1039.32 | 394.883 | 1422.22 | 461.998 |
| right Nucleus prepositus | 1275.69 | 307.609 | 1210.8 | 295.378 |
| right Parapyramidal nucleus | 331.241 | 87.554 | 595.529 | 149.953 |
| right Vestibular nuclei | 991.112 | 99.5328 | 1197.27 | 269.197 |
| right Lateral vestibular nucleus | 523.035 | 103.314 | 747.523 | 264.969 |
| right Medial vestibular nucleus | 1092.45 | 112.302 | 1368.77 | 287.671 |
| right Spinal vestibular nucleus | 1053.54 | 125.091 | 1133.63 | 301.694 |
| right Superior vestibular nucleus | 719.715 | 38.9248 | 817.687 | 180.746 |
| right Nucleus x | 3953.87 | 1291.64 | 2073.59 | 538.07 |
| right Hypoglossal nucleus | 1981.36 | 394.144 | 1375.38 | 562.422 |
| right Nucleus y | 2070.92 | 623.683 | 2411.35 | 524.503 |
| right Medulla, behavioral state related | 919.241 | 209.259 | 1210.09 | 282.512 |
| right Nucleus raphe magnus | 1208.26 | 394.664 | 1623.68 | 503.91 |
| right Nucleus raphe pallidus | 708.142 | 206.102 | 766.04 | 346.914 |
| right Nucleus raphe obscurus | 407.007 | 102.695 | 721.862 | 143.12 |
| right Cerebellum | 2255.72 | 204.491 | 3880.9 | 652.895 |
| right Cerebellar cortex | 2335.59 | 211.577 | 3986.76 | 662.506 |
| right Vermal regions | 2149.97 | 276.636 | 5166.14 | 876.456 |
| right Lingula (I) | 479.156 | 161.646 | 3814.19 | 1542.09 |
| right Central lobule | 1347.18 | 351.335 | 6317.96 | 1857.9 |
| right Lobule II | 1234.55 | 272.196 | 7206.5 | 2260.04 |
| right Lobule III | 1402.14 | 394.672 | 5884.41 | 1671.46 |
| right Culmen | 1953.37 | 403.398 | 4567.48 | 1047.44 |
| right Lobules IV-V | 1953.37 | 403.398 | 4567.48 | 1047.44 |
| right Declive (VI) | 2495.75 | 418.746 | 3635.05 | 1363.55 |
| right Folium-tuber vermis (VII) | 2883.43 | 401.517 | 2795.31 | 566.804 |
| right Pyramus (VIII) | 3468.59 | 721.528 | 6246.56 | 1009.24 |
| right Uvula (IX) | 2958.94 | 487.614 | 6728.63 | 1114.58 |
| right Nodulus (X) | 1790.17 | 695.291 | 6688.64 | 1516.58 |
| right Hemispheric regions | 2457.58 | 212.079 | 3211.66 | 553.751 |
| right Simple lobule | 2572.13 | 628.581 | 4736.2 | 1241.94 |
| right Ansiform lobule | 2729.63 | 331.694 | 3136.18 | 534.611 |
| right Crus 1 | 3743.63 | 502.16 | 3237.38 | 701.447 |
| right Crus 2 | 1600.48 | 238.64 | 3023.5 | 361.007 |
| right Paramedian lobule | 1776.22 | 291.622 | 3362.63 | 387.236 |
| right Copula pyramidis | 1401.81 | 360.375 | 3463.95 | 638.14 |
| right Paraflocculus | 2249.08 | 613.936 | 1717.08 | 437.187 |
| right Flocculus | 5132.85 | 450.72 | 2706.15 | 619.814 |
| right Cerebellar nuclei | 402.808 | 48.4896 | 1644.13 | 595.278 |
| right Fastigial nucleus | 433.458 | 112.912 | 2124.01 | 919.035 |
| right Interposed nucleus | 454.65 | 57.7389 | 799.797 | 249.861 |
| right Dentate nucleus | 293.487 | 58.3505 | 3390.96 | 2001.2 |
| right Vestibulocerebellar nucleus | 129.364 | 22.6894 | 466.495 | 176.834 |
| right fiber tracts | 4344.7 | 386.749 | 5252.84 | 338.408 |
| right cranial nerves | 1714.65 | 74.3157 | 2068.11 | 211.28 |
| right vomeronasal nerve | 7119.44 | 1195.28 | 4396.57 | 1348.79 |
| right olfactory nerve | 2294.57 | 117.201 | 2765.66 | 327.345 |
| right olfactory nerve layer of main olfactory bulb | 776.001 | 96.3573 | 1525.12 | 473.785 |
| right lateral olfactory tract, general | 3402.27 | 510.229 | 2641.89 | 197.394 |
| right lateral olfactory tract, body | 1395.08 | 378.001 | 824.165 | 226.586 |
| right dorsal limb | 14974.9 | 3066.23 | 13122.2 | 1058.95 |
| right anterior commissure, olfactory limb | 9035.55 | 1175.95 | 9708.76 | 1072.51 |
| right optic nerve | 2039.68 | 272.325 | 2389.66 | 528.805 |
| right brachium of the superior colliculus | 2854.23 | 733.402 | 2214.03 | 627.433 |
| right superior colliculus commissure | 6466.08 | 1763.27 | 6454.28 | 837.412 |
| right optic chiasm | 1385.45 | 362.514 | 3720.03 | 1517.06 |
| right optic tract | 2098.63 | 421.025 | 1833.56 | 477.664 |
| right oculomotor nerve | 3431.11 | 565.696 | 3635.18 | 490.778 |
| right medial longitudinal fascicle | 1981.21 | 212.845 | 2183.14 | 408.215 |
| right posterior commissure | 7810.29 | 1718.34 | 7890.5 | 1106.5 |
| right trochlear nerve | 4247.79 | 546.352 | 4908.55 | 973.386 |
| right trigeminal nerve | 376.798 | 74.6787 | 608.49 | 92.0082 |
| right motor root of the trigeminal nerve | 1640.59 | 409.585 | 1159.06 | 305.739 |
| right sensory root of the trigeminal nerve | 344.47 | 72.332 | 594.407 | 99.5626 |
| right spinal tract of the trigeminal nerve | 298.823 | 48.3567 | 484.616 | 99.185 |
| right facial nerve | 1918.64 | 481.177 | 1221.52 | 224.141 |
| right genu of the facial nerve | 4784.4 | 1316.12 | 3108.7 | 622.856 |
| right vestibulocochlear nerve | 952.818 | 75.9621 | 1236.64 | 208.564 |
| right vestibular nerve | 380.62 | 89.9998 | 354.003 | 86.6715 |
| right cochlear nerve | 1054.45 | 91.2074 | 1393.4 | 234.771 |
| right trapezoid body | 124.75 | 31.436 | 272.683 | 114.585 |
| right dorsal acoustic stria | 1047.37 | 371.156 | 689.061 | 131.705 |
| right lateral lemniscus | 831.029 | 138.146 | 1593.7 | 296.56 |
| right inferior colliculus commissure | 6263.67 | 1225.53 | 5250.43 | 1174.07 |
| right brachium of the inferior colliculus | 2581.35 | 696.064 | 1844.21 | 341.352 |
| right vagus nerve | 2425.86 | 591.953 | 5636.57 | 1774.9 |
| right solitary tract | 2425.86 | 591.953 | 5636.57 | 1774.9 |
| right dorsal roots | 1840.89 | 372.029 | 1997.69 | 457.951 |
| right cervicothalamic tract | 1840.89 | 372.029 | 1997.69 | 457.951 |
| right dorsal column | 585.091 | 423.135 | 3375.53 | 1315.21 |
| right cuneate fascicle | 585.091 | 423.135 | 3375.53 | 1315.21 |
| right medial lemniscus | 1882.88 | 389.764 | 1951.62 | 495.877 |
| right cerebellum related fiber tracts | 875.718 | 137.597 | 2560.52 | 621.459 |
| right cerebellar commissure | 78.5035 | 30.7716 | 4901.56 | 1754.47 |
| right cerebellar peduncles | 948.518 | 143.442 | 1311.26 | 363.174 |
| right superior cerebelar peduncles | 1253.94 | 173.313 | 1614.71 | 312.805 |
| right superior cerebellar peduncle decussation | 1329.05 | 451.286 | 1897.67 | 730.398 |
| right uncinate fascicle | 506.255 | 111.933 | 1526.61 | 538.792 |
| right ventral spinocerebellar tract | 622.922 | 65.9265 | 1160.69 | 155.19 |
| right middle cerebellar peduncle | 965.338 | 348.075 | 1209.65 | 389.41 |
| right inferior cerebellar peduncle | 624.855 | 155.452 | 1138.4 | 434.866 |
| right dorsal spinocerebellar tract | 1140.1 | 421.704 | 1396.23 | 312.113 |
| right arbor vitae | 853.912 | 147.037 | 3045.85 | 751.241 |
| right supra-callosal cerebral white matter | 14928.1 | 1262.08 | 16078.5 | 1979.99 |
| right lateral forebrain bundle system | 7446.79 | 890.595 | 8607.84 | 444.461 |
| right corpus callosum | 7493.24 | 732.35 | 9322.87 | 572.546 |
| right corpus callosum, anterior forceps | 9653.02 | 1220.27 | 11874.5 | 1087.78 |
| right external capsule | 13553.7 | 1896.57 | 15450.8 | 1544.12 |
| right corpus callosum, extreme capsule | 9825.31 | 579.137 | 9782.53 | 1406.08 |
| right genu of corpus callosum | 5803.52 | 854.213 | 6699.04 | 1171.39 |
| right corpus callosum, posterior forceps | 12726.3 | 1742.54 | 14110.6 | 1244.72 |
| right corpus callosum, body | 4452.7 | 406.402 | 6407.87 | 953.549 |
| right corpus callosum, splenium | 6950.95 | 991.603 | 9447.23 | 1228.65 |
| right corticospinal tract | 865.316 | 124.429 | 1072.65 | 124.269 |
| right internal capsule | 1064.95 | 176.464 | 1296.97 | 157.917 |
| right cerebal peduncle | 650.687 | 151.693 | 914.663 | 238.882 |
| right pyramid | 165.886 | 99.494 | 156.373 | 49.4421 |
| right pyramidal decussation | 2747.26 | 1064.73 | 3938.68 | 1213.96 |
| right thalamus related | 17545.5 | 2979.39 | 17991.3 | 1710.97 |
| right external medullary lamina of the thalamus | 692.329 | 215.364 | 2574.51 | 1324.52 |
| right optic radiation | 19511.3 | 3472.12 | 20132.8 | 2024.83 |
| right auditory radiation | 13180 | 1682.91 | 12430.5 | 1567.05 |
| right extrapyramidal fiber systems | 1441.21 | 198.944 | 1448.79 | 184.439 |
| right cerebral nuclei related | 3735.11 | 581.825 | 4220.5 | 968.991 |
| right nigrostriatal tract | 3735.11 | 581.825 | 4220.5 | 968.991 |
| right tectospinal pathway | 984.206 | 144.658 | 919.976 | 108.34 |
| right direct tectospinal pathway | 0 | 0 | 0 | 0 |
| right doral tegmental decussation | 3789.47 | 896.856 | 3742.69 | 653.856 |
| right crossed tectospinal pathway | 944.815 | 144.547 | 880.336 | 111.979 |
| right rubrospinal tract | 1402.34 | 235.81 | 1381.97 | 200.268 |
| right ventral tegmental decussation | 2258.1 | 326.262 | 3558.48 | 706.935 |
| right medial forebrain bundle system | 7203.04 | 736.667 | 7741.25 | 475.137 |
| right cerebrum related | 7751.79 | 816.521 | 8275.83 | 524.3 |
| right amygdalar capsule | 11663.9 | 1843.91 | 11361 | 1663.27 |
| right anterior commissure, temporal limb | 4256.03 | 1183.16 | 4684.45 | 593.504 |
| right cingulum bundle | 19052.2 | 2170.71 | 19558.4 | 1439.47 |
| right fornix system | 4834.4 | 502.539 | 5394.41 | 456.941 |
| right alveus | 6672.32 | 647.432 | 7756.51 | 563.918 |
| right dorsal fornix | 1359.65 | 379.008 | 3026.32 | 714.98 |
| right fimbria | 1832.02 | 169.298 | 2575.4 | 577.809 |
| right postcommissural fornix | 5303.3 | 986.31 | 5393.39 | 585.981 |
| right medial corticohypothalamic tract | 3458.33 | 2004.7 | 4166.67 | 1344.57 |
| right columns of the fornix | 5358.07 | 1003.89 | 5429.81 | 598.621 |
| right hippocampal commissures | 6778.93 | 1138.85 | 6438.81 | 1172.94 |
| right dorsal hippocampal commissure | 6995.51 | 1206.97 | 6573.68 | 1186.75 |
| right ventral hippocampal commissure | 3681.95 | 617.962 | 4510.13 | 1200.74 |
| right stria terminalis | 2452.9 | 327.259 | 3047.34 | 477.602 |
| right commissural branch of stria terminalis | 4878.33 | 953.095 | 6989.15 | 1613.35 |
| right hypothalamus related | 3044.61 | 300.612 | 3690.19 | 636.438 |
| right medial forebrain bundle | 5660.82 | 1062.7 | 6917.29 | 1522.63 |
| right supraoptic commissures | 707.68 | 314.341 | 1106.55 | 135.765 |
| right mammillary related | 4449.57 | 457.398 | 4391.93 | 938.032 |
| right principal mammillary tract | 8656.72 | 1087.63 | 8129.35 | 2054.74 |
| right mammillothalamic tract | 5136.38 | 658.557 | 4747.65 | 1289.68 |
| right mammillotegmental tract | 2688.22 | 339.238 | 3362.82 | 479.832 |
| right mammillary peduncle | 946.32 | 245.864 | 1203.47 | 332.592 |
| right epithalamus related | 2148.29 | 250.125 | 3120.91 | 622.053 |
| right stria medullaris | 2026.47 | 416.28 | 3363.5 | 886.226 |
| right fasciculus retroflexus | 2350.89 | 311.964 | 3263.35 | 963.189 |
| right habenular commissure | 2129.16 | 454.994 | 1252.45 | 265.319 |
| right ventricular systems | 3301.3 | 302.549 | 3632.66 | 300.465 |
| right lateral ventricle | 4050.53 | 435.067 | 3862.75 | 311.329 |
| right subependymal zone | 5707.45 | 1001.19 | 7731.82 | 1153.29 |
| right choroid plexus | 1252.59 | 152.213 | 1365.48 | 178.462 |
| right third ventricle | 4179.46 | 491.543 | 5127.27 | 508.139 |
| right cerebral aqueduct | 1175.9 | 188.014 | 3486.34 | 638.199 |
| right fourth ventricle | 513.494 | 148.246 | 1381.56 | 375.314 |
| right lateral recess | 381.851 | 87.1691 | 1025.46 | 338.124 |
| right central canal, spinal cord/medulla | 4622.22 | 1458.79 | 3200 | 1306.39 |
